# Enzyme activity as an actionable axis for small-molecule precision oncology

**DOI:** 10.64898/2026.04.14.717586

**Authors:** Kyohhei Fujita, Mako Kamiya, Shingo Dan, Ryo Tachibana, Minoru Kawatani, Ryosuke Kojima, Rumi Hino, Kimihiko Kobayashi, Shotaro Inoue, Miyuki Tani, Yuko Hirata, Shun Kawashima, Kanami Yamazaki, Yumiko Nishimura, Yoshimi Ohashi, Sho Isoyama, Akihiro Nakada, Nobuyoshi Matsumoto, Yuji Ikegaya, Jun Nakajima, Yasuteru Urano

## Abstract

Current precision oncology—molecular targeted therapies and immunotherapies—relies on genomic or expressed biomarkers, yet most cancer patients remain ineligible for these treatments. Here, we establish enzyme activity as an actionable and orthogonal axis for precision cancer medicine. Strategic activity-based screening of mouse organs and human clinical specimens with a panel of enzyme-reactive fluorescence probes identified β-galactosidase 1 (GLB1) and β-hexosaminidases (HEX) as broadly elevated tumor-selective biomarkers. Leveraging these activities, we developed 7-ethyl-10-hydroxycamptothecin (SN38)-based GLB1-and HEX-reactive prodrugs. These prodrugs exhibited dramatically reduced systemic toxicities and improved therapeutic windows, compared to a clinically used SN38-based prodrug, irinotecan (CPT-11). Both prodrugs demonstrated activity-dependent therapeutic efficacy, affording a dramatic reduction of tumor volumes across multiple *in vivo* models, including a subcutaneous patient-derived xenograft (PDX) of lung squamous cell carcinoma that lacked genetic alterations targeted by current precision medicine. Furthermore, this strategy is broadly applicable across various cytotoxic payloads, establishing a generalizable platform for small-molecule precision medicines. Our results define an enzyme-targeting paradigm for precision oncology, in which fluorescence probes serve as companion diagnostic tools to guide development and selection of appropriately targeted prodrugs, which are expected to provide safer and more efficacious treatment options for cancer patients with elevated enzyme activities.

---

Molecular targeted therapies based on genetic diagnostics and cancer immunotherapies based on expression analysis have become mainstream for current precision oncology.^1,2^ Common targets include epidermal growth factor receptor (EGFR), anaplastic lymphoma kinase (ALK), B-type Raf kinase (BRAF) and programmed cell death protein 1 (PD-1).^3,4^ These therapies demonstrate remarkable efficacy in specific patient populations, but more than 80% of cancer patients are ineligible, highlighting the shortage of efficient strategies for developing molecular targeted drugs.^5,6^ In addition, the cost of these innovative cancer therapeutics is high. Accordingly, the majority of patients continue to rely on small-molecule cytotoxic chemotherapeutics such as platinum, topoisomerase inhibitors, taxanes, nucleoside analogs or anthracyclines.^7^ However, these drugs are associated with severe nonspecific toxicities,^8–10^ exacerbated by the difficulty in achieving cancer selectivity. Therefore, there is an urgent need for broadly applicable and less expensive precision cancer treatment strategies, particularly those leveraging affordable small-molecule therapeutics.

Diverse enzyme activities become dysregulated in cancer due to changes in gene expression, post-translational modifications and the intracellular milieu,^11^ and enzyme activities elevated in cancer cells can be valuable therapeutic biomarkers to mediate drug uncaging.^12^ One approach to reduce side-effects is to employ small-molecule prodrugs designed to release active drugs upon reaction with such biomarker enzymes. So far, various prodrugs targeting biomarker enzymes such as cathepsins (CATs), caspase-3, γ-glutamyl transpeptidase (GGT), aminopeptidase N (APN) and β-glucuronidase, have been developed.^12–14^ However, despite tremendous efforts, none has yet been approved by the US Food and Drug Administration (FDA), mainly because of concerns about high background enzyme activities in normal organs causing undesirable non-specific drug release. In addition, the optimal pairing of target enzyme activities and cancer types remains largely unexplored. Therefore, the identification of highly tumor-selective biomarker enzyme activities, tailored to specific cancer types, is critical for developing safer and more efficacious prodrugs.

Since enzyme activities are influenced by post-translational modifications and the intracellular milieu, it is difficult to identify practical biomarker enzymes by gene expression analyses. Therefore, to discover cancer-specific biomarker enzyme activities, we have developed a fluorescence probe library consisting of 800 probes targeting aminopeptidases and glycosidases, some of which are known to exhibit elevated activities in malignant tissues.^15–17^ Unbiased activity-based screening using human surgical specimens has identified efficient cancer imaging probe/biomarker enzyme pairs in various cancers.^15,17–19^ The selected probes were able to sensitively visualize cancer lesions in topical applications.^20^ For example, probes targeting GGT, puromycin sensitive aminopeptidase (PSA) and dipeptidyl peptidase-IV (DPP-IV) have achieved sensitive lung cancer imaging in surgical specimens.^15,16,21^ However, high enzyme activity of these targets in normal organs limits their systemic use, as we previously described in the case of GGT-and DPP-IV-reactive probes.^22,23^ Thus, to discover practical enzyme activities for the development of clinically efficient prodrugs, it is critically important to consider the target enzyme activity levels in systemic organs.

Here, we establish enzyme activity as an actionable and orthogonal axis for precision oncology beyond genomics-and expression-driven approaches, and introduce a strategy for enzyme activity–based patient stratification and selective drug activation in tumors. Through activity-based screening of mouse organs and human clinical specimens, we identify tumor-selective enzymatic activities and develop corresponding small-molecule prodrugs that exhibit activity-dependent efficacy with reduced systemic toxicities *in vivo*, including in tumors lacking actionable genetic alterations targeted by current precision therapies. Importantly, this strategy is broadly applicable across diverse cytotoxic payloads, establishing a generalizable platform for small-molecule precision medicines targeting enzymatic activities.

## Results

Activity-based screening to identify tumor-selective biomarker enzyme activities for prodrug development.

Firstly, we established an activity-based screening strategy, in which mouse organ lysates and clinical cancer specimens are used to directly identify suitable substrate/biomarker enzyme pairs for the development of tumor-selective prodrugs (**Fig. 1a**). To avoid non-specific drug release, enzyme activities in major systemic organs must also be determined. Thus, we initially screened the reactivity of our panel of aminopeptidase-or glycosidase-reactive fluorescence probes, including top-performing probes for cancer imaging identified in our previous studies (**Fig. S1, Table S1**),^15,17^ with lysates derived from mouse major organs (**Extended data Fig. 1a-b**). To estimate the systemic stability of these substrates, we defined the *in vitro* background score (B_org,*vitro*_) as the product of the probe conversion rate (CVR) to fluorophore in each organ and the organ weight (kg) (**Fig. 1b, Extended data Fig. 1a, Table S2**). The *in vitro* total background score (B_tot,*vitro*_) was defined as the sum of the background scores in the examined organs (definitions are summarized in **Table S3**). Although these scores do not account for pharmacokinetic factors, they were considered to provide an approximate estimate of substrate stability following systemic administration. The B_tot,*vitro*_ of glycosidase-reactive probes were much lower than those of aminopeptidase-reactive probes, suggesting that these substrates are more stable after systemic administration (**Fig. 1b**). Since several glycosidase activities are known to be elevated in cancer,^17^ this result suggested that these activities might be utilized to develop safer prodrugs.

**Figure 1.**
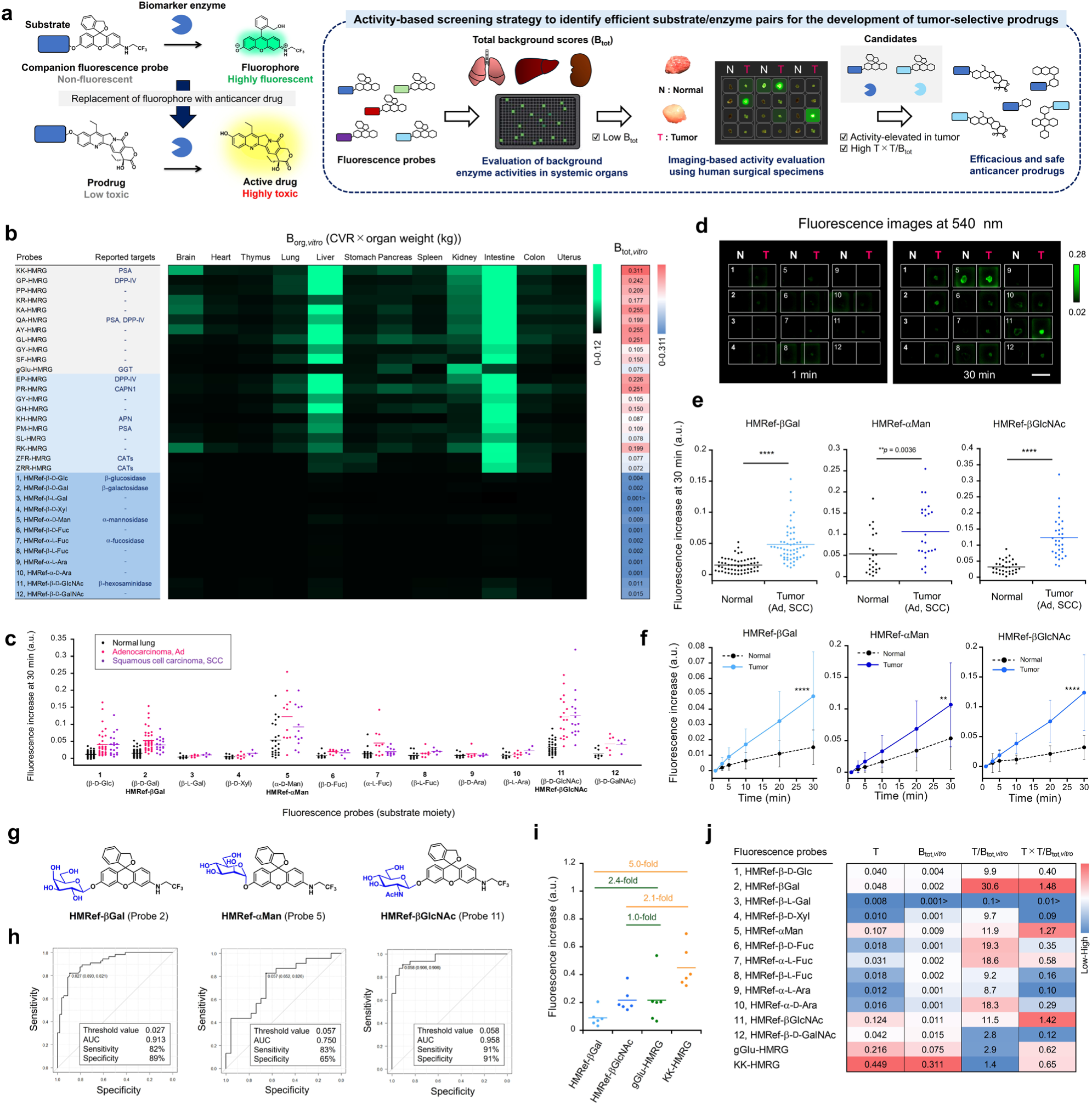
Discovery of substrate/biomarker enzyme pairs for the development of efficient anticancer prodrugs. (**a**) Activity-based screening strategy to identify efficient substrate/biomarker enzyme activity pairs. (**b**) The *in vitro* background scores (B_org,*vitro*_) of evaluated fluorescence probes. The fluorescence probes were incubated with each mouse organ lysate for 1 h. Background scores for each organ were calculated by multiplying the CVR of fluorescence probes by the organ weight (kg). The *in vitro* total background scores (B_tot,*vitro*_) were obtained as the sum of the background scores in all examined organs. [fluorescence probe] = 1 µM, [organ lysate] = 0.1 mg/mL. (**c**) Direct screening of glycosidase-reactive fluorescence probes with human lung cancer specimens. Ad, SCC and normal lung tissues were used (N = 10-53 for normal, N = 6-39 for Ad, N = 4-18 for SCC). Fluorescence increase represents increase at 30 min from 1 min after addition of fluorescence probes. Black, pink, and purple dots represent fluorescence increases in normal lung, Ad and SCC tissues, respectively. (**d**) Representative example of screening of fluorescence probes using surgically resected human lung cancer (SCC) and normal tissues. The probe number of the applied probe is shown on each well. [fluorescence probe] = 50 µM. Scale bar, 1 cm. (**e**) Fluorescence increase of HMRef-βGal and HMRef-βGlcNAc in all evaluated lung cancer tissues (Ad + SCC). \*\*\*\**p*<0.0001 by Welch’s *t*-test. Normal (N = 56), Ad (N = 39) and SCC (N = 17) tissues were examined with HMRef-βGal. Normal (N = 23), Ad (N = 11) and SCC (N = 12) tissues were examined with probe HMRef-αMan. Normal (N = 32), Ad (N = 17) and SCC (N = 15) tissues were examined with probe HMRef-βGlcNAc. The line in the diagram represents the average. (**f**) Time-dependent fluorescence increase of HMRef-βGal, HMRef-αMan and HMRef-βGlcNAc in lung normal and cancer (Ad and SCC) tissues from (**e**). Black lines represent fluorescence increases in normal lung tissues. Light blue, deep blue or blue lines represent fluorescence increases in lung cancer (Ad and SCC) tissues. Error bars represent s.d. \*\*\*\**p*<0.0001, \*\**p*<0.01 by Welch’s *t*-test. (**g**) Chemical structures of HMRef-βGal, HMRef-αMan and HMRef-βGlcNAc. (**h**) ROC curves for lung cancer detection. Threshold value, sensitivity, specificity and AUC of each probe were evaluated from the ROC curve. Normal (N = 56), Ad (N = 39) and SCC (N = 17) tissues were examined with HMRef-βGal. Normal (N = 23), Ad (N = 11) and SCC (N = 12) tissues were examined with probe HMRef-αMan. Normal (N = 32), Ad (N = 17) and SCC (N = 15) tissues were examined with probe HMRef-βGlcNAc. (**i**) Comparison of fluorescence increases of HMRef-βGal, HMRef-βGlcNAc, gGlu-HMRG and KK-HMRG. Fluorescence increase represents increase at 30 min from 1 min after addition of fluorescence probes. The line in the diagram represents the average. (**j**) Selection of suitable substrates for the development of tumor-selective prodrugs based on T × T/B_tot,*vitro*_ scores. T values represent average fluorescence increase in clinical lung cancer specimens. B_tot,*vitro*_ scores represent *in vitro* total background scores from (**b**).

Next, we evaluated whether these glycosidase activities are elevated in cancer tissues. Here, we focused on non-small cell lung cancer (NSCLC), the leading cause of cancer-related mortality, in which approximately 40–60% of patients are ineligible for current precision medicine. We screened our 12 glycosidase-reactive fluorescence probes in clinical lung cancer specimens by *ex vivo* fluorescence imaging. Specimens of normal lung tissue and NSCLC (adenocarcinoma (Ad) or squamous cell carcinoma (SCC)), from the same patient were divided into pieces and incubated with each probe to evaluate the corresponding glycosidase activities (**Fig. 1c-d, Extended data Fig. 1c, Fig. S2**). Notably, probe 1; HMRef-β-D-Glc (HMRef-βGlc), probe 2; HMRef-β-D-Gal (HMRef-βGal), probe 5; HMRef-α-D-Man (HMRef-αMan), probe 7; HMRef-α-L-Fuc (HMRef-αFuc); and probe 11; HMRef-β-D-GlcNAc (HMRef-βGlcNAc) showed markedly greater fluorescence increases in lung Ad and SCC tissues than in normal lung tissues (**Fig. 1c-g**). The sensitivity and specificity for cancer detection were calculated to be 60% and 93% (AUC = 0.841) for HMRef-βGlc, 82% and 89% (AUC = 0.913) for HMRef-βGal, 83% and 65% (AUC = 0.750) for HMRef-αMan, 91% and 46% (AUC = 0.787) for HMRef-αFuc, 91% and 91% (AUC = 0.958) for HMRef-βGlcNAc, respectively, indicating that the target enzyme activities of these probes are broadly elevated in NSCLC patients (**Table 1 and Fig. 1h, Fig. S3**). These probes showed fluorescence increases that are equivalent to or only a few-fold lower than those of previously reported lung cancer imaging probes, GGT-reactive gGlu-HMRG and PSA-reactive KK-HMRG,^15,21^ despite exhibiting extremely low B_tot,*vitro*_ (**Fig. 1i, b**). For prodrug development, high activation in tumors combined with low background activation in systemic organs is desirable. To identify such substrates, we considered two parameters: fluorescence increase in tumors (T) and tumor/background (T/B_tot,*vitro*_) ratio. To take account of both parameters, we calculated the T × T/B_tot,*vitro*_ score of each probe. Among the evaluated probes, HMRef-βGal, HMRef-αMan and HMRef-βGlcNAc especially exhibited high T × T/B_tot,*vitro*_ scores, suggesting high selectivity for tumors (**Fig. 1j**). These scores are higher than those of gGlu-HMRG and KK-HMRG. In addition, HMRef-βGal, HMRef-αMan and HMRef-βGlcNAc are significantly more stable in human serum than fluorescence probes targeting previously reported target enzymes for lung cancer imaging: gGlu-HMRG, KK-HMRG, CATs-reactive Z-Phe-Arg-HMRG (ZFR-HMRG) and DPP-IV-reactive EP-HMRG (**Extended data Fig. 1d**).^24,25^ Since HMRef-αMan showed increased fluorescence in normal lung tissues of some patients, suggesting a potential risk of pulmonary toxicity (**Fig. 1e**), we selected HMRef-βGal and HMRef-βGlcNAc for further elaboration to construct prodrugs for lung cancer treatment.

**Table 1.** Performance of fluorescence probes for the detection of lung cancer. Threshold value, sensitivity and specificity of each probe were evaluated from the receiver operating characteristic curve. AUC, Area under the curve. Values for gGlu-HMRG and KK-HMRG were taken from the literature.^15,21^

| Probes | Target enzymes | Lung cancer types | Number of patients | Threshold values | Sensitivity | Specificity | AUC |
| --- | --- | --- | --- | --- | --- | --- | --- |
| 1. HMRef-β-D-Glc | β-glucosidases | Ad | 35 | 0.011 | 89% | 60% | 0.860 |
|  |  | SCC | 18 | 0.026 | 61% | 94% | 0.802 |
|  |  | Ad + SCC | 53 | 0.024 | 60% | 93% | 0.841 |
| 2. HMRef-βGal | β-galactosidases | Ad | 39 | 0.024 | 92% | 77% | 0.917 |
|  |  | SCC | 17 | 0.027 | 77% | 100% | 0.900 |
|  |  | Ad + SCC | 56 | 0.027 | 82% | 89% | 0.913 |
| 5. HMRef-αMan | α-mannosidases | Ad | 11 | 0.060 | 91% | 73% | 0.810 |
|  |  | SCC | 12 | 0.044 | 83% | 58% | 0.701 |
|  |  | Ad + SCC | 23 | 0.057 | 83% | 65% | 0.750 |
| 7. HMRef-α-L-Fuc | α-fucosidases | Ad | 10 | 0.011 | 80% | 70% | 0.676 |
|  |  | SCC | 12 | 0.010 | 92% | 42% | 0.736 |
|  |  | Ad + SCC | 22 | 0.008 | 91% | 46% | 0.787 |
| 11. HMRef-βGlcNAc | β-hexosaminidases | Ad | 17 | 0.061 | 82% | 94% | 0.969 |
|  |  | SCC | 15 | 0.069 | 93% | 93% | 0.951 |
|  |  | Ad + SCC | 32 | 0.058 | 91% | 91% | 0.958 |
| gGlu-HMRG | GGT | Ad | 19 | - | 79% | 74% | 0.795 |
|  |  | Various lung cancer | 73 | - | 44% | 85% | 0.623 |
| KK-HMRG | PSA | Ad | 19 | - | - | - | - |
|  |  | SCC | 13 | - | - | - | - |
|  |  | Ad + SCC | 32 | - | 85% | 73% | 0.842 |

### Evaluation of systemic selectivity of the target enzymes of HMRef-βGal and HMRef-βGlcNAc *in vivo*

To further examine *in vivo* tumor selectivity of the target enzymes of HMRef-βGal and HMRef-βGlcNAc, we intravenously administered the probes to mice and evaluated the background enzyme activity of various organs. We also compared the background activation of probes for reported enzyme targets such as GGT, PSA and CATs. Twenty min post-injection of 5 mg/kg HMRef-βGal, HMRef-βGlcNAc, gGlu-HMRG, KK-HMRG or ZFR-HMRG to female ICR mice, major organs were resected and subjected to fluorescence imaging (**Fig. 2a, Fig. S4**). HMRef-βGal and HMRef-βGlcNAc showed only minimal fluorescence in all examined organs, whereas gGlu-HMRG, KK-HMRG and ZFR-HMRG exhibited significant fluorescence signals in several organs, indicating that GGT, PSA and CATs activities are abundant in some normal tissues, resulting in substantial background activation (**Fig. 2b**). To quantitatively assess these results, the product of the fluorescence intensity and organ weight (kg) was calculated for each organ (B_org,*vivo*_), and the sum of these values was defined as the *in vivo* total background score (B_tot,*vivo*_) (**Table S3**). HMRef-βGal and HMRef-βGlcNAc exhibited markedly lower B_tot,*vivo*_ scores than probes targeting reported target enzymes, suggesting that the enzyme activities hydrolyzing these probes are extremely low in normal tissues (**Fig. 2c**). Although these results do not take account of pharmacokinetic factors, positive trends were observed between B_org,*vitro*_ and B_org,*vivo*_ scores, suggesting that B_tot,*vitro*_ can be used to predict background enzyme activities *in vivo* (**Extended data Fig. 2a-b**). Moreover, using T values from lung cancer specimens, HMRef-βGal and HMRef-βGlcNAc showed high values of T/B_tot,*vivo*_, consistent with the activity-based screening, suggesting high tumor-selectivity (**Fig. 2c**).

**Figure 2.**
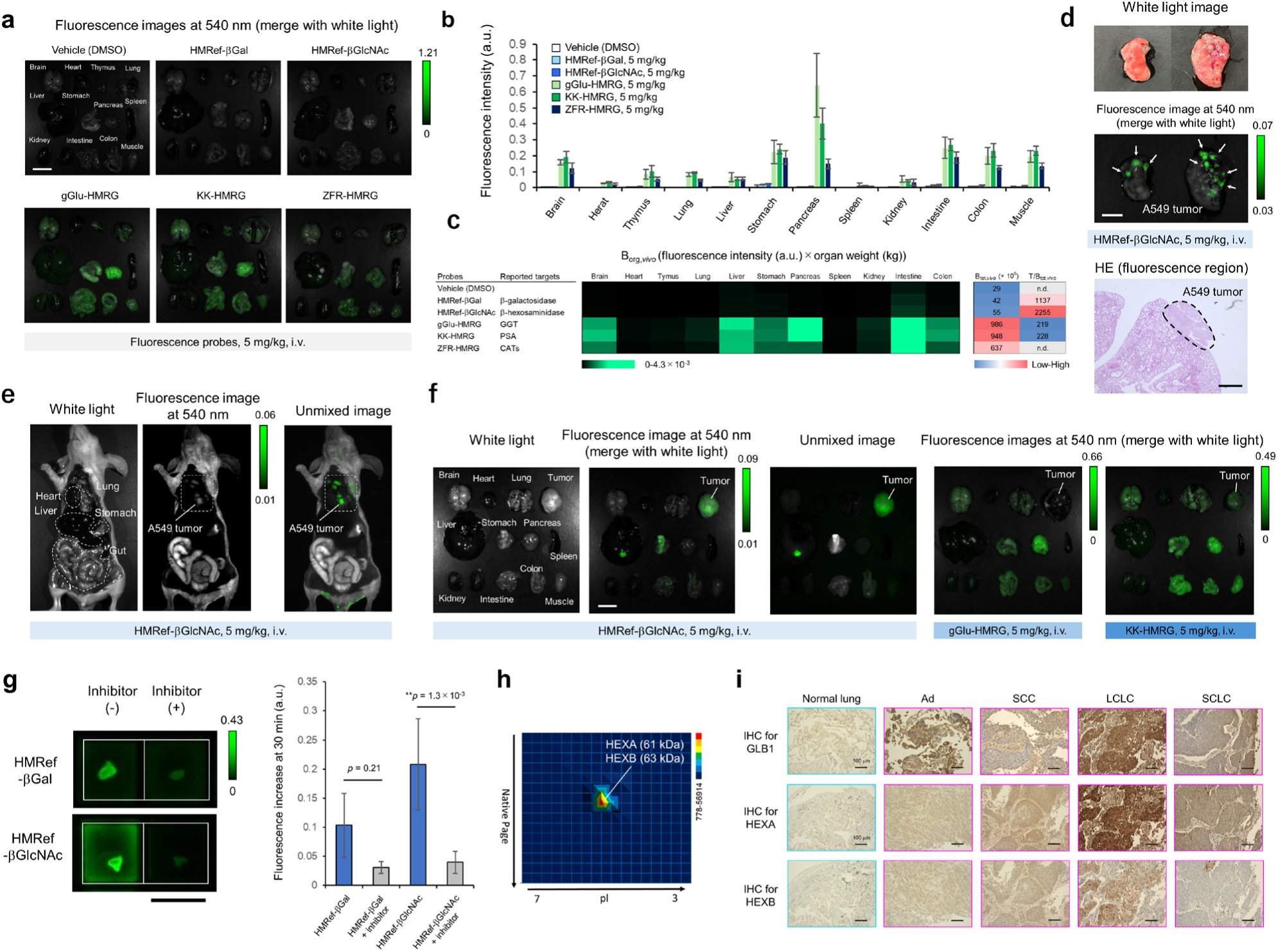
Evaluation of *in vivo* selectivity of target enzyme activities and their identification. (**a**) Fluorescence images of freshly resected organs 20 min after intravenous administration of 5 mg/kg fluorescence probe (HMRef-βGal, HMRef-βGlcNAc, gGlu-HMRG, KK-HMRG or ZFR-HMRG) to female ICR mice (n = 3). Ex/Em = 465 nm/515 nm long pass. Scale bar, 1 cm. (**b**) Fluorescence intensities of organs in (**a**) (n = 3). Error bars represent s.d. (**c**) *In vivo* background scores of each organ (B_org,*vivo*_), calculated by multiplying the mean fluorescence intensity by the corresponding organ weight (kg). *In vivo* total background score (B_tot,*vivo*_) is the sum of the background scores in all organs. T values in **Fig. 1** (**j**) were used to calculate T/B_tot,*vivo*_ scores. n.d. = no data. (**d**) *Ex vivo* fluorescence imaging of freshly resected lung 30 min after intravenous administration of 5 mg/kg HMRef-βGlcNAc to lung orthotopic A549 tumor-bearing mouse (upper). Scale bar, 5 mm. Ex/Em = 465 nm/515 nm long pass. Fluorescent regions were well matched with pathologically identified cancer regions (bottom). Scale bar, 1 mm. (**e**) Fluorescence imaging of lung cancer regions 30 min after intravenous administration of 5 mg/kg HMRef-βGlcNAc to lung orthotopic A549 tumor-bearing mouse. Selective fluorescence activation in cancer regions is seen in the unmixed fluorescence image (green, HMRef signal; white, autofluorescence signal). Ex/Em = 465 nm/515 nm long pass. (**f**) *Ex vivo* fluorescence imaging of freshly resected tumor and each organ 20 min after intravenous administration of 5 mg/kg HMRef-βGlcNAc, gGlu-HMRG or KK-HMRG to a subcutaneous lung cancer PDX model. PDX tumor was selectively visualized in the unmixed fluorescence image (green, HMRef signal; white, autofluorescence signal) after administration of HMRef-βGlcNAc. Ex/Em = 465 nm/515 nm long pass. Scale bar, 1 cm. (**g**) Fluorescence images of lung cancer tissues without (left) and with inhibitor (right) at 30 min. Blue bars: fluorescence increase in the absence of inhibitors. Gray bars: fluorescence increase in the presence of inhibitors. Fluorescence increase represent increase at 30 min from 1 min after addition of fluorescence probes. One-way ANOVA with Tukey’s multiple comparisons test was used to calculate exact *p* values. Ex/Em = 465 nm/515 nm long pass. [Fluorescent probe] = 50 µM, [N-(n-nonyl)deoxygalactonoijirimycin] = 500 µM for HMRef-βGal, [PUGNAc] = 500 µM for HMRef-βGlcNAc. Error bars represent s.d. Scale bar, 1 cm. (**h**) DEG assay for Ad tissue using HMRef-βGlcNAc. HEXA (61 kDa) and HEXB (63 kDa) were identified by peptide mass fingerprinting analysis from the fluorescent spot on 2D gel. (**i**) IHC staining for GLB1, HEXA and HEXB in normal lung, Ad, SCC, LCLC and SCLC tissues. These enzymes are overexpressed in lung cancer. Scale bars, 100 µm.

To further evaluate the tumor-selectivity, HMRef-βGlcNAc, which exhibited the highest T/B_tot,*vivo*_ score and the highest sensitivity and specificity in the screening, was intravenously administered to orthotopic lung tumor-bearing mice. We used lung Ad-derived A549 tumor since it can highly activate the probe (**Extended data Fig. 2c-e, Fig. S5**). In the unmixed fluorescence images, where the background autofluorescence was eliminated, the probe was able to selectively visualize cancer lesions with low background signals in normal organs (**Fig. 2d-e, Extended data Fig. 2f-g**). We also intravenously administered HMRef-βGlcNAc, HMRef-βGal, gGlu-HMRG or KK-HMRG to mice bearing a subcutaneously implanted lung SCC patient-derived xenograft (PDX), and performed fluorescence imaging. In the unmixed image, HMRef-βGlcNAc especially exhibited highly selective activation in tumor tissues (**Fig. 2f**), suggesting that HMRef-βGlcNAc is more strongly activated than HMRef-βGal at tumor sites (**Extended data Fig. 2h**). In contrast, probes targeting GGT and PSA exhibited significant background fluorescence in normal organs and were unable to selectively visualize tumors (**Fig. 2f, Extended data Fig. 2i-j**), as anticipated based on their T/B_tot,*vitro*_ scores (**Fig. 1j**). These findings suggested that the target enzymes of HMRef-βGlcNAc and HMRef-βGal exhibit lower activities in normal organs and greater lung tumor selectivity than previously reported biomarker enzymes.

### Identification of the target enzymes of HMRef-βGal and HMRef-βGlcNAc

In order to confirm the target enzymes of HMRef-βGal and HMRef-βGlcNAc, we evaluated the fluorescence increase of these probes in lung cancer specimens with or without enzyme inhibitors. We found that the fluorescence increases of HMRef-βGal and HMRef-βGlcNAc were significantly suppressed by the β-galactosidase inhibitor N-(n-nonyl)deoxygalactonoijirimycin and the β-hexosaminidase (HEX) inhibitor PUGNAc, respectively, strongly indicating that the activation of these probes was induced by these glycosidase activities (**Fig. 2g**). Similar results were obtained in A549 (Ad) and NCI-H226 (SCC) lung cancer cell lines (**Extended data Fig. 2c-e**). We considered that the target of HMRef-βGal would be β-galactosidase 1 (GLB1),^26^ since this isozyme is a major human lysosomal β-galactosidase, and we confirmed that the probe is activated by GLB1 (**Extended data Fig. 2k**). In the case of HMRef-βGlcNAc, we assumed the target enzyme would be HEX. Since there are several isozymes of HEX,^27–29^ we set out to identify the responsible subtype by diced electrophoresis gel (DEG) assay, which employs a combination of 2D-gel fluorometric assay and peptide mass fingerprinting,^30^ using HMRef-βGlcNAc and lysate from an Ad specimen. We observed a single fluorescent spot on the gel, and identified β-hexosaminidase subunit α (HEXA) and subunit β (HEXB) encoded by *HEXA* and *HEXB* genes by peptide mass fingerprinting analysis, suggesting that β-hexosaminidase A (HexA, heterodimer of HEXA and HEXB), β-hexosaminidase B (HexB, homodimer of HEXB), and β-hexosaminidase S (HexS, homodimer of HEXA) activities are associated with the fluorescence increase (**Fig. 2h, Fig. S6, Table. S4**). We also evaluated the reactivity of HMRef-βGlcNAc with recombinant HEXA and HEXB, and confirmed that the probe reacts with them (**Extended data Fig. 2k**). Finally, to confirm that GLB1 and HEX are target enzymes of these probes, we evaluated the expression levels of GLB1, HEXA and HEXB in surgical specimens by IHC staining. Normal lung tissues showed little target staining, whereas Ad and SCC tissues showed stronger staining (**Fig. 2i**, **Fig. S7-8**), consistent with the involvement of GLB1 and HEX as the responsible enzymes. Furthermore, these enzymes also tend to be overexpressed in large cell lung cancer (LCLC) and small cell lung cancer (SCLC) (**Fig. 2i**), suggesting that they may be promising targets in various lung cancer types.

### Development of SN38-based anticancer prodrugs targeting the discovered biomarker enzymes

Having identified GLB1 and HEX, we next applied the discovered substrate/enzyme pairs to the development of prodrugs for lung cancer therapy. To design the prodrugs, we replaced the fluorophore of the hit probes with a cytotoxic anticancer drug (**Fig. 1a**). As an initial proof-of-concept payload, 7-ethyl-10-hydroxycamptothecin (SN38) was selected because it exhibits potent anticancer activity against diverse cancers. Since masking of the hydroxyl group at the 7-position of SN38 is known to suppress its toxicity,^31^ we introduced each substrate sugar at this position. In clinical settings, CPT-11 is widely used as an SN38-based prodrug for treating various cancers including NSCLC,^32–34^ but its therapeutic potential is limited by nonspecific toxicities, including severe gut damage, caused by carboxylesterases (CES)-mediated drug release in liver and gut.^35,36^ In contrast, the prodrugs designed in this study are expected to produce active SN38 selectively in cancer sites after one-step cleavage of their substrate sugar in the presence of the target enzyme (**Fig. 3a**). Based on this design concept, we synthesized SN38-βGal and SN38-βGlcNAc (**Fig. 3b**). We confirmed prodrug reactivities with GLB1, HEXA and HEXB proteins by LC-MS analysis (**Fig. S9**). We next conducted MTT assays to examine the *in vitro* cell toxicity of the developed prodrugs towards NCI-H226 lung SCC cells, SHP-77 lung SCLC cells and HEK/lacZ cells, which exhibited different levels of β-galactosidase and HEX activities in fluorescence imaging (**Fig. 3c**, **Fig. S10**). In NCI-H226 cells, SN38-βGal and SN38-βGlcNAc showed comparable IC_50_ values. In contrast, in SHP-77 cells, which exhibit strong HEX activities but low GLB1 activity, SN38-βGlcNAc exhibited a significantly lower IC_50_ than SN38-βGal. Conversely, in HEK/lacZ cells with high expression of β-galactosidase (lacZ), SN38-βGal showed a significantly lower IC_50_ (**Fig. 3d**, **Extended data Fig. 3a**). These results correlated well with both the fluorescence intensity of the probes and the amount of SN38 produced (**Fig. 3e**). In SHP-77 cells, production of SN38 from SN38-βGal was negligible, and the log IC_50_ showed an approximately 2-log shift compared to that of SN38. These results indicate that the developed prodrugs are activated in an enzyme activity-dependent manner. To further support this conclusion, we evaluated time-dependent changes in the numbers of live A549 lung Ad cells in the presence and absence of target enzyme inhibitors by means of time-lapse imaging. Live-cell nuclei were stained with SPY650-DNA. Both SN38-βGal and SN38-βGlcNAc caused concentration-dependent cell growth inhibition, while their efficacy was significantly suppressed in the presence of the corresponding glycosidase inhibitor (**Fig. 3f**, **Extended data Fig. 3b, Fig. S11-13**, **Supplementary Video 1**). Indeed, treatment with these prodrugs induced S-phase arrest with nuclear enlargement, consistent with replication stress, whereas co-treatment with the corresponding glycosidase inhibitor suppressed this effect and allowed cell proliferation (**Fig. 3g, Extended data Fig. 3c**). These results confirm that the developed prodrugs suppress cancer cell growth in an enzyme-activity-dependent manner.

**Figure 3.**
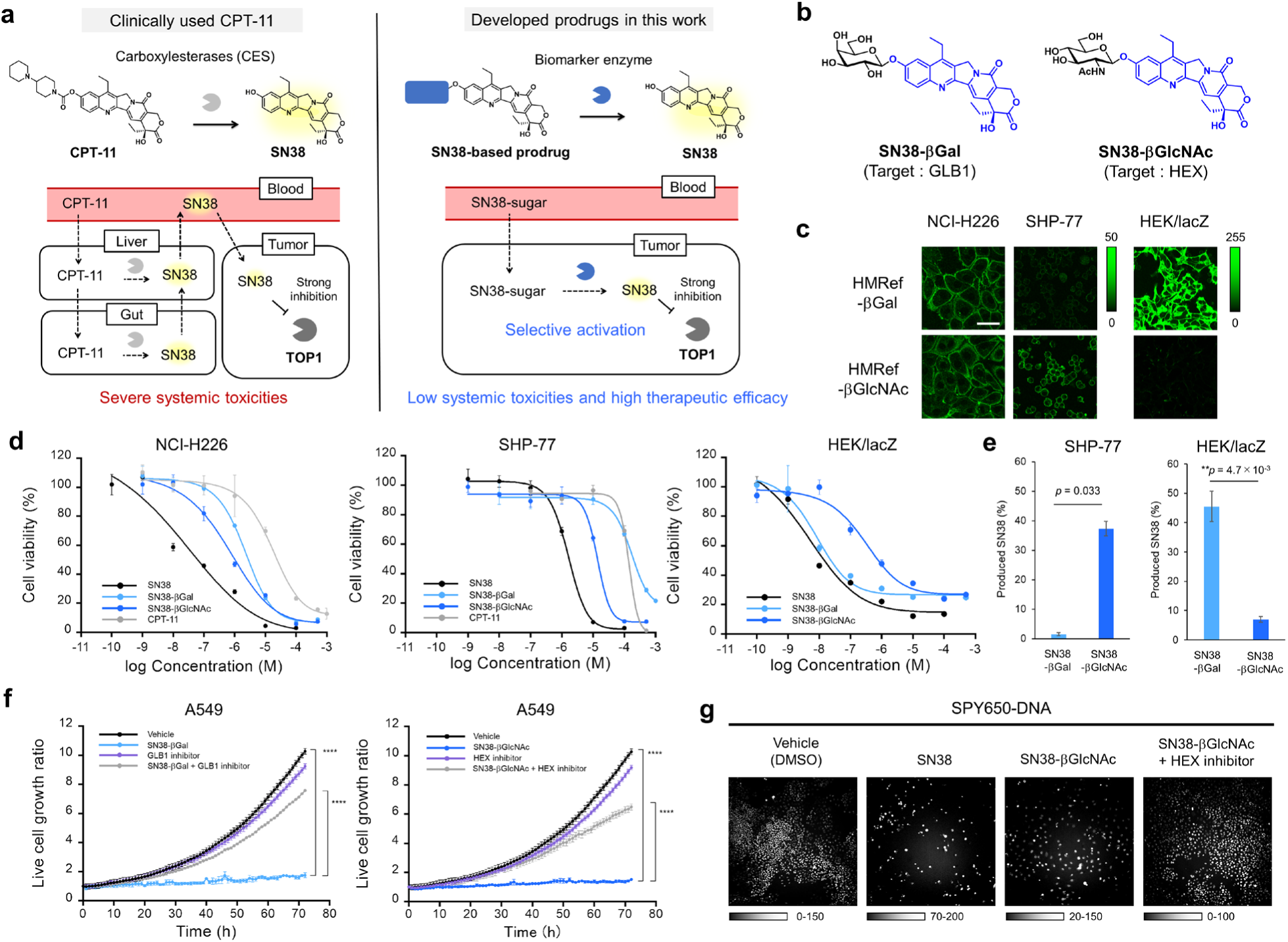
Development of SN38-based anticancer prodrugs targeting identified biomarker enzyme activities. (**a**) The design concept of the developed prodrugs in this study. CPT-11 is activated in liver and gut by CES, whereas the developed prodrugs are expected to be selectively activated in tumor sites. (**b**) Chemical structures of the developed anticancer prodrugs, SN38-βGal and SN38-βGlcNAc. (**c**) Live cell fluorescence imaging of β-galactosidase and β-hexosaminidase activities using HMRef-βGal and HMRef-βGlcNAc in NCI-H226, SHP-77 and HEK/lacZ cells. Fluorescence images were captured 1 h after addition of fluorescence probes. [Fluorescence probes] = 20 µM. (**d**) MTT assays of SN38-βGal and SN38-βGlcNAc for NCI-H226, SHP-77 and HEK/lacZ cells. Error bars represent s.d. (**e**) Production rates of SN38 from SN38-βGal or SN38-βGlcNAc in SHP-77 and HEK/lacZ cells. Production rates were measured by LC-MS 24 h after incubation with each prodrug. Welch’s *t*-tests were used to calculate exact *p* values. Error bars represent s.d. (**f**) Time-dependent growth ratio of A549 cells in the presence and absence of SN38-βGal or SN38-βGlcNAc with or without each inhibitor. Cell growth was significantly suppressed by the prodrugs, and the inhibition was significantly suppressed in the presence of the corresponding enzyme inhibitor. \*\*\*\**p*<0.0001 by one-way ANOVA with Tukey’s multiple comparisons test. Error bars represent s.d. [SN38-βGal] = 10 µM, [SN38-βGlcNAc] = 1 µM, [N-(n-nonyl)deoxygalactonoijirimycin] = 20 µM for SN38-βGal, [PUGNAc] = 20 µM for SN38-βGlcNAc. (**g**) Fluorescence images of A549 cells 72 h after addition of each drug. Nuclei were stained with SPY650-DNA. [SN38] = 1 µM, [SN38-βGlcNAc] = 1 µM, [PUGNAc] = 20 µM for SN38-βGlcNAc. Scale bar, 200 µm. Treatment with SN38-βGlcNAc induced S-phase arrest, whereas co-treatment with the HEX inhibitor PUGNAc suppressed this effect and allowed cell proliferation.

### Evaluation of SN38-βGal and SN38-βGlcNAc in the JFCR39 cancer cell panel

To evaluate pan-cancer efficacy of SN38-βGal and SN38-βGlcNAc, we performed growth-inhibition analysis using the JFCR39 cancer cell panel, which comprises 30 cell lines in NCI60 together with 6 stomach cancer and 3 breast cancer cell lines.^37–39^ Fluorescence-based enzyme activity profiling revealed that GLB1 and HEX activities were observed across diverse cancer cell lines beyond lung cancer, and both prodrugs inhibited proliferation across various tumor types (**Extended data Fig. 3d, Fig. S14**). The enzyme activity levels and the drug efficacy (-log GI_50_(SN38/prodrug)) also tended to correlate with the amount of SN38 produced, supporting the enhanced efficacy in enzyme activity–positive cancer cells (**Fig. S15**). Notably, HEX activities were generally higher than GLB1 activity in cell lines, consistent with the superior efficacy of SN38-βGlcNAc. In addition, SN38-βGlcNAc generally exhibited lower GI₅₀ values than CPT-11, probably due to the markedly higher HEX activities in cancer cells compared with CES activities (**Extended data Fig. 3b-d, Fig. S11-12**).

Importantly, the developed prodrugs exhibited marked growth-inhibitory effects in some cell lines that were not highly responsive to cytotoxic chemotherapeutic drugs (cisplatin, carboplatin, oxaliplatin, gemcitabine, paclitaxel and docetaxel) or to molecular targeted agents (gefitinib, alectinib, crizotinib, entrectinib and dabrafenib) used in current NSCLC treatment (**Extended data Fig. 3d**).^38,39^ These findings indicated that the developed prodrugs may be effective against at least some tumors resistant to platinum-based drugs, nucleoside analogs, taxanes or current molecular targeted therapies.

### Repeated-dose toxicity studies of SN38-βGal and SN38-βGlcNAc

To evaluate the safety of the developed prodrugs, we initially conducted toxicology studies in ICR mice (dosing schedules for all *in vivo* experiments are provided in **Extended Data Table. 1**). We determined the maximum tolerated dose (MTD) for a one-week regimen of three times/week intravenous injection of prodrugs, and confirmed that the administration of SN38-βGal at doses up to 120 mg/kg and SN38-βGlcNAc at doses up to 200 mg/kg did not affect body weight (**Extended Data Fig. 4a-b**). No notable abnormalities were observed in biochemical test values, organ weights or pathological analysis at these doses, suggesting that repeated administration of these prodrugs might be feasible even at high doses of 120-200 mg/kg (**Extended Data Fig. 4c-d, Table. S5**).

Next, we compared the safety of SN38-βGal and SN38-βGlcNAc with that of CPT-11. Healthy female ICR mice were given intravenous administration of SN38-βGal, SN38-βGlcNAc or CPT-11 [40 mg/kg 3 times/week for 3 weeks]. The CPT-11 group exhibited a significant decrease of approximately 10% in body weight, whereas the SN38-βGal and SN38-βGlcNAc groups showed no significant change (**Fig. 4a**). Furthermore, histopathological evaluation revealed severe organ damage to the intestine and colon in the CPT-11 group, whereas no notable organ damage was observed in the SN38-βGal and SN38-βGlcNAc groups (**Fig. 4b**, **Extended Data Fig. 4e**). In addition, no significant changes in biochemical tests and organ weights were observed in the SN38-βGal and SN38-βGlcNAc groups, whereas CPT-11 caused increased spleen weight and decreased kidney weight (**Table. S6**, **Extended Data Fig. 4f**). These results suggested that the developed prodrugs show markedly lower systemic toxicities compared to CPT-11.

**Figure 4.**
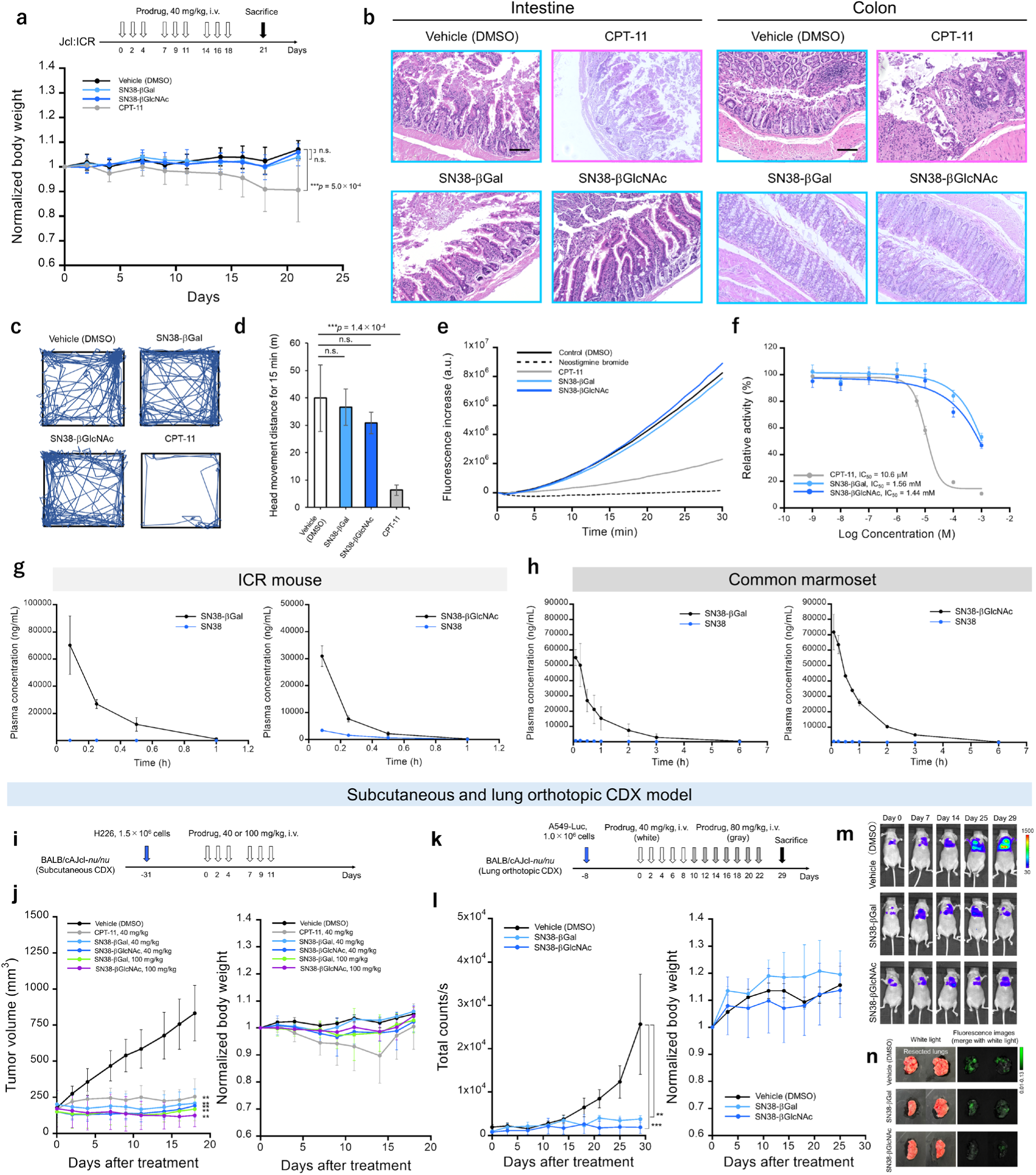
Toxicity, pharmacokinetic and efficacy studies of SN38-βGal and SN38-βGlcNAc. (**a**) Repeated-dose toxicity evaluation of CPT-11, SN38-βGal and SN38-βGlcNAc (n = 8 for each group). One-way ANOVA with Tukey’s multiple comparisons test was used to calculate exact *p* values. n.s. = not significant. Error bars represent s.d. (**b**) Pathological analysis of intestine and colon resected after the repeated-dose toxicity evaluation. Significant intestine and colon damage was observed in CPT-11 group, whereas SN38-βGal and SN38-βGlcNAc did not show notable damage. Scale bars, 100 µm. (**c**) Evaluation of the head movement trajectory in mice during 15 min following intravenous injection of each drug (40 mg/kg). A significant decrease in the locomotion was observed following the administration of CPT-11, whereas SN38-βGal and SN38-βGlcNAc did not cause a decrease. (**d**) Evaluation of head movement distance in mice during the 15 min following intravenous injection of each drug at 40 mg/kg (n = 4 for each group). The significance of differences was determined by one-way ANOVA with Tukey’s HSD (\*\*\**p* < 0.001). n.s. = not significant. Error bars represent s.d. (**e**) Evaluation of AChE-inhibitory activity of CPT-11, SN38-βGal and SN38-βGlcNAc. Neostigmine bromide was used as positive control. CPT-11 significantly suppressed AChE activity. (**f**) Evaluation of the IC_50_ values for AChE inhibition by CPT-11, SN38-βGal and SN38-βGlcNAc. Error bars represent s.d. (**g**) Prodrugs and released SN38 plasma concentrations over a 1 h period after a single prodrug infusion of female ICR mice at a dose of 40 mg/kg (n = 3 for each time point). Error bars represent s.d. (**h**) Prodrug and released SN38 plasma concentrations over 6 h after a single 11-min prodrug infusion of a female common marmoset at a dose of 10 mg/kg (n = 3). Error bars represent s.d. (**i**) Representation of the prodrug treatment in subcutaneous NCI-H226 tumor-bearing mice. (**j**) Anti-tumor efficacy (left) and weight change (right) in subcutaneous NCI-H226 tumor-bearing mice following treatment with vehicle (n = 7), SN38-βGal (40 mg/kg × 6, n = 4), SN38-βGlcNAc (40 mg/kg × 6, n = 5), CPT-11 (40 mg/kg × 6, n = 8), SN38-βGal (100 mg/kg × 6, n = 4) and SN38-βGlcNAc (40 mg/kg × 6, n = 4). \*\**p*<0.01 vs. vehicle by one-way ANOVA with Tukey’s multiple comparisons test. Error bars represent s.d. (**k**) Representation of the prodrug treatment in lung orthotopic A549-Luc tumor-bearing mice. (**l**) Anti-tumor efficacy (left) and weight change (right) in lung orthotopic A549-Luc tumor-bearing mice following treatment with vehicle (n = 3), SN38-βGal (40 mg/kg × 5 followed by 80 mg/kg × 7, n = 4) and SN38-βGlcNAc (40 mg/kg × 5 followed by 80 mg/kg × 7, n = 5). Tumor growth inhibition was quantified using a luciferin–luciferase bioluminescence assay. \*\*\**p*<0.001 vs. vehicle, \*\**p*<0.01 vs. vehicle by one-way ANOVA with Tukey’s multiple comparisons test. Error bars represent s.d. (**m**) Luciferin–luciferase bioluminescence images of treated mice from (**l**). (**n**) Fluorescence imaging of lung orthotopic A549-Luc tumor after treatment with SN38-βGal and SN38-βGlcNAc in (**l**). Tumors were visualized by intravenous administration of 5 mg/kg HMRef-βGlcNAc. Ex/Em = 465 nm/515 nm long pass.

### Comparison of the acute toxicities of SN38-βGal, SN38-βGlcNAc and CPT-11

CPT-11 is known to inhibit AChE and to cause cholinergic syndrome as an acute toxicity.^40^ Indeed, the systemic administration of 40 mg/kg of CPT-11 significantly decreased the locomotion of mice, which can be attributed to cholinergic syndrome. However, the systemic administration of 40 mg/kg of SN38-βGal and SN38-βGlcNAc did not significantly affect the locomotion (**Fig. 4c-d**, **Extended Data Fig. 4g, Fig. S16, Supplementary Video 2**). The LD₅₀ values were determined to be over 300 mg/kg for SN38-βGal and over 500 mg/kg for SN38-βGlcNAc, which are markedly higher than the reported value of approximately 130 mg/kg for CPT-11 (**Table S7**). These results suggested that the developed prodrugs may be less likely to cause cholinergic syndrome. To confirm this, we evaluated the AChE-inhibitory activities of SN38-βGal, SN38-βGlcNAc and CPT-11. The IC₅₀ values for AChE were 1.56 mM for SN38-βGal and 1.44 mM for SN38-βGlcNAc, which are more than 100-fold higher than that of CPT-11 (10.6 µM) (**Fig. 4e-f**). Quantum mechanics/molecular mechanics (QM/MM) calculations suggested that CPT-11 has strong hydrophobic interactions with the AChE hydrophobic pocket composed of Y133, F297, Y337 and F338, whereas the hydrophilic sugar moieties in the developed prodrugs reduce this affinity (**Extended Data Fig. 5a-c**). Consistent with this, the binding energy of CPT-11 was more than 10 kcal/mol higher than that of the developed prodrugs. These results support the view that SN38-βGal and SN38-βGlcNAc show greatly reduced acute toxicity compared to CPT-11.

### Pharmacological analysis of SN38-βGal and SN38-βGlcNAc

The half-lives of SN38-βGal and SN38-βGlcNAc in ICR mice were 0.165 h and 0.149 h, respectively, after a single intravenous administration of 40 mg/kg (**Fig. 4g, Fig. S17**). In the common marmoset, SN38-βGal and SN38-βGlcNAc showed half-lives of 0.972 h and 0.794 h, respectively, after a single 11-min intravenous infusion of 10 mg/kg (**Fig. 4h, Fig. S17**). In the marmoset, less than 1.6% of the SN38-βGal dose and 1.2% of the SN38-βGlcNAc dose were converted to active SN38 in the circulation, indicating high stability in the systemic circulation of non-human primates (**Extended Data Table. 2, Table S8**).

### Evaluation of SN38-βGal and SN38-**β**GlcNAc *in vivo* against lung cancer cell line-derived xenograft (CDX) models

To examine the efficacy of SN38-βGal and SN38-βGlcNAc in lung cancer, we selected NCI-H226 cells from the JFCR39 panel, because they are resistant to current molecular targeted agents and sensitive to SN38-βGal and SN38-βGlcNAc (**Extended data Fig. 3d**), and performed *in vivo* studies using the NCI-H226 CDX model. Mice bearing a subcutaneous NCI-H226 tumor were treated by systemic administration of SN38-βGal or SN38-βGlcNAc [40 mg/kg or 100 mg/kg 3 times/week for 2 weeks] or CPT-11 [40 mg/kg 3 times/week for 2 weeks] (**Fig. 4i**). SN38-βGal and SN38-βGlcNAc showed significant anti-cancer effects at both doses, with greater efficacy than CPT-11 (**Fig. 4j**). No significant body weight changes were observed in mice treated with SN38-βGal or SN38-βGlcNAc, whereas CPT-11 caused a body weight loss of about 10-20% by 7 to 14 days, followed by a return to near baseline by days 14 to 18 (**Fig. 4j**). Thus, SN38-βGal and SN38-βGlcNAc suppressed the growth of NCI-H226 tumors with lower toxicity than CPT-11.

Next, we examined an orthotopic CDX model. Mice bearing A549 orthotopic tumors were treated by systemic administration of SN38-βGal or SN38-βGlcNAc [40 mg/kg every 2 days for 9 days, followed by 80 mg/kg every 2 days for 13 days] (**Fig. 4k**). Both prodrugs showed significant anti-cancer effects without causing weight loss (**Fig. 4l-n**). Consistent with the results in H226 xenografts, both prodrugs showed comparable or superior efficacy to CPT-11 in the A549 orthotopic model (**Extended Fig. 6a-b**).

### Generalizability of the enzyme-targeting prodrug platform across diverse cytotoxic payloads *in vivo*

To demonstrate the generalizability of our prodrug development strategy, we converted multiple cytotoxic agents to which A549 cells showed differential sensitivities (doxorubicin (DOX), monomethyl auristatin E (MMAE), paclitaxel (PTX), melphalan (MEL) and etoposide (ETP)) into HEX-targeting prodrugs (DOX-βGlcNAc, MMAE-βGlcNAc, PTX-βGlcNAc, MEL-βGlcNAc and ETP-βGlcNAc) by incorporating a βGlcNAc trigger through a self-immolative linker at the hydroxyl or amino group of these drugs (**Fig. 5a**). Upon intraperitoneal administration of these prodrugs and SN38-βGlcNAc to an A549-Luc peritoneal dissemination mouse model with confirmed high HEX activity (**Fig. 5b**), SN38-βGlcNAc, DOX-βGlcNAc, MMAE-βGlcNAc and PTX-βGlcNAc—whose payloads exhibit low GI₅₀ values in A549 cells (**Extended Fig. 3d**)—demonstrated excellent anti-tumor efficacy with near-complete tumor regression observed in several mice (**Fig. 5c-f**). In contrast, MEL-βGlcNAc and ETP-βGlcNAc, with less potent payloads toward A549 cells, exhibited limited efficacy at the tested doses; however, dose escalation achieved measurable tumor growth suppression. MMAE-βGlcNAc caused transient body weight loss under the current dosing regimen without sustained toxicity, whereas no marked weight changes were observed in the other groups (**Fig. 5d**). Collectively, these results demonstrate that rational selection of cytotoxic payloads based on cellular drug sensitivity enhances therapeutic efficacy and confirms the versatility of our prodrug development platform.

**Figure 5.**
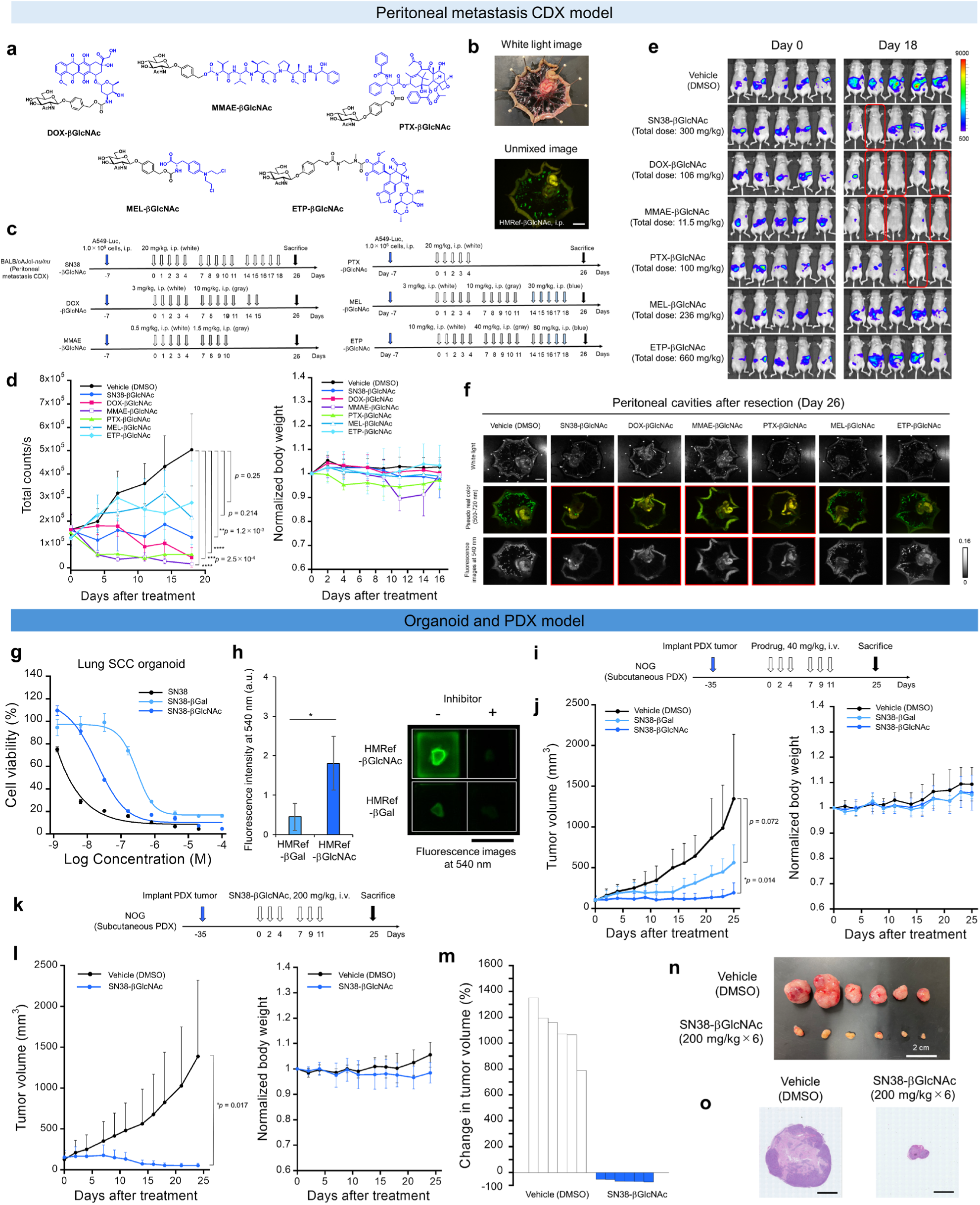
*In vivo* therapeutic efficacy of developed prodrugs in CDX and PDX models. (**a**) Chemical structures of the developed anticancer prodrugs, DOX-βGlcNAc, MMAE-βGlcNAc, PTX-βGlcNAc, MEL-βGlcNAc and ETP-βGlcNAc. (**b**) Fluorescence visualization of A549-Luc peritoneal metastases following the intraperitoneal administration of HMRef-βGlcNAc (50 µM, 300 µL). The peritoneal cavity was resected 1 h after probe administration, and subjected to fluorescence imaging and spectral unmixing (green, HMRef signal; yellow, autofluorescence signal). A549 cancer lesions exhibited HEX activities. Ex/Em = 465 nm/515 nm long pass. Scale bars, 1 cm. (**c**) Representation of the prodrug treatments in A549-Luc peritoneal metastasis mouse model. (**d**) Anti-tumor efficacy (left) and weight change (right) in A549-Luc peritoneal metastasis model mice following treatment with vehicle, SN38-βGlcNAc, DOX-βGlcNAc, MMAE-βGlcNAc, PTX-βGlcNAc, MEL-βGlcNAc and ETP-βGlcNAc (n = 5 for each group). Tumor growth inhibition was quantified using a luciferin–luciferase bioluminescence assay. Each drug was administered by intraperitoneal injection. \*\*\*\**p*<0.0001, \*\*\**p*<0.001, \*\**p*<0.01 vs. vehicle by one-way ANOVA with Tukey’s multiple comparisons test. Error bars represent s.d. (**e**) Luciferin–luciferase bioluminescence images of treated mice from (**d**). Mice with near-complete tumor regression are enclosed in red lines. (**f**) Fluorescence imaging of A549-Luc metastases after treatment with prodrugs in (**d**). After resection of peritoneal cavities, A549-Luc metastasis lesions were visualized by the topical administration of gGlu-HMRG (10 µM, 500 µL). Fluorescence images were captured 10 min after administration of the probe. Ex/Em = 465 nm/515 nm long pass. Mice with near-complete tumor regression are enclosed in red lines. Scale bars, 1 cm. (**g**) Cell viability assay of SN38, SN38-βGal and SN38-βGlcNAc for lung SCC PDO. Error bars represent s.d. (**h**) *Ex vivo* fluorescence imaging of GLB1 and HEX activities with and without inhibitors in resected lung SCC PDX tumor using HMRef-βGlcNAc and HMRef-βGal. [HMRef-βGlcNAc] = 50 µM, [HMRef-βGal] = 50 µM, [N-(n-nonyl)deoxygalactonoijirimycin] = 500 µM for HMRef-βGal, [PUGNAc] = 500 µM for HMRef-βGlcNAc. Ex/Em = 465 nm/515 nm long pass. Scale bar, 1 cm. Error bars represent s.d. (**i**) Representation of the prodrug treatment in subcutaneous lung SCC PDX tumor-bearing mice. (**j**) Anti-tumor efficacy (left) and weight change (right) in subcutaneous lung SCC PDX tumor-bearing mice following treatment with vehicle (n = 5), SN38-βGal (40 mg/kg × 6, n = 5) and SN38-βGlcNAc (40 mg/kg × 6, n = 4). One-way ANOVA with Tukey’s multiple comparisons test was used to calculate exact *p* values. Error bars represent s.d. (**k**) Representation of the high dose prodrug treatment in subcutaneous lung SCC PDX tumor-bearing mice. (**l**) Anti-tumor efficacy (left) and weight change (right) in subcutaneous lung SCC PDX tumor-bearing mice following treatment with vehicle (n = 6) and SN38-βGlcNAc (200 mg/kg × 6, n = 6). \**p*<0.05 by Welch’s *t*-test. Error bars represent s.d. (**m**) Change in tumor volume from (**l**). (**n**) White light image of resected PDX tumors after treatment from (**l**). Scale bar, 2 cm. (**o**) Representative HE-staining images of resected PDX tumors after treatment in (**l**). Scale bars, 3 mm.

### Evaluation of SN38-βGal and SN38-βGlcNAc *in vivo* against a PDX model

Finally, we evaluated the prodrugs in a subcutaneous PDX mouse model derived from lung SCC, which was negative for all currently used genetic diagnostic markers of lung cancer (**Extended Data Fig. 6c**). Notably, the tumor exhibited high HEXB expression at both the mRNA and protein levels, and its patient-derived tumor organoid (PDO) showed high sensitivity to SN38 (**Fig. 5g**). Indeed, this PDX tumor exhibited a higher expression level of HEXB than GLB1 (**Extended Data Fig. 6d**). Consistent with those findings, SN38-βGlcNAc showed a lower IC_50_ value than SN38-βGal in PDO viability assays (**Fig. 5g**). Evaluation of tumor enzyme activities using fluorescence probes revealed that HEX-reactive HMRef-βGlcNAc showed a larger fluorescence increase than GLB1-reactive HMRef-βGal (**Fig. 5h, Extended Data Fig. 6e**). This result indicates that SN38-βGlcNAc would be more effective than SN38-βGal for the treatment of this tumor. We therefore evaluated SN38-βGal and SN38-βGlcNAc *in vivo* against this PDX model [40 mg/kg 3 times/week for 2 weeks] (**Fig. 5i**). Indeed, SN38-βGlcNAc demonstrated a more potent anticancer effect than SN38-βGal, consistent with the fluorescence imaging and PDO viability assays (**Fig. 5j**, **Extended Data Fig. 6f-h, Fig. S18**). The therapeutic window of SN38-βGlcNAc is wide, and a higher dose of SN38-βGlcNAc [200 mg/kg 3 times/week for 2 weeks] resulted in a greater tumor growth inhibition compared to the 40 mg/kg treatment group, leading to a dramatic reduction in tumor volume (**Fig. 5k-n**, **Extended Data Fig. 6i-j**). This was considered to reflect the high HEX activities in the PDX tumors. No significant weight loss was observed during the treatment (**Fig. 5l**). Additionally, histopathological analysis revealed chemotherapy-specific damage in the tumor cells of the treated groups (**Fig. 5o**, **Extended Data Fig. 6j**). These results suggested that, even in patients ineligible for current molecular targeted therapies or cancer immunotherapies, highly effective therapeutic outcomes might be achieved by our present approach.

## Discussion and conclusion

Here, we establish enzyme activity as an actionable and orthogonal axis for precision cancer medicine, and introduce a broadly applicable strategy to stratify patients for selective delivery of affordable small-molecule cytotoxic agents to tumors.

Despite tremendous efforts over the past two decades, the development of safe and efficacious enzyme-reactive prodrugs has remained challenging due to the lack of suitable biomarker enzyme activities. To address this, we have established an activity-based screening strategy to identify efficient substrate/biomarker enzyme pairs for the development of highly tumor-selective prodrugs, and identified GLB1 and HEX as promising tumor-selective biomarker enzymes. In particular, the B_tot,*vitro*_ and T × T/B_tot,*vitro*_ scores introduced in this study are useful to predict the *in vivo* efficacy and safety of prodrugs. The advantage of this strategy is the ability to unbiasedly find new substrate/biomarker enzyme pairs directly by using organs and clinical cancer specimens to screen a panel of fluorescence probes. In this work, B_tot,*vitro*_ scores were determined using mouse organs, though analysis of human organ specimens is expected to provide more clinically relevant predictions for human trials.

Importantly, our activity-based prodrug development strategy affords a broadly adaptable platform for rapidly converting conventional small-molecule cytotoxic agents into highly tumor-selective therapeutics across diverse cancer types. Once suitable enzyme activities are identified in a given cancer using our strategy, the drug moiety can be replaced with a variety of cytotoxic agents—including topoisomerase inhibitors, anthracyclines, microtubule inhibitors, taxanes and alkylating agents—according to cellular sensitivity, thereby conferring tumor selectivity. As demonstrated in our previous studies on cancer imaging probes, elevated enzymatic activities vary across cancer types;^11,15,17^ however, the substrate moieties can also be easily replaced with hit substrates identified for each cancer type. Collectively, these features establish a generalizable strategy for developing tumor-selective prodrugs with diverse payloads and substrate moieties. The SN38-based prodrug CPT-11 is widely used to treat diverse cancers, despite severe nonspecific toxicities.^32–34^ As a proof of concept for our strategy, we successfully imparted high cancer selectivity to SN38. Compared with CPT-11, our GLB1-or HEX-reactive SN38-based prodrugs showed greater anticancer efficacy against enzyme-activity-positive tumors with dramatically reduced toxicity in mouse models (**Fig. 4a-d, j**). These results suggest that GLB1 and HEX are safer and more efficacious target enzymes with higher cancer selectivity than CES, which is abundantly expressed in the liver and gut. Furthermore, the developed prodrugs exhibited a significantly broader therapeutic window compared with CPT-11 (**Fig. 4a, Extended Data Fig. 4b, Table S7**), and thus can be administered at higher doses than CPT-11, thereby affording greater therapeutic efficacy. They may therefore be promising alternatives to CPT-11.

Notably, the developed prodrugs were highly effective against various cancer cell lines that are insensitive to current molecular targeted therapies in JFCR39 panel analysis (**Extended data Fig. 3d**). Indeed, in the efficacy studies using lung cancer PDO and PDX mouse models that were negative for current molecular targets including PD-L1 (**Extended data Fig. 6c**), but exhibited high HEX activity, SN38-βGlcNAc showed remarkably potent tumor growth inhibition (**Fig. 5k-o**). Since GLB1 and HEX activities are broadly elevated in lung cancer patients (**Table 1**), these results suggested that the developed prodrugs may be effective in at least some of the approximately 40-60% of NSCLC patients who are not eligible for current molecular targeted therapies and cancer immunotherapies.^41–43^ Furthermore, combination with PD-1-based immunotherapies could enhance the therapeutic efficacy of SN38-βGlcNAc, suggesting a potential synergistic benefit with existing cancer immunotherapies. (**Extended data Fig. 7a-e**). In addition, GLB1 and HEX activities or expression levels are elevated in other cancers such as breast, prostate and brain, ^17,44,45^ so the developed prodrugs may also be promising candidates for the treatment of these cancers.

Since the companion fluorescence probes enabled rapid and sensitive detection of GLB1 and HEX activities in clinical lung cancer specimens (**Extended Fig. 8a-g, Fig. S19-22**), activity-based diagnostics with these probes can be used to select the most appropriate target enzyme for individual tumors. Cytotoxic payloads with clinically validated efficacy in each cancer type should be prioritized for prodrug development. In future applications, PDO has emerged as an approach to predict responses to chemotherapeutic agents *in vitro*,^46^ and our results further suggest that PDO-based assays may help to refine the selection of therapeutic payloads tailored to individual patients (**Fig. 5g**). Additionally, the sensitivity to SN38-βGal and SN38-βGlcNAc strongly depends on the expression of Schlafen 11 (SLFN11), a restriction factor for replicative stress induced by DNA-targeting anticancer drugs (**Fig. S23**),^47^ suggesting that assessing such molecular determinants may improve efficacy prediction. Although tumor heterogeneity may lead to enzyme-negative cell populations, the bystander effect of the released payload is expected to mitigate this limitation, consistent with the observed efficacy in the PDX model (**Fig. 5l-n**). Co-administration of different prodrugs may also increase therapeutic efficacy by targeting multiple enzymes.

In conclusion, we propose a novel concept of small-molecule precision medicine targeting biomarker enzymatic activities, in which cancer patients are stratified based on activity-based diagnostics, and treated with prodrugs optimized for the elevated enzyme activities in their tumors. In other words, fluorescence detection of target enzyme activities in surgically resected or biopsy specimens enables a personalized treatment strategy using enzyme-reactive prodrugs selected to match the patient’s enzyme activity profile. We believe this approach offers a versatile and economically sustainable platform for precision oncology based on affordable small-molecule diagnostic probes and prodrugs.

## Methods

### Synthesis and characterization of compounds

Synthetic protocols are given in the Supplementary Information. NMR spectra were recorded on a Bruker AVANCE III and 400 Nanobay at 400 MHz for ^1^H NMR and 101 MHz for ^13^C NMR. High-resolution mass spectra (HRMS) were measured with a MicroTOF (Bruker).

### Human surgical lung specimens

74 lung cancer patients who underwent surgical treatment during 2013-2021 at the University of Tokyo Hospital were included in this study. Some patients were duplicated in plural analyses. Fresh and frozen human surgical lung specimens were used. Fresh surgical specimens were evaluated within 60-120 min after resection. Frozen specimens were frozen immediately after resection, stored at-80°C, and thawed at room temperature before use.

### Animal studies

All animal studies were performed in accordance with the policies of the Animal Ethics Committee of the University of Tokyo. Jcl:ICR mice, BALB/cAJcl-*nu*/*nu* mice, C57BL/6 mice and NOD.Cg-Prkdc<scid>Il2rg/ShiJic (NOG) mice were purchased from CLEA Japan, Inc., Tokyo, Japan. Tumor volumes were measured with a digital caliper using the formula L × W^2^/2.

### Evaluation of *in vitro* background score of each fluorescence probe

Organs of healthy female 7-week-old Jcl:ICR mice were resected, shredded with scissors, and transferred to a Lysing Matrix D (6913100, MP Biomedicals). Then, 1 mL of tissue protein extraction reagent (T-PER) (Thermo Fisher Scientific) was added and the mixture was homogenized in a FastPrep-24 (MP Biomedicals). After centrifugation, the supernatant was collected, and protein quantification was performed using a Pierce BCA Protein Assay Kit (Thermo Fisher Scientific). Fluorescence probes (1 µM) were incubated with 0.1 mg/mL lysate of each organ in PBS (-) in 384-well black plate (4514, Corning) for 1 h at 37 °C. Fluorescence intensity was measured using Envision plate reader (PerkinElmer). Excitation/emission (Ex/Em) settings were FITC 485 (485/14 nm) and FITC 535 (535/25 nm), respectively. The conversion rate of probe to fluorophore (CVR) in each organ was calculated by dividing the fluorescence intensity of the probe by that of the resultant fluorophore at the same measurement time. The *in vitro* background score in each organ (B_org,*vitro*_) was calculated as the product of CVR and organ weight (kg). The *in vitro* total background score (B_tot,*vitro*_) was calculated as the sum of the B_org,*vitro*_ scores in the examined organs. Organ weights were obtained from healthy female 7-week-old Jcl:ICR mice (n = 3).

### Direct screening of fluorescence probes with human surgical specimens

A 50 µM solution of fluorescence probe (200 µL) in PBS (-) containing 0.5% *v*/*v* DMSO as a co-solvent was added to each well of an 8-well chamber (µ-Slide 8 well; Ibidi) containing a human surgical specimen (tumor or non-tumor tissue). Fluorescence intensities and images of specimens were recorded with the Maestro *in-vivo* imaging system (CRi Inc.) before and at 1, 3, 5, 10, 20 and 30 min after probe administration (Ex/Em = 490 nm/550 nm long-pass). The tunable filter was automatically adjusted in 10 nm increments from 500 nm to 720 nm, while the camera sequentially captured images at each wavelength interval. Fluorescence at 540 nm was extracted and fluorescence intensities were quantified by drawing ROIs with the Maestro software. Exposure time was set at 100-50 msec depending on the fluorescence intensity from specimens. Stage and lamp settings: position 1.

### Evaluation of *in vivo* background scores in each organ

A PBS (-) solution of each fluorescence probe, HMRef-βGal, HMRef-βGlcNAc, gGlu-HMRG, KK-HMRG and ZFR-HMRG (5 mg/kg) containing 20% or less DMSO (*v/v*) as a co-solvent (100 µL) was intravenously administered to 7-week-old female Jcl:ICR mice (n = 3 for each group). After 20 min, mice were sacrificed, and major organs were resected. The contents of the stomach and guts were removed. Fluorescence images (Ex/Em = 490 nm/550 nm long-pass) of resected organs were captured using the Maestro *in vivo* imaging system (CRi Inc.) with an exposure time of 100-500 msec depending on the fluorescence intensity. Stage and lamp settings: position 1. The *in vivo* background score (B_org,*vivo*_) was calculated as the products of the fluorescence intensity and organ weight (kg). The *in vivo* total background scores (B_tot,*vivo*_) were calculated as the sum of the B_org,*vivo*_ scores in the examined organs.

### Evaluation of systemic selectivity in tumor-bearing mice

For lung orthotopic A549 tumor-bearing BALB/cAJcl-*nu*/*nu* mice, a PBS (-) solution of HMRef-βGlcNAc (5 mg/kg) containing 20% or less DMSO (*v/v*) as a co-solvent (100 µL) was intravenously administered via the tail vein. After 30 min, mice were sacrificed, and fluorescence images of resected lung or thoracotomized mice were captured using the Maestro *in vivo* imaging system (CRi Inc.). For the PDX study, NOG mice bearing a lung SCC PDX tumor (Product number: F-PDX-000162 (DLUN013), Fukushima Translational Research Foundation) derived from a 54-year-old man were used. A PBS (-) solution of each fluorescence probe, HMRef-βGlcNAc, HMRef-βGal, gGlu-HMRG and KK-HMRG (5 mg/kg) containing 20% or less DMSO (*v/v*) as a co-solvent (100 µL) was intravenously administered via the tail vein. After 20 min, mice were sacrificed, and major organs and tumors were resected. Fluorescence images (Ex/Em = 490 nm/550 nm long-pass) of resected organs and tumors were captured using the Maestro *in vivo* imaging system (CRi Inc.) with an exposure time of 100-500 msec. Stage and lamp settings: position 1. Unmixing of fluorescence spectra was performed using Maestro software.

### DEG assay

DEG assay was performed according to the reported method.^30^ Human surgical lung cancer specimens were lysed using Lysing Matrix D (6913100, MP Biomedicals) and a FastPrep-24 homogenizer (MP Biomedicals).

### Histological and immunohistochemical analysis

Excised specimens were immediately fixed with 10 % formaldehyde for at least 72 h. Formalin-fixed paraffin-embedded tissues were sectioned at 4 µm thickness and stained with hematoxylin and eosin for histopathological evaluation. Protocols of IHC analysis are given in the Supplementary Information. Experienced pathologists examined each sample in a blind manner.

### Confocal imaging of enzyme activities in cultured cancer cells

For fluorescence microscopy, about 10,000 cells in 200 µL of medium supplemented with 10% fetal bovine serum and 1% penicillin/streptomycin/amphotericin B were seeded in each well of an 8-well chamber (µ-Slide 8-well; Ibidi) and cultured for 1 day. The medium was replaced with 200 µL of phenol red-and serum-free RPMI-1640 or DMEM containing 20 µM fluorescence probe. After incubation at 37°C for 1 h or 3 h in a humidified incubator under 5% CO_2_ in air, differential interference contrast (DIC) and fluorescence images were captured using a confocal fluorescence microscope (TCS SP8 STED, Leica). Fluorescence intensities were quantified using ImageJ software. The light source was a white light laser. Ex/Em = 498 nm/500-600 nm.

### Evaluation of *in vitro* cytotoxicity by MTT assay

The *in vitro* cytotoxicity to A549, NCI-H226, SHP-77 and HEK/lacZ cells was determined using a Cell Proliferation kit I (MTT) (11465007001, Roche). Briefly, 10,000 cells/well in 100 µL cultured medium containing 10% fetal bovine serum (FBS) and 1% penicillin/streptomycin were plated in 96-well plates and incubated for 24 h at 37°C. FBS-containing medium was removed and replaced with OptiMEM (Thermo Fisher Scientific) containing 1% penicillin/streptomycin. The cells were treated with serial dilutions of SN38-βGal, SN38-βGlcNAc, SN38 or CPT-11 dissolved in DMSO and further incubated for 72 h at 37°C. Then, 10 µL of MTT labeling reagent (3-[4,5-dimethylthiazol-2-yl]-2,5-diphenyltetrazolium bromide) was added to each well (final concentration 0.5 mg/mL), and incubation was continued for 4 h to form a colored product, formazan. Solubilization buffer (10% SDS in 0.01 M HCl; 100 µL) was added to each well, and incubation was continued for 24 h. The absorbance at 560 nm was measured using an Envision plate reader to evaluate cell viability. Absorbance at 720 nm was used as the reference. Since the FBS affected the stability of prodrugs, the assay was performed using Opti-MEM medium containing no serum.

### Time-lapse imaging analyses of cell growth inhibition

A549 (1,300 cells/well) in 50 µL of RPMI-1640 medium supplemented with 5% heat-inactivated human serum (H3667-100ML, Sigma-Aldrich) were plated in a 384-well black plate (Revvity) with SPY650-DNA (CY-SC501, Cosmo Bio Co., Ltd.) at 250 nM and SYTOX Green (S34860, Thermo Fisher) at 30 nM. At 24 h after cell plating, DMSO, SN38, CPT-11, SN38-βGal or SN38-βGlcNAc in 25 µL medium with 250 nM SPY650-DNA and 30 nM SYTOX Green were added to each well and the fluorescence images of SPY650-DNA-stained cells and SYTOX Green-stained cells, as well as bright-field images, were obtained every hour using a live cell high-throughput imaging system, Operetta CLS (Revvity). Ex/Em = 630/655-760 nm for SPY650-DNA and 475/500-550 nm for SYTOX Green. Quantitative image analysis was performed using Harmony 4.9 software (Revvity).^47,48^

### Evaluation of GI_50_ values in the JFCR39 cancer cell panel

GI_50_ values were determined by colorimetric assay using sulforhodamine B as reported.^38,49^ For SN38, CPT-11, SN38-βGal and SN38-βGlcNAc assays, RPMI-1640 medium supplemented with 5% heat-inactivated human serum (H3667-100ML, Sigma-Aldrich) was used. For other drug assays, RPMI-1640 medium supplemented with 5% fetal bovine serum was used.

### Repeated dose toxicity studies

SN38-βGal, SN38-βGlcNAc and CPT-11 were intravenously administered to 7-week-old female Jcl:ICR mice via the tail vein at a dose of 40 mg/kg three times per week for 3 weeks (n = 8 for each group). The vehicle group received the corresponding volume of DMSO. Body weight changes were monitored for 21 days following the initial administration. On day 21, all mice were sacrificed, and liver, kidney, spleen, intestine and colon were collected for pathological examination. Liver, kidney and spleen weights were measured. Blood samples were collected for biochemical analysis. The statistical significance of weight changes was analyzed by one-way ANOVA with Tukey’s multiple comparisons test.

### Evaluation of the locomotion of mice

Trajectories of mice were recorded in a homemade open-square field measuring 40 × 40 × 40 cm with white walls and a black floor. A camera was placed above the apparatus to monitor the animal’s behavior at 30 fps. Immediately after intravenous drug administration (40 mg/kg) to 7-week-old female Jcl:ICR mice, each mouse was placed in the open field, and its trajectory was monitored for 15 min (n = 4 for each group). Mice were not allowed to explore the open field before the experiment. Data analyses were performed using MATLAB (MathWorks) and ImageJ.^50^ The statistical significance of differences in locomotion was analyzed by one-way ANOVA with Tukey’s multiple comparisons test.

### Measurement of AChE inhibition

AChE inhibition assays were performed using an Acetylcholinesterase Activity Assay Kit, Fluorometric, SensoLyte 520 (AS-72242; Funakoshi) according to the manufacturer’s instructions. Assay buffer containing serial concentrations of SN38-βGal, SN38-βGlcNAc, or CPT-11 (10 µL) was dispensed into a 96-well black plate (3915, Costar), followed by the addition of assay buffer containing human recombinant AChE (0.05 ng, 40 µL). Subsequently, the AChE substrate and acetylcholine were added in assay buffer according to the kit protocol (50 µL). Fluorescence intensity was measured using an EnVision plate reader (PerkinElmer). Ex/Em = FITC 485 (485/14 nm)/FITC 535 (535/25 nm). Neostigmine bromide, was used as the AChE inhibitor control.

### Pharmacokinetic studies

Pharmacokinetic studies were conducted in mice and common marmosets. Female Jcl:ICR mice with free access to food and water received intravenous administration of SN38-βGal and SN38-βGlcNAc (40 mg/kg) dissolved in PBS (-) containing 10% or less DMSO (n = 3 for each group). Blood samples were collected from the inferior vena cava at 5, 15, 30, and 60 min post-dose under medetomidine/midazolam/butorphanol anesthesia. Blood samples were treated with heparin sodium and 100 μM glycosidase inhibitor (N-(n-nonyl)deoxygalactonojirimycin for SN38-βGal or PUGNAc for SN38-βGlcNAc) to prevent enzymatic degradation, followed by centrifugation at 1,200 × g for 15 min at 4°C to obtain plasma, which was stored at −80°C until analysis. For marmoset studies, male 39-month-old common marmosets were used (n = 3 for each group). SN38-βGal or SN38-βGlcNAc (1.0 mg/mL) was intravenously administered via the tail vein at a dose of 10 mg/kg (10 mL/kg) using an infusion pump at a constant rate of 350 μL/min (approximately 11 min). Blood samples (220 μL) were collected at 5, 15, 30, and 45 min, and 1, 2, 3, and 6 h post-infusion using heparinized syringes and centrifuged at 1,200 × g for 15 min at 4°C. The plasma was immediately mixed with an equal volume of 200 μM N-(n-nonyl)deoxygalactonojirimycin (for SN38-βGal) or PUGNAc (for SN38-βGlcNAc) to prevent enzymatic degradation, then stored at −80°C until analysis. For sample preparation, plasma samples (50 μL) were mixed with 200 μL of MeOH/MeCN (1:1, *v*/*v*) containing camptothecin as an internal standard, incubated for 10 min at room temperature, and centrifuged at 1,700 × g for 15 min at 4°C. The supernatant was filtered using a Cosmo Spin Filter G (06549-44, Nacalai Tesque) by centrifugation at 12,000 × g for 5 min at 4°C. The filtrate (50 μL) was diluted with Milli-Q water (50 μL) and subjected to LC-MS analysis (Aquity UPLC H-Class Plus, Waters). Plasma concentrations were quantified using calibration curves generated from plasma standards prepared using the same procedure. For the marmoset analysis, the initial concentration was set to zero. Plasma half-lives were calculated by estimating the elimination phase using the concentration values at 15–60 min for mice and 2–6 h for marmosets.

### Therapeutic efficacy in H226 subcutaneous tumor-bearing mice

Female 7-to 9-week-old BALB/cAJcl-*nu*/*nu* mice with free access to food and water before experiments were used. NCI-H226 cells (1.5 × 10^6^ cells cells) were injected into the right flank, and tumor growth was monitored until the median size reached about 160 mm^3^. Drugs were intravenously administered via the tail vein (SN38-βGal and SN38-βGlcNAc: 40 mg/kg or 100 mg/kg three times per week for 2 weeks; CPT-11: 40 mg/kg three times per week for 2 weeks). The dosing solutions were prepared in PBS (-) containing less than 24 µL of DMSO as a cosolvent, and were administered at a total volume of 100 µL. Tumor volumes and weight changes were monitored over 4 weeks after initial administration. The statistical significance of differences in anticancer efficacy was analyzed by one-way ANOVA with Tukey’s multiple comparisons test.

### Therapeutic efficacy in A549 lung orthotopic tumor-bearing mice

Female 7-week-old BALB/cAJcl-*nu*/*nu* mice with free access to food and water before experiments were used. For tumor implantation, luciferase-expressing A549-Luc cells (1.0 × 10^6^ cells) were suspended in 50 µL of PBS (-) containing 5% Matrigel and injected into the left lung under medetomidine/midazolam/butorphanol anesthesia. Before treatment, tumor implantation was confirmed by luciferin-luciferase bioluminescence imaging using an IVIS *in vivo* imager (PerkinElmer). Eight days after tumor implantation, SN38-βGal and SN38-βGlcNAc were intravenously administered via the tail vein [40 mg/kg every 2 days for 9 days, followed by 80 mg/kg every 2 days for 13 days]. The dosing solutions were prepared in PBS (-) containing less than 19 µL of DMSO as a cosolvent (total volume: 100 µL). Lung tumor growth was monitored by bioluminescence imaging under isoflurane anesthesia, 20 min after intraperitoneal administration of luciferin (3.0 mg) dissolved in 200 µL PBS (-). Weight changes were monitored for 25 days after initial administration of prodrugs. The statistical significance of differences in anticancer efficacy was analyzed by one-way ANOVA with Tukey’s multiple comparisons test. On day 29, lungs were resected 20 min after intravenous administration of 5 mg/kg HMRef-βGlcNAc. Fluorescence images (Ex/Em = 490 nm/550 nm long-pass) of resected lungs were captured using the Maestro *in vivo* imaging system (CRi Inc.) to compare the therapeutic efficacy. Stage and lamp settings: position 2B.

### Therapeutic efficacy in A549 peritoneal metastasis mouse model

Female 7-week-old BALB/cAJcl-*nu*/*nu* mice with free access to food and water before experiments were used. For tumor implantation, luciferase-expressing A549-Luc cells (1.0 × 10^6^ cells) were suspended in 200 µL of PBS (-) and injected intraperitoneally. Before treatment, tumor implantation was confirmed by luciferin-luciferase bioluminescence imaging using an IVIS *in vivo* imager (PerkinElmer). Seven days after tumor implantation, SN38-βGlcNAc, DOX-βGlcNAc, MMAE-βGlcNAc, PTX-βGlcNAc, MEL-βGlcNAc and ETP-βGlcNAc were administered by intraperitoneal injection. The dosing solutions were prepared in PBS (-) containing less than 17 µL of DMSO as a cosolvent (total volume: 100 µL). Peritoneal metastasis growth was monitored by bioluminescence imaging under isoflurane anesthesia, 20 min after intraperitoneal administration of luciferin (3.0 mg) dissolved in 200 µL PBS (-). Weight changes were monitored for 16 days after initial administration of prodrugs. The statistical significance of differences in anticancer efficacy was analyzed by one-way ANOVA with Tukey’s multiple comparisons test. On day 26, mice were sacrificed and the peritoneal cavities were resected. Fluorescence images (Ex/Em = 490 nm/550 nm long-pass) of resected cavities were captured using the Maestro *in vivo* imaging system (CRi Inc.) 10 min after topical application of 50 µM gGlu-HMRG PBS(-) solution (500 µL) to visualize A549-Luc peritoneal metastases. Stage and lamp settings: position 2.

### PDO viability assay

Lung SCC PDO (Product number: F-PDO-000124 (RLUN016-2), Fukushima Translational Research Foundation) derived from tumor (F-PDX-000162 (DLUN013) was cultured in Cancer Cell Expansion Media plus (032-25745, Wako) at 37°C in a humidified incubator under 5% CO_2_ in air. Centrifugation at 200 G for 2 min afforded a pellet (150 µL), which was suspended in 15 mL of medium. The required volume of suspension was collected, and cell aggregates were dissociated using a CellPet FT (JTEC Corporation) with a 70 µm filter. The resulting suspension was diluted 30-fold, seeded in 96-well spheroid plates (4520, Corning) at 150 µL/well and incubated overnight. Serial dilutions of SN38, SN38-βGal and SN38-βGlcNAc in DMSO (0.15 µL) were added and incubated for 6 days at 37°C in a humidified incubator under 5% CO_2_ in air. CellTiter-Glo 3D (G9683, Promega) was added to each well, and vortexed for 5 min. After 10 min incubation at room temperature, chemiluminescence was measured using an EnSpire 2300 (PerkinElmer) to quantify ATP, and organoid viability and IC_50_ values were determined.

### Therapeutic efficacy in PDX mouse model

Lung SCC PDX tumor (Product number: F-PDX-000162 (DLUN013), Fukushima Translational Research Foundation) derived from a 54-year-old man was used. Female NOG mice with free access to food and water before experiments were transplanted with tumor fragments (30 mg) into the shaved right flank under medetomidine/midazolam/butorphanol anesthesia. Tumor growth was monitored until the median size reached about 100-140 mm^3^. SN38-βGal and SN38-βGlcNAc were intravenously administered via the tail vein at a dose of 40 mg/kg three times per week for 2 weeks. The dosing solutions were prepared in PBS (-) containing less than 9 µL of DMSO as a cosolvent (total volume: 100 µL). For high dose treatment, SN38-βGlcNAc was intravenously administered via the tail vein at a dose of 200 mg/kg three times per week for 2 weeks. The dosing solutions were prepared in PBS (-) containing less than 20 µL of DMSO as a cosolvent (total volume: 100 µL). The statistical significance of differences in anticancer efficacy was analyzed by one-way ANOVA with Tukey’s multiple comparisons test.

## Data Availability

All data and materials are available upon request.

## Acknowledgements

This research was supported by Grants-in-Aid for Scientific Research (KAKENHI) (JP22K20528, JP23K14317, JP25K18593 to K.F.), Japan Science and Technology Agency (JST) ACT-X program (JPMJAX222G to K.F.), Masason Foundation (to K.F.), KAKENHI (JP22H04922(AdAMS) to S.D.), and Nippon Foundation (2022003175 to S.D.), KAKENHI (JP19H05632 and JP24H00050 to Y.U.), JST Mirai Program (JPMJMI24G2 to Y.U.), and JST Moonshot Research and Development Program (JPMJMS2022 to Y.U.). We thank Prof. Yoshiko Miura and Dr.

Tomonobu Koizumi for their support through the JST ACT-X program. We thank Tokyo Central Pathology Laboratory Co., Ltd. for preparing unstained histological slide sections and performing IHC staining for HEX as a fee-for-service. We thank BioSafety Research Center Inc. and FUJIFILM Systems Co., Ltd. for conducting the biochemical analysis of blood as a fee-for-service. We thank CLEA Japan, Inc. for conducting the collection of blood samples of common marmosets in pharmacokinetic study as a fee-for-service. We thank Fukushima Translational Research Foundation for conducting PDO viability assay and providing the PDX tumor (DLUN013) as a fee-for-service.

## Author contributions

K.F. and Y.U. designed and supervised the project. K.F. conducted the experimental work. K.F., Mako.K., S.D., Minoru.K., R.K. and Y.U. analyzed the results. K.F. wrote the manuscript with input from all the authors. S.K. and J.N. helped conduct experiments involving human surgical specimens. S.I. and M.T. partly helped conduct the evaluation of probe CVR and background scores. R.H. conducted pathological evaluation. K.K. conducted the MTT assay for SHP-77 cells. S.D., K.Y., Y.N., Y.O., S.I. and Y.H. helped conduct the experiment using the JFCR39 cell panel. R.T. conducted QM/MM calculations. N.M. and Y.I. helped conduct the evaluation of trajectories. A.N. helped the pharmacokinetic analysis. The authors declare no competing interests.

## Extended data (10 figures or tables)

**Extended data Figure 1.**
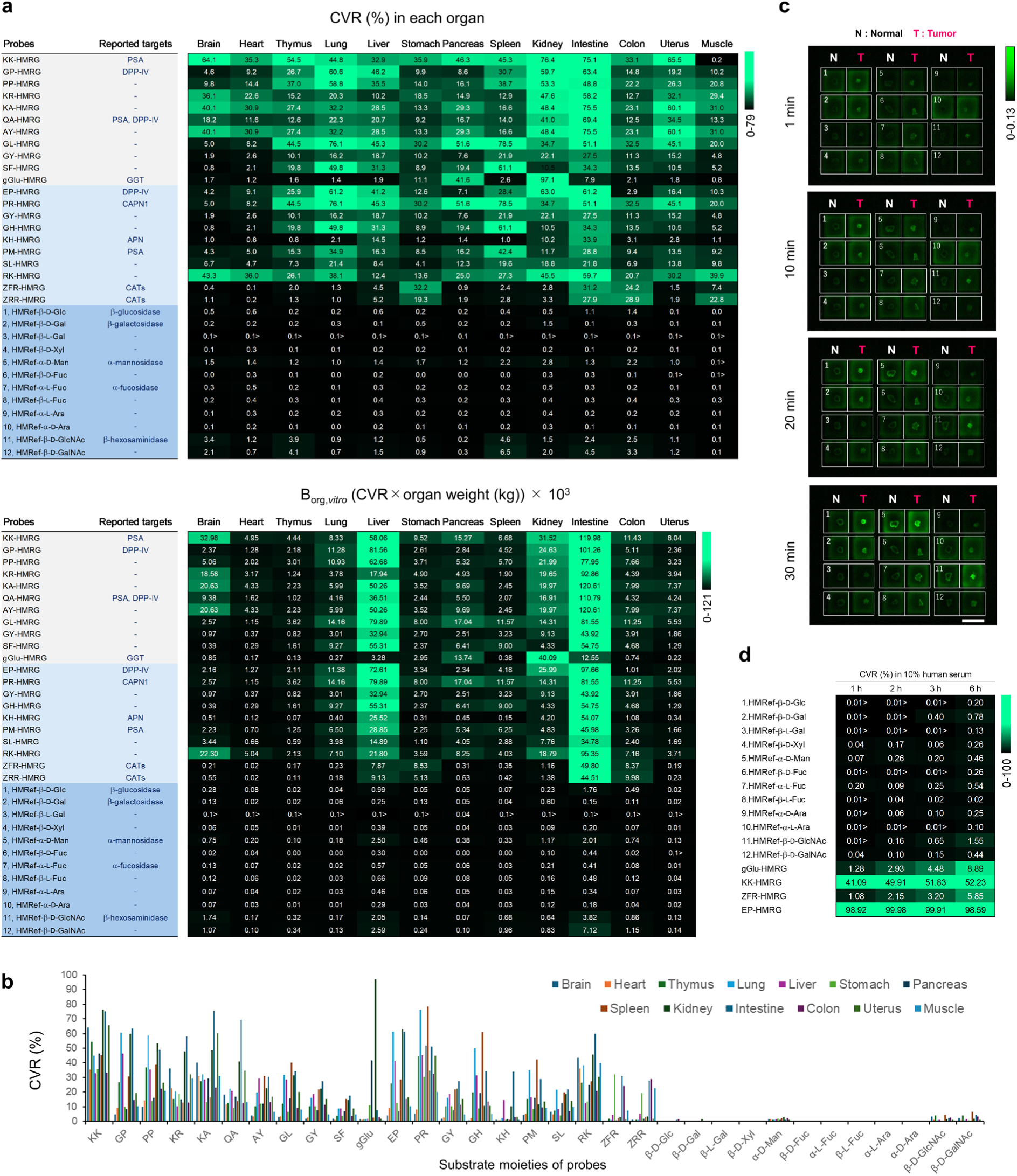
Activity-based screening of fluorescence probes for efficient prodrug development. (**a**) Heat map of CVR at 1 h of fluorescence probes in each mouse organ lysate (upper). Incubation was performed at 37°C for 1 h. [Fluorescence probes] = 1 µM, [organ lysate] = 0.1 mg/mL. The *in vitro* background scores (B_org,*vitro*_) × 10^3^ of evaluated fluorescence probes (bottom). Exact values corresponding to **Fig. 1b** are shown. The fluorescence probes were incubated with each mouse organ lysate for 1 h. Background scores for each organ were calculated by multiplying the CVR of fluorescence probes by the organ weight (kg). [fluorescence probe] = 1 µM, [organ lysate] = 0.1 mg/mL. (**b**) Comparison of CVR at 1 h of probes in each mouse organ lysate from (**a**). (**c**) Representative example of time-dependent fluorescence images of *ex vivo* screening using clinical lung cancer specimen. The probe number of the applied probe is shown on each well. [Fluorescence probes] = 50 µM. Ex/Em = 465 nm/515 nm long pass. Scale bar, 1 cm. (**d**) CVR of fluorescence probes in 10% *v/v* human serum. Incubation was performed at 37°C. Glycosidase-reactive probes showed lower CVR compared with aminopeptidase-reactive probes. [Fluorescence probes] = 1 µM.

**Extended data Figure 2.**
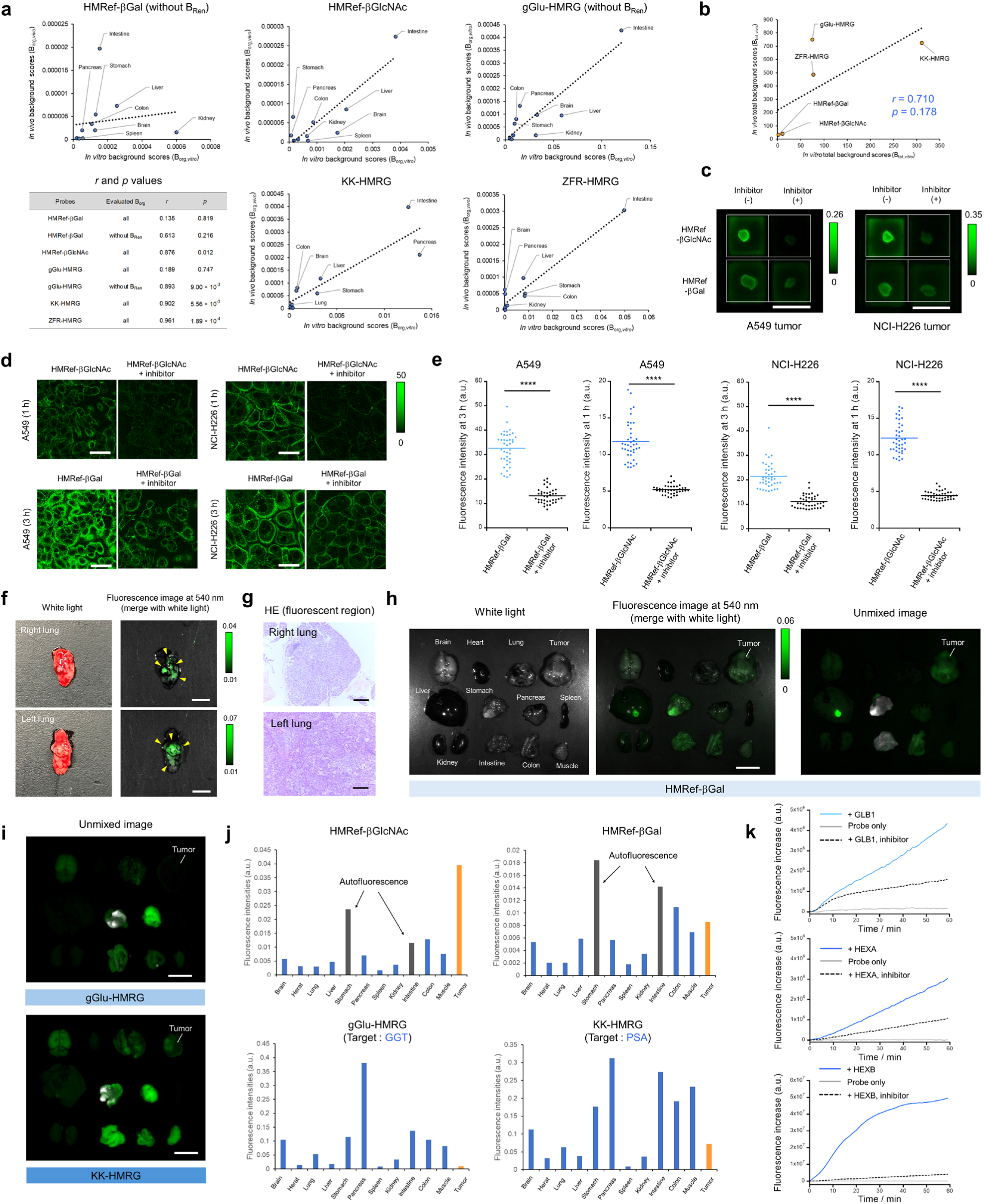
Evaluation of *in vivo* cancer selectivity and target reactivity of hit fluorescence probes. (**a**) Correlations between B_org,*vitro*_ and B_org,*vivo*_ for each fluorescence probe. Strong positive trends were observed in HMRef-βGlcNAc, KK-HMRG and ZFR-HMRG. The *in vivo* background scores of kidney (B_Ren_) appeared as an outlier for HMRef-βGal and gGlu-HMRG; upon its exclusion, a clear positive correlation was observed. This likely reflects the lack of pharmacokinetic considerations in B_org,*vitro*_ scores, as well as the underestimation of green fluorescence in the kidney due to its coloration in fluorescence imaging. (**b**) Correlations between B_tot,*vitro*_) and B_tot,*vivo*_. Although the sample size was limited (n = 5), a positive trend was observed (*r* = 0.710). (**c**) *Ex vivo* fluorescence imaging of glycosidase activities of resected NCI-H226 subcutaneous tumors using HMRef-βGal or HMRef-βGlcNAc with and without each glycosidase inhibitor. Fluorescence intensities were significantly suppressed in the presence of inhibitors. [Fluorescence probes] = 50 µM, [N-(n-nonyl)deoxygalactonoijirimycin] = 500 µM for HMRef-βGal, [PUGNAc] = 500 µM for HMRef-βGlcNAc. Ex/Em = 465 nm/515 nm long pass. Scale bars, 1 cm. (**d**) Live cell fluorescence imaging of HMRef-βGal or HMRef-βGlcNAc in A549 and NCI-H226 cells with and without each inhibitor. Incubations were performed at 37°C for 1 h or 3 h. The excitation and emission wavelengths were 495 nm/505-540 nm. [Fluorescence probes] = 20 µM, [N-(n-nonyl)deoxygalactonoijirimycin] = 50 µM for HMRef-βGal, [PUGNAc] = 50 µM for HMRef-βGlcNAc. Scale bars, 50 µm. (**e**) The dot-plot diagram of fluorescence intensities of A549 and NCI-H226 single-cell at 1 h or 3 h after addition of HMRef-βGal or HMRef-βGlcNAc with and without each inhibitor (n = 40 for each group). The line in the diagram represents the average. Fluorescence intensities were significantly suppressed in the presence of inhibitors. \*\*\*\**p*<0.0001 by Welch’s t-test. [Fluorescence probes] = 20 µM, [N-(n-nonyl)deoxygalactonoijirimycin] = 50 µM for HMRef-βGal, [PUGNAc] = 50 µM for HMRef-βGlcNAc. (**f**) White light and fluorescence images of resected lungs after fluorescence imaging shown in **Fig. 2e**. Selective fluorescence activation in cancer regions was clearly observed in resected lungs. Lung cancer lesions are indicated with yellow arrows. Scale bars, 5 mm. (**g**) Histological analysis of fluorescent regions in resected lungs in (**f**). Scale bars, 200 µm. (**h**) *Ex vivo* white light (left), fluorescence image (middle) and unmixed fluorescence image (green, HMRef signal; white, autofluorescence signal) (right) of freshly resected tumor and each organ 20 min after intravenous administration of 5 mg/kg HMRef-βGal to a subcutaneous lung cancer PDX model. Scale bar, 1 cm. (**i**) *Ex vivo* fluorescence unmixed images (green, HMRG signal; white, autofluorescence signal) of freshly resected tumor and each organ 20 min after intravenous administration of 5 mg/kg gGlu-HMRG or KK-HMRG to a subcutaneous lung cancer PDX model mice from **Fig. 2f**. Ex/Em = 465 nm/515 nm long pass. Scale bar, 1 cm. (**j**) Fluorescence intensities at 540 nm of each organ and tumor from **Fig. 2f** and (**h**). Blue bars represent fluorescence intensities derived from probe activation in each organ. Gray bars represent fluorescence intensities mainly derived from autofluorescence of organs. Orange bars represent fluorescence intensities derived from probe activation in tumor tissues. (**k**) Evaluation of the reactivity of fluorescence probes with GLB1, HEXA and HEXB recombinant proteins. HMRef-βGal and HMRef-βGlcNAc exhibited fluorescence increases in the presence of corresponding target enzymes. Observed fluorescence increases of these probes were significantly suppressed in the presence of the corresponding inhibitor. [Fluorescence probes] = 1 µM, [inhibitors] = 10 µM, [recombinant proteins] = 10 ng/mL. N-(n-Nonyl)deoxygalactonoijirimycin was used as an inhibitor of GLB1. PUGNAc was used as an inhibitor of HEXA and HEXB.

**Extended data Figure 3.**
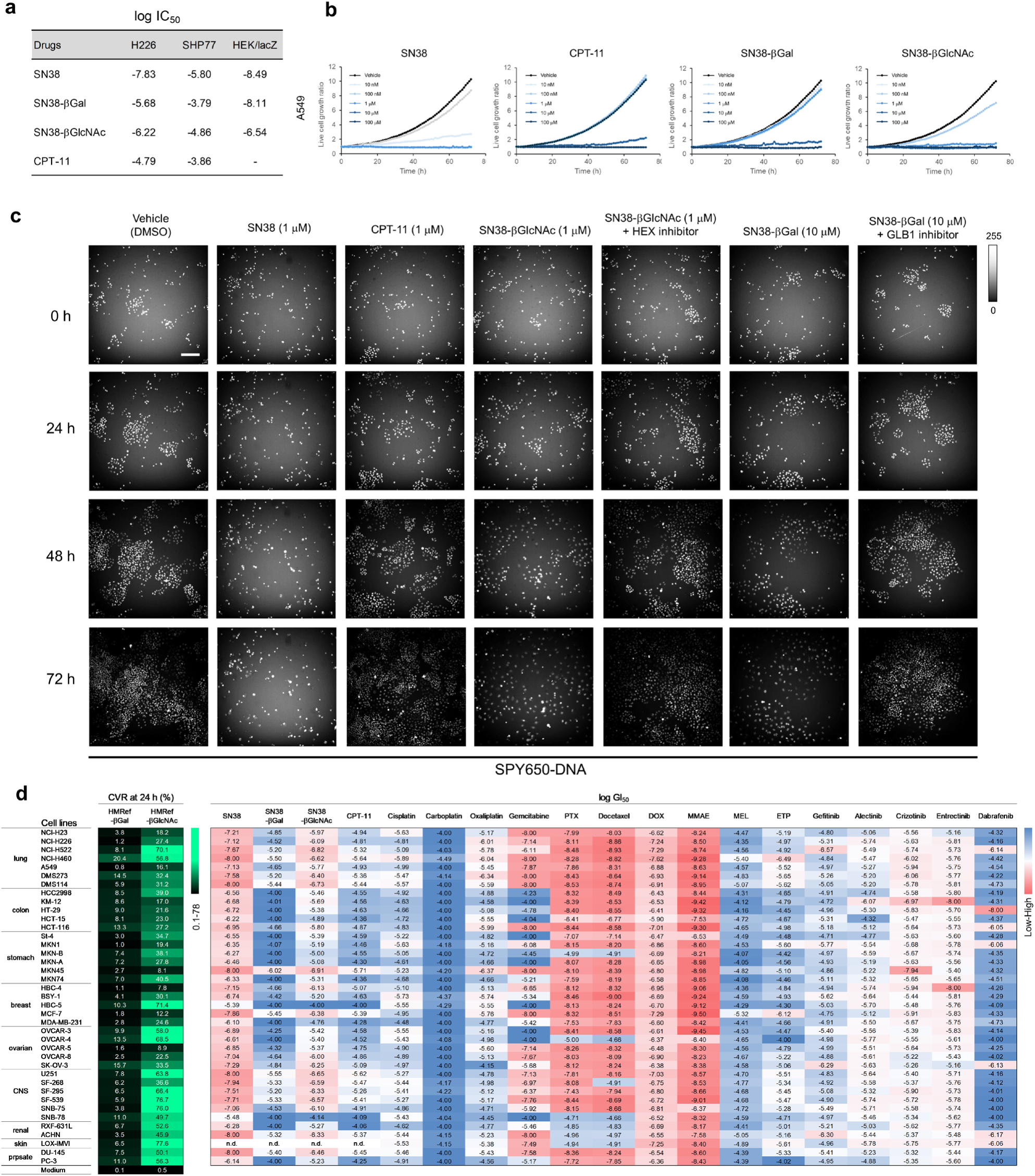
*In vitro* evaluation of the developed prodrugs in various cancer cell lines. (**a**) Log IC_50_ values observed in MTT assay from **Fig. 3d**. (**b**) Time-dependent cell growth ratio of A549 cells in the presence of various concentrations of SN38, CPT-11, SN38-βGal or SN38-βGlcNAc. (**c**) Time-dependent fluorescence images of A549 cells in the presence of each drug. Nuclei were stained with SPY650-DNA. [SN38] = 1 µM, [CPT-11] = 1 µM, [SN38-βGlcNAc] = 1 µM, [SN38-βGal] = 10 µM, [PUGNAc] = 20 µM for SN38-βGlcNAc, [N-(n-nonyl)deoxygalactonoijirimycin] = 20 µM for SN38-βGal. Scale bar, 200 µm. Treatment with prodrugs induced S-phase arrest, whereas co-treatment with the glycosidase inhibitors suppressed this effect and allowed cell proliferation. (**d**) Analysis of CVR of fluorescence probes at 24 h (left) and log GI_50_ values of SN38, CPT-11, SN38-βGal, SN38-βGlcNAc, cytotoxic chemotherapeutic drugs and molecular-targeted agents used in current lung cancer therapy (right) in the JFCR39 cancer cell panel. GI_50_ values of cisplatin, carboplatin, oxaliplatin, gemcitabine, PTX, docetaxel, DOX, MEL and ETP were cited from the literature.^38,39^ n.d. = no data.

**Extended data Figure 4.**
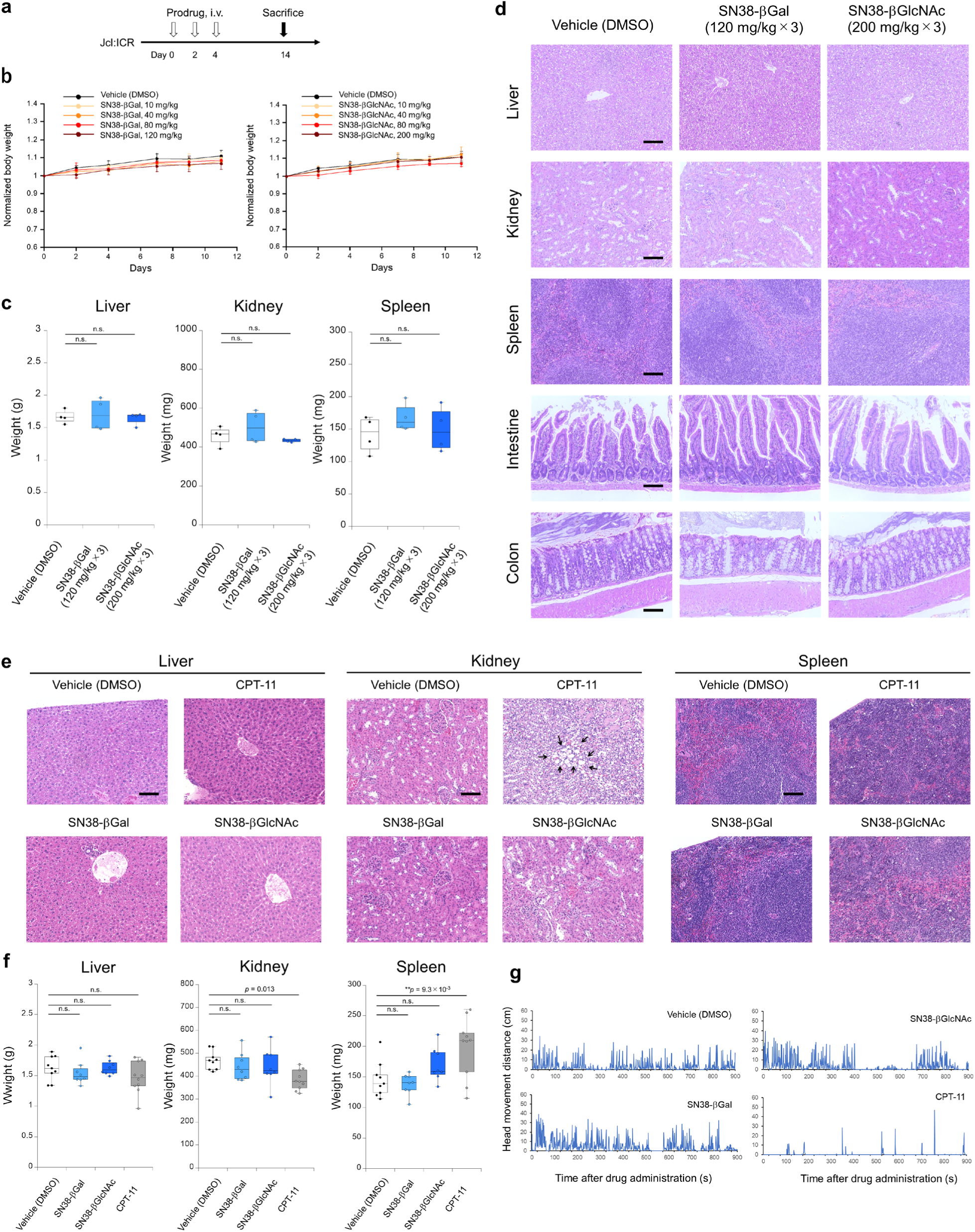
*In vivo* systemic toxicity studies of SN38-βGal and SN38-βGlcNAc in mice. (**a**) Representation of the MTD study of SN38-βGal and SN38-βGlcNAc. (**b**) Tolerability of various doses of SN38-βGal and SN38-βGlcNAc in non–tumor-bearing female ICR mice after three intravenous administrations (n = 4 for each group). Error bars represent s.d. (**c**) Organ weight after the MTD study shown in (**b**). No significant weight changes of organs were observed in either the SN38-βGal-or the SN38-βGlcNAc-treated group. One-way ANOVA with Tukey’s multiple comparisons test was used to calculate exact *p* values. n.s. = not significant. Error bars represent s.d. (**d**) Histological analysis of resected organs after the MTD study shown in (**b**). No notable abnormalities in organs were observed in the SN38-βGal-or SN38-βGlcNAc-treated group. Scale bars, 100 µm. (**e**) Histological analysis of resected organs after the repeated dose toxicity study shown in **Fig. 4a**. No notable abnormalities in organs were observed in the SN38-βGal-or SN38-βGlcNAc-treated group. Scale bars, 100 µm. (**f**) Organ weight after the repeated dose toxicity study shown in **Fig 4a**. No significant weight changes of liver, kidney and spleen were observed in the SN38-βGal-or SN38-βGlcNAc-treated group. A significant increase of spleen weight and a significant decrease of kidney weight were observed in the CPT-11-treated group. One-way ANOVA with Tukey’s multiple comparisons test was used to calculate exact *p* values. n.s. = not significant. Error bars represent s.d. (**g**) Locomotion evaluation of mice after intravenous administration of 40 mg/kg of each drug shown in **Fig. 4c-d**. Head movement distance was observed in each time interval. A significant locomotion decrease was observed in the CPT-11-treated group.

**Extended data Figure 5.**
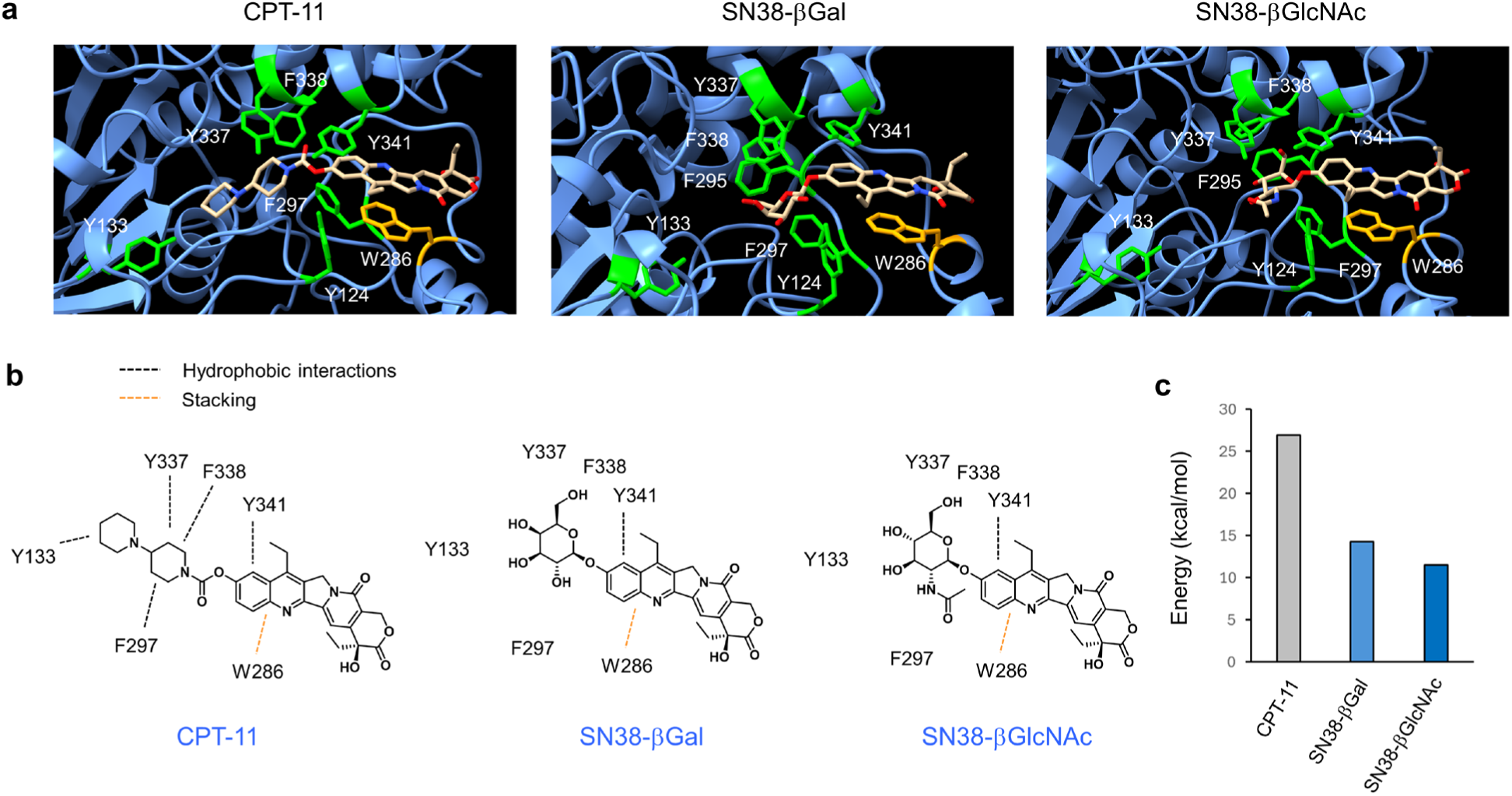
QM/MM calculation to predict the mechanism of low affinity of SN38-βGal and SN38-βGlcNAc for AChE. (**a**) Docking images of SN38-βGal, SN38-βGlcNAc and CPT-11 with hAChE obtained from QM/MM calculations. (**b**) Predicted key amino residues of hAChE for the interaction of prodrugs. Black dotted lines indicated predicted hydrophobic interactions. Orange dotted lines indicated predicted stacking interactions. CPT-11 exhibits strong hydrophobic interactions with the hAChE hydrophobic pocket composed of Y133, F297, Y337 and F338, whereas the hydrophilic sugar moieties in the developed prodrugs reduce this affinity. (**c**) QM/MM-calculated energy levels (kcal/mol) of CPT-11, SN38-βGal and SN38-βGlcNAc. QM/MM calculations suggested that the direct conjugation of the hydrophilic substrate to the 7-hydroxyl group of SN38 decreases the interaction with the hydrophobic pocket of AChE. These findings provide important insights for the general design of SN38-based prodrugs.

**Extended data Figure 6.**
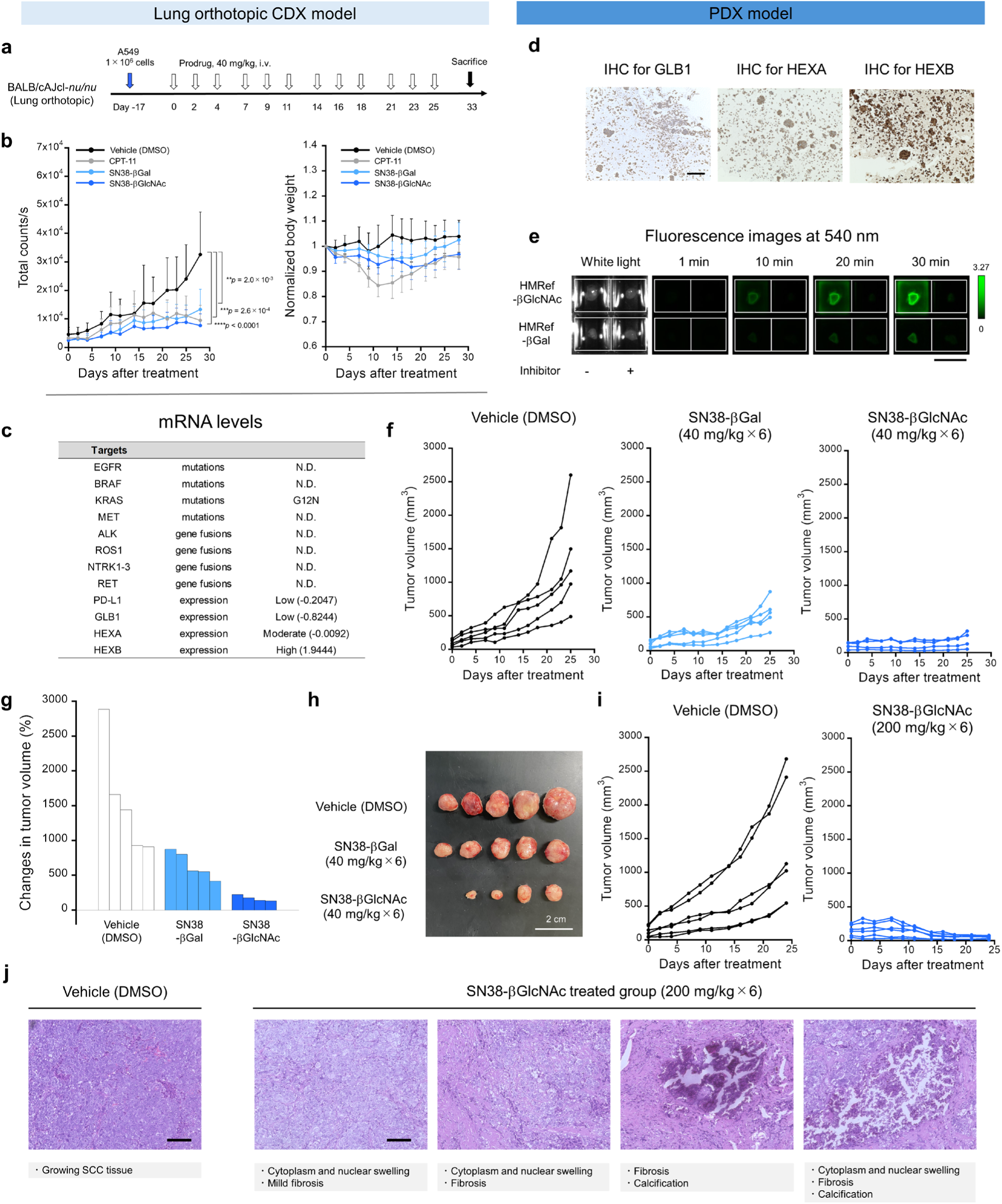
*In vivo* therapeutic efficacy of SN38-βGal and SN38-βGlcNAc in lung orthotopic CDX and lung SCC subcutaneous PDX models. (**a**) Representation of the prodrug treatment in lung orthotopic A549 tumor-bearing mice. (**b**) Anti-tumor efficacy (left) and weight change (right) in lung orthotopic A549 tumor-bearing mice following treatment with vehicle (n = 5), SN38-βGal (40 mg/kg × 12, n = 6), SN38-βGlcNAc (40 mg/kg × 12, n = 7) and CPT-11 (40 mg/kg × 12, n = 6). Tumor growth inhibition was quantified using a luciferin–luciferase bioluminescence assay. One-way ANOVA with Tukey’s multiple comparisons test was used to calculate exact *p* values. Error bars represent s.d. (**c**) mRNA expression levels of therapeutic targets for current lung cancer therapy and biomarker glycosidases discovered in the evaluated PDX tumor in **Fig. 5i-o**. The tumor has a high expression level of HEXB. N.D. = not detected. (**d**) IHC analysis for GLB1, HEXA and HEXB of evaluated PDX tumor in **Fig. 5i-o**. HEXB especially showed strong staining. Scale bar, 200 µm. (**e**) White light (left) and time-dependent fluorescence images at 540 nm (right) of PDX tumor after administration of HMRef-βGal or HMRef-βGlcNAc with and without the corresponding glycosidase inhibitor. [HMRef-βGal] = 50 µM, [HMRef-βGlcNAc] = 50 µM, [N-(n-nonyl)deoxygalactonoijirimycin] = 500 µM for HMRef-βGal, [PUGNAc] = 500 µM for HMRef-βGlcNAc. Ex/Em = 465 nm/515 nm long pass. Scale bar, 1 cm. (**f**) Individual tumor growth from **Fig. 5j**. (**g**) Changes in tumor volume for each mouse shown in (**f**). (**h**) White light image of resected PDX tumors after treatment, from **Fig. 5j**. SN38-βGal-and SN38-βGlcNAc-treated groups exhibited the tumor growth inhibition. Scale bar, 2 cm. (**i**) Individual tumor growth, from **Fig. 5l**. (**j**) Histological analysis of resected PDX tumors after treatment (200 mg/kg × 6), from **Fig. 5l**. The SN38-βGlcNAc-treated group exhibited chemotherapeutic damage such as cytoplasmic and nuclear swelling, fibrosis and calcification. Scale bars, 100 µm.

**Extended data Figure 7.**
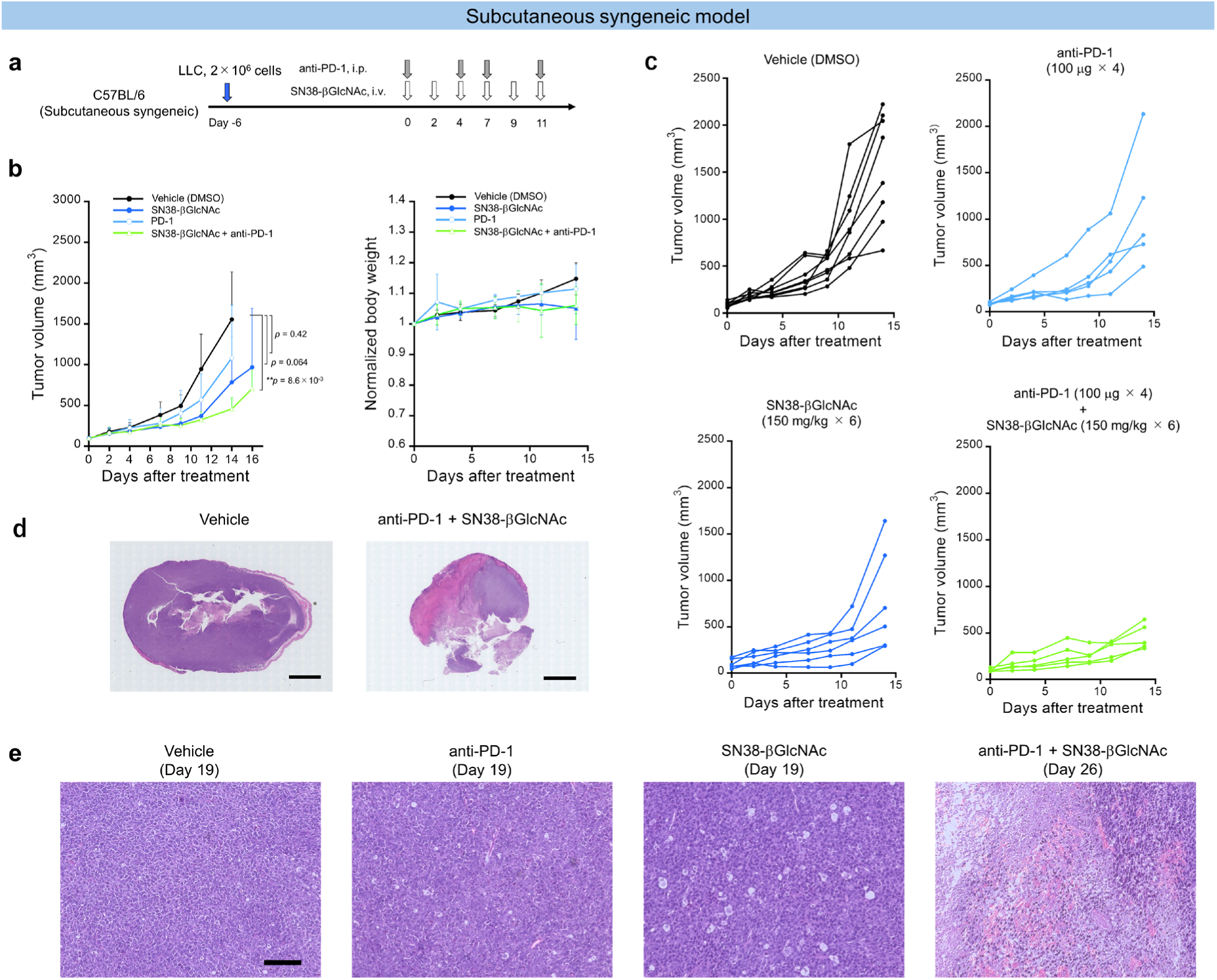
*In vivo* therapeutic efficacy of SN38-βGal and SN38-βGlcNAc in subcutaneous syngeneic models. (**a**) Representation of the drug treatment using subcutaneous LLC tumor-bearing mice. (**b**) Anti-tumor efficacy (left) and weight change (right) following the treatment of vehicle (n = 8), SN38-βGlcNAc (150 mg/kg × 6, n = 6), anti-PD-1 (100 µg × 4, n = 5) and the combination of SN38-βGlcNAc (150 mg/kg × 6) with anti-PD-1 (100 µg × 4) (n = 5). One-way ANOVA with Tukey’s multiple comparisons test was used to calculate exact *p* values. Error bars represent s.d. (**c**) Individual tumor growth from (**b**) following treatment with vehicle (n = 8), SN38-βGlcNAc (150 mg/kg × 6, n = 6), anti-PD-1 (100 µg × 4, n = 5) and the combination of SN38-βGlcNAc (150 mg/kg × 6) with anti-PD-1 (100 µg × 4) (n = 5). (**d**) Representative HE-staining images of resected tumors after treatment shown in (**b**). Scale bars, 3 mm. (**e**) Histological analysis of resected LLC tumors after treatment shown in (**b**). The anti-PD-1-or SN38-βGlcNAc-treated group exhibited chemotherapeutic damage such as cytoplasmic and nuclear swelling. The anti-PD-1 + SN38-βGlcNAc-treated group exhibited cytoplasmic and nuclear swelling, fibrosis and necrosis. Scale bars, 100 µm.

**Extended data Figure 8.**
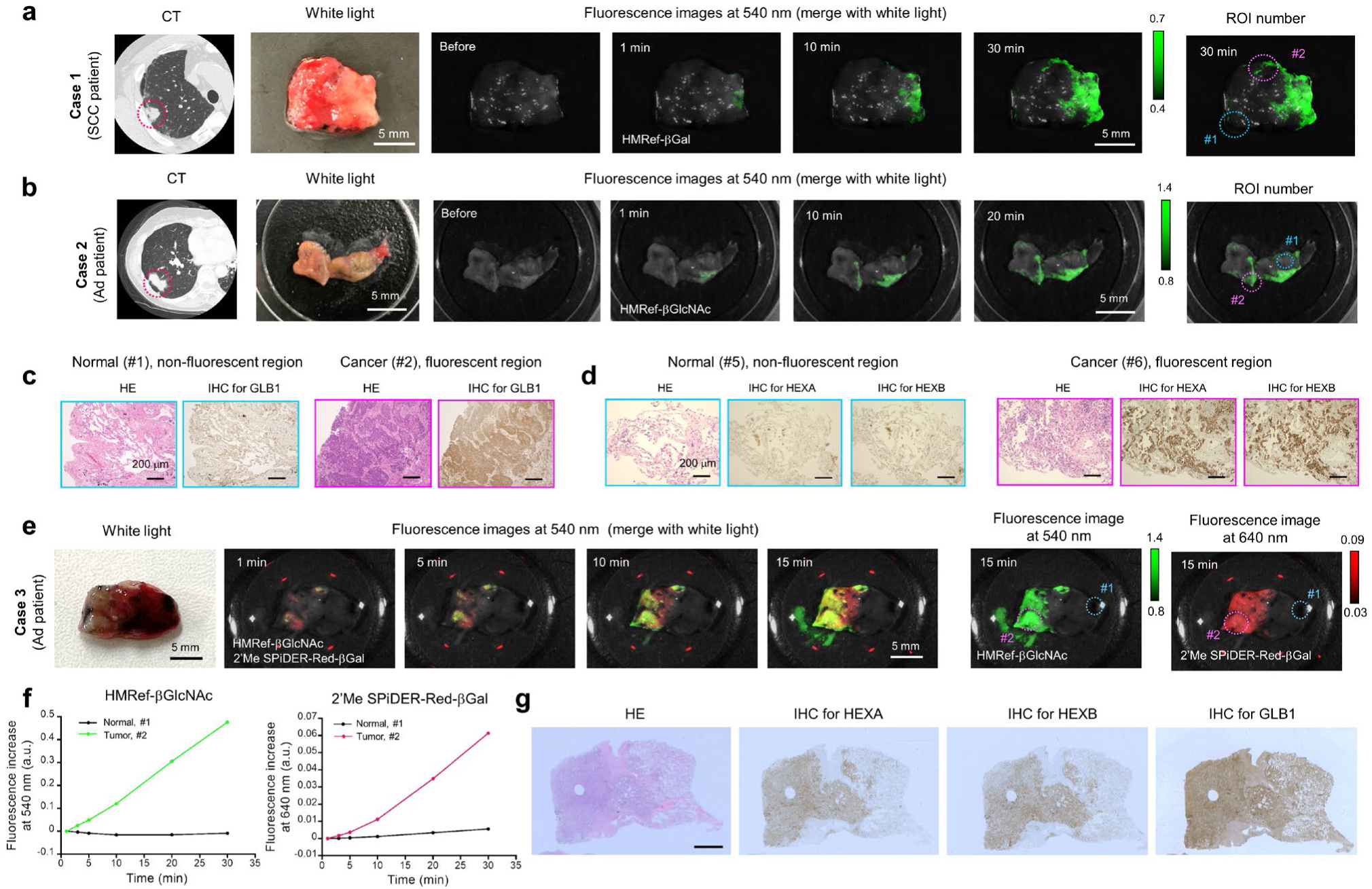
Activity-based fluorescence detection of biomarker enzyme activities in clinical lung cancer specimens. (**a**) CT image of SCC patient (left), corresponding white light image (middle) and time-dependent fluorescence images at 540 nm (right) of surgically resected fresh lung specimens containing both normal and SCC tissues after administration of HMRef-βGal (Case 1). Ex/Em = 465 nm/515 nm long pass. Scale bar, 5 mm. [HMRef-βGal] = 50 µM. (**b**) CT image of Ad patient (left), corresponding white light image (middle) and time-dependent fluorescence images at 540 nm of surgically resected fresh lung specimens containing both normal and Ad tissues after administration of HMRef-βGlcNAc (Case 2). Ex/Em = 465 nm/515 nm long pass. Scale bar, 5 mm. [HMRef-βGlcNAc] = 50 µM. (**c**) Histological analysis and IHC analysis for GLB1 of boxed regions with no fluorescence activation (blue box, #1) or strong fluorescence activation (pink box, #2) in (**a**). Fluorescence regions were well matched with pathologically SCC regions and GLB1-overexpressing regions. Scale bar, 200 µm. (**d**) Histological analysis and IHC analysis for HEXA and HEXB of boxed regions with no fluorescence activation (blue box, #1) or strong fluorescence activation (pink box, #2) in (**b**). Fluorescence regions were well matched with pathologically identified Ad regions and HEXA and HEXB-overexpressing regions. Scale bar, 200 µm. (**e**) White light image (left) and time-dependent fluorescence images at 540 nm and 640 nm (right) of surgically resected fresh lung specimens containing both normal and Ad tissues after administration of HMRef-βGlcNAc and 2’Me-4CH_2_F-Sirhodol-βGal (2’Me-SPiDER-Red-βGal) (Case 3). When a cocktail of red-emitting GLB1-reactive 2’Me-SPiDER-Red-βGal and green-emitting HMRef-βGlcNAc was used, the activities of both enzymes could be simultaneously detected in the same clinical specimen. Ex/Em = 465 nm/515 nm long pass for green, Ex/Em = 570 nm/610 nm long pass for red. Scale bar, 5 mm. [HMRef-βGlcNAc] = 50 µM, [2’Me-SPiDER-Red-βGal] = 50 µM. (**f**) Time-dependent fluorescence increase of HMRef-βGlcNAc and 2’Me-SPiDER-Red-βGal in boxed regions with no fluorescence activation (blue box, #1) or strong fluorescence activation (pink box, #2) in (**e**). (**g**) Histological analysis and IHC analysis for HEXA, HEXB and GLB1 of the case 3 specimen in (**e**). H.E staining indicated that the fluorescence-activated regions coincided well with pathologically diagnosed cancer cell-containing regions, while non-fluorescent regions were diagnosed as normal lung tissue. IHC staining confirmed that the pathologically diagnosed cancer regions exhibited higher target expression levels than normal regions. Scale bar, 2 mm. The evaluated fluorescence probes should be useful for the sensitive detection of elevated GLB1 and HEX activities in surgical specimens.

---

**Extended data Table 1.**
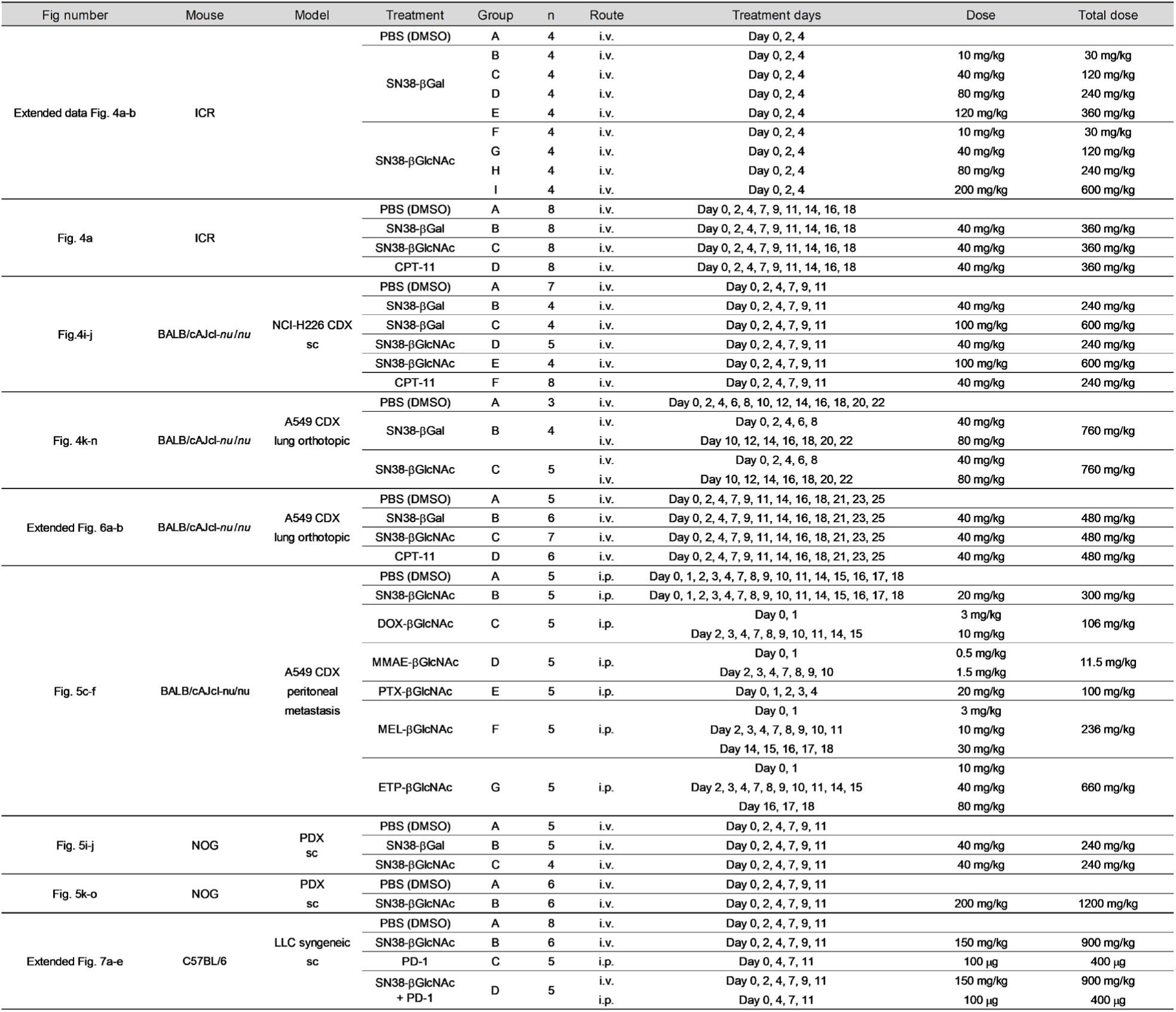
Summary of treatment data.

**Extended data Table 2.** Pharmacokinetic parameters of developed prodrugs and release of SN38 in mouse and common marmoset.

|  | Prodrugs | Dose | *t*<sub>1/2</sub> (h) | S.D. | AUC<sub>0-1h</sub> (µg · h/mL) | S.D. | AUC<sub>0-∞</sub> (µg · h/mL) | S.D. | CL<sub>tot</sub> (mL/min/kg) | S.D. | V<sub>d</sub> (L/kg) | S.D. |
| --- | --- | --- | --- | --- | --- | --- | --- | --- | --- | --- | --- | --- |
| Mouse | SN38-βGal | 40 mg/kg | 0.165 |  | 23.91 |  | 24.206 |  | 27.5 |  | 0.344 |  |
|  | SN38 |  | 1.255 |  | 0.173 |  | 0.454 |  |  |  |  |  |
|  | SN38-βGlcNAc | 40 mg/kg | 0.149 |  | 8.92 |  | 8.97 |  | 74.3 |  | 0.621 |  |
|  | SN38 |  | 0.203 |  | 1.020 |  | 1.055 |  |  |  |  |  |
|  | Prodrugs | Dose | *t*<sub>1/2</sub> (h) | S.D. | AUC<sub>0-6h</sub> (µg · h/mL) | S.D. | AUC<sub>0-∞</sub> (µg · h/mL) | S.D. | CL<sub>tot</sub> (mL/min/kg) | S.D. | V<sub>d</sub> (L/kg) | S.D. |
| Common marmoset | SN38-βGal | 10 mg/kg | 0.972 | 0.137 | 52.73 | 20.65 | 53.27 | 20.96 | 3.5 | 1.3 | 0.215 | 0.052 |
|  | SN38 |  | 1.419 | 0.355 | 0.820 | 0.270 | 0.84 | 0.26 |  |  |  |  |
|  | SN38-βGlcNAc | 10 mg/kg | 0.794 | 0.054 | 78.26 | 5.54 | 78.64 | 5.64 | 2.2 | 0.2 | 0.142 | 0.008 |
|  | SN38 |  | 1.557 | 0.051 | 0.909 | 0.202 | 0.952 | 0.207 |  |  |  |  |

Mouse: Values for dose of 40 mg/kg, bolus injection, n=3 for each time point. Common marmoset : Values for dose of 10 mg/kg, 11 min infusion, n=3.

## Notes

### Competing Interest Statement

The authors have declared no competing interest.

## References

1 Min, H.-Y. & Lee, H.-Y. Molecular targeted therapy for anticancer treatment. Experimental & Molecular Medicine 54, 1670–1694 (2022). 10.1038/s12276-022-00864-3

2 Waldman, A. D., Fritz, J. M. & Lenardo, M. J. A guide to cancer immunotherapy: from T cell basic science to clinical practice. Nature Reviews Immunology 20, 651–668 (2020). 10.1038/s41577-020-0306-5

3 Araghi, M. et al. Recent advances in non-small cell lung cancer targeted therapy; an update review. Cancer Cell International 23, 162 (2023). 10.1186/s12935-023-02990-y

4 Huang, Q. et al. Advances in molecular pathology and therapy of non-small cell lung cancer. Signal Transduction and Targeted Therapy 10, 186 (2025). 10.1038/s41392-025-02243-6

5 Marquart, J., Chen, E. Y. & Prasad, V. Estimation of the Percentage of US Patients With Cancer Who Benefit From Genome-Driven Oncology. JAMA Oncology 4, 1093–1098 (2018). 10.1001/jamaoncol.2018.1660

6 Haslam, A. & Prasad, V. Estimation of the Percentage of US Patients With Cancer Who Are Eligible for and Respond to Checkpoint Inhibitor Immunotherapy Drugs. JAMA Network Open 2, e192535–e192535 (2019). 10.1001/jamanetworkopen.2019.2535

7 Anand, U. et al. Cancer chemotherapy and beyond: Current status, drug candidates, associated risks and progress in targeted therapeutics. Genes & Diseases 10, 1367–1401 (2023). 10.1016/j.gendis.2022.02.007

8 Oun, R., Moussa, Y. E. & Wheate, N. J. The side effects of platinum-based chemotherapy drugs: a review for chemists. Dalton Transactions 47, 6645–6653 (2018). 10.1039/C8DT00838H

9 van den Boogaard, W. M. C., Komninos, D. S. J. & Vermeij, W. P. Chemotherapy Side-Effects: Not All DNA Damage Is Equal. Cancers 14, 627 (2022).

10 Lai, J.-I., Chao, T.-C., Liu, C.-Y., Huang, C.-C. & Tseng, L.-M. A systemic review of taxanes and their side effects in metastatic breast cancer. Frontiers in Oncology Volume 12–2022 (2022). 10.3389/fonc.2022.940239

11 Fujita, K. & Urano, Y. Activity-Based Fluorescence Diagnostics for Cancer. Chemical Reviews 124, 4021–4078 (2024). 10.1021/acs.chemrev.3c00612

12 Sun, I.-C., Yoon, H. Y., Lim, D.-K. & Kim, K. Recent Trends in In Situ Enzyme-Activatable Prodrugs for Targeted Cancer Therapy. Bioconjugate Chemistry 31, 1012–1024 (2020). 10.1021/acs.bioconjchem.0c00082

13 Martin, H., Lázaro, L. R., Gunnlaugsson, T. & Scanlan, E. M. Glycosidase activated prodrugs for targeted cancer therapy. Chemical Society Reviews 51, 9694–9716 (2022). 10.1039/D2CS00379A

14 Han, H.-H. et al. The design of small-molecule prodrugs and activatable phototherapeutics for cancer therapy. Chemical Society Reviews 52, 879–920 (2023). 10.1039/D2CS00673A

15 Kuriki, Y. et al. Development of a fluorescent probe library enabling efficient screening of tumour-imaging probes based on discovery of biomarker enzymatic activities. Chemical Science 13, 4474–4481 (2022). 10.1039/D1SC06889J

16 Kawashima, S. et al. Rapid imaging of lung cancer using a red fluorescent probe to detect dipeptidyl peptidase 4 and puromycin-sensitive aminopeptidase activities. Scientific Reports 12, 9100 (2022). 10.1038/s41598-022-12665-9

17 Fujita, K. et al. Rapid and Accurate Visualization of Breast Tumors with a Fluorescent Probe Targeting α-Mannosidase 2C1. ACS Central Science 6, 2217–2227 (2020). 10.1021/acscentsci.0c01189

18 Takahashi, R. et al. Real-Time Fluorescence Imaging to Identify Cholangiocarcinoma in the Extrahepatic Biliary Tree Using an Enzyme-Activatable Probe. Liver Cancer, 1–13 (2023). 10.1159/000530645

19 Kobayashi, K. et al. Rapid imaging of pulmonary metastasis from colorectal cancer with a red fluorescence probe targeting puromycin-sensitive aminopeptidase and dipeptidyl peptidase IV. Scientific Reports 15, 43930 (2025). 10.1038/s41598-025-27717-z

20 Fujita, K., Kamiya, M. & Urano, Y. Rapid and Sensitive Detection of Cancer Cells with Activatable Fluorescent Probes for Enzyme Activity. *Methods in molecular biology (Clifton*, N.J.) 2274, 193–206 (2021). 10.1007/978-1-0716-1258-3_17

21 Hino, H. et al. Rapid Cancer Fluorescence Imaging Using A γ-Glutamyltranspeptidase-Specific Probe For Primary Lung Cancer. Transl Oncol 9, 203–210 (2016). 10.1016/j.tranon.2016.03.007

22 Nakada, A., Maruyama, T., Kamiya, M., Hanaoka, K. & Urano, Y. Rapid Visualization of Deeply Located Tumors In Vivo by Intravenous Administration of a γ - Glutamyltranspeptidase-Activated Fluorescent Probe. Bioconjugate Chemistry 33, 523–529 (2022). 10.1021/acs.bioconjchem.2c00039

23 Tanaka, H. et al. Dipeptidylpeptidase-4-targeted activatable fluorescent probes visualize senescent cells. Cancer science 115, 2762–2773 (2024). 10.1111/cas.16229

24 Fujii, T., Kamiya, M. & Urano, Y. In Vivo Imaging of Intraperitoneally Disseminated Tumors in Model Mice by Using Activatable Fluorescent Small-Molecular Probes for Activity of Cathepsins. Bioconjugate Chemistry 25, 1838–1846 (2014). 10.1021/bc5003289

25 Onoyama, H. et al. Rapid and sensitive detection of early esophageal squamous cell carcinoma with fluorescence probe targeting dipeptidylpeptidase IV. Scientific Reports 6, 26399 (2016). 10.1038/srep26399

26 Brunetti-Pierri, N. & Scaglia, F. GM1 gangliosidosis: Review of clinical, molecular, and therapeutic aspects. Molecular Genetics and Metabolism 94, 391–396 (2008). 10.1016/j.ymgme.2008.04.012

27 Norflus, F., Yamanaka, S. & Proia, R. L. Promoters for the Human β-Hexosaminidase Genes, HEXA and HEXB. DNA and Cell Biology 15, 89–97 (1996). 10.1089/dna.1996.15.89

28 Gao, Y., Wells, L., Comer, F. I., Parker, G. J. & Hart, G. W. Dynamic O-Glycosylation of Nuclear and Cytosolic Proteins: CLONING AND CHARACTERIZATION OF A NEUTRAL, CYTOSOLIC β-N-ACETYLGLUCOSAMINIDASE FROM HUMAN BRAIN*. Journal of Biological Chemistry 276, 9838–9845 (2001). 10.1074/jbc.M010420200

29 Gutternigg, M., Rendić, D., Voglauer, R., Iskratsch, T. & Wilson, Iain B. H. Mammalian cells contain a second nucleocytoplasmic hexosaminidase. Biochemical Journal 419, 83–90 (2009). 10.1042/bj20081630

30 Komatsu, T. et al. Diced Electrophoresis Gel Assay for Screening Enzymes with Specified Activities. Journal of the American Chemical Society 135, 6002–6005 (2013). 10.1021/ja401792d

31 Venditto, V. J. & Simanek, E. E. Cancer Therapies Utilizing the Camptothecins: A Review of the in Vivo Literature. Molecular Pharmaceutics 7, 307–349 (2010). 10.1021/mp900243b

32 Yang, X.-Q., Li, C.-Y., Xu, M.-F., Zhao, H. & Wang, D. Comparison of first-line chemotherapy based on irinotecan or other drugs to treat non-small cell lung cancer in stage IIIB/IV: a systematic review and meta-analysis. BMC Cancer 15, 949 (2015). 10.1186/s12885-015-1978-2

33 Fujita, K., Kubota, Y., Ishida, H. & Sasaki, Y. Irinotecan, a key chemotherapeutic drug for metastatic colorectal cancer. World journal of gastroenterology 21, 12234–12248 (2015). 10.3748/wjg.v21.i43.12234

34 Wang-Gillam, A. et al. NAPOLI-1 phase 3 study of liposomal irinotecan in metastatic pancreatic cancer: Final overall survival analysis and characteristics of long-term survivors. European Journal of Cancer 108, 78–87 (2019). 10.1016/j.ejca.2018.12.007

35 Kweekel, D., Guchelaar, H.-J. & Gelderblom, H. Clinical and pharmacogenetic factors associated with irinotecan toxicity. Cancer Treatment Reviews 34, 656–669 (2008). 10.1016/j.ctrv.2008.05.002

36 Campbell, J. M. et al. Irinotecan-induced toxicity pharmacogenetics: an umbrella review of systematic reviews and meta-analyses. The Pharmacogenomics Journal 17, 21–28 (2017). 10.1038/tpj.2016.58

37 Shoemaker, R. H. The NCI60 human tumour cell line anticancer drug screen. Nat Rev Cancer 6, 813–823 (2006). 10.1038/nrc1951

38 Dan, S. et al. An integrated database of chemosensitivity to 55 anticancer drugs and gene expression profiles of 39 human cancer cell lines. Cancer research 62, 1139–1147 (2002).

39 Dan, S. et al. Correlating phosphatidylinositol 3-kinase inhibitor efficacy with signaling pathway status: in silico and biological evaluations. Cancer research 70, 4982–4994 (2010). 10.1158/0008-5472.Can-09-4172

40 Gandia, D. et al. CPT-11-induced cholinergic effects in cancer patients. Journal of Clinical Oncology 11, 196–197 (1993). 10.1200/jco.1993.11.1.196

41 Taha, T. et al. Treatment of Rare Mutations in Patients with Lung Cancer. Biomedicines 9, 534 (2021).

42 Yu, H., Boyle, T. A., Zhou, C., Rimm, D. L. & Hirsch, F. R. PD-L1 Expression in Lung Cancer. Journal of Thoracic Oncology 11, 964–975 (2016). 10.1016/j.jtho.2016.04.014

43 Chevallier, M., Borgeaud, M., Addeo, A. & Friedlaender, A. Oncogenic driver mutations in non-small cell lung cancer: Past, present and future. World J Clin Oncol 12, 217–237 (2021). 10.5306/wjco.v12.i4.217

44 Wagner, J. et al. Overexpression of the Novel Senescence Marker β-Galactosidase (GLB1) in Prostate Cancer Predicts Reduced PSA Recurrence. PLOS ONE 10, e0124366 (2015). 10.1371/journal.pone.0124366

45 Prabha, M., Swamy, N. & Ravi, V. Specific activity of glycosidases in brain tumors and their expression in primary explants culture. Journal of Biochemical Technology 5, 654–665 (2013).

46 Veninga, V. & Voest, E. E. Tumor organoids: Opportunities and challenges to guide precision medicine. Cancer Cell 39, 1190–1201 (2021). 10.1016/j.ccell.2021.07.020

47 Ogawa, A. et al. SLFN11-mediated ribosome biogenesis impairment induces TP53-independent apoptosis. Molecular Cell 85, 894–912.e810 (2025). 10.1016/j.molcel.2025.01.008

48 Takeuchi, K. et al. Prediction of cell cycle distribution after drug exposure by high content imaging analysis using low-toxic DNA staining dye. Pharmacology Research & Perspectives 12, e1203 (2024). 10.1002/prp2.1203

49 Skehan, P. et al. New Colorimetric Cytotoxicity Assay for Anticancer-Drug Screening. JNCI: Journal of the National Cancer Institute 82, 1107–1112 (1990). 10.1093/jnci/82.13.1107

50 Kudara, M., Kato-Ishikura, E., Ikegaya, Y. & Matsumoto, N. Ramelteon administration enhances novel object recognition and spatial working memory in mice. Journal of Pharmacological Sciences 152, 128–135 (2023). 10.1016/j.jphs.2023.04.002

