## Supplementary Information for "Enzyme activity as an actionable axis for small-molecule precision oncology"

[1]Graduate School of Medicine, [2]Graduate School of Pharmaceutical Sciences, [3]Department of Thoracic Surgery and [4]Institute for AI and Beyond, The University of Tokyo, 7-3-1 Hongo, Bunkyo-ku, Tokyo 113-0033, Japan. [5]ACT-X, Japan Science and Technology Agency, 4-1-8 Honcho, Kawaguchi, Saitama 332-0012, Japan. [6]Division of Molecular Pharmacology, Cancer Chemotherapy Center, Japanese Foundation for Cancer Research, 3-8-31 Ariake, Koto-ku, Tokyo 135-8550, Japan. [7]Laboratory for Chemistry and Life Science, Institute of Integrated Research, Institute of Science Tokyo, Yokohama, Kanagawa 226-8501, Japan. [8]Department of Sports and Health Science, Daito Bunka University, 560 Iwadono, Higashimatsuyama, Saitama 355-8501, Japan. [9]DMPK Research Laboratory, Watarase Research Center, Discovery Research Headquarters, Kyorin Pharmaceutical Co., Ltd, 1848, Nogi, Nogi-machi, Shimotsuga-gun, Tochigi, 329-0114, Japan. [10]Center for Information and Neural Networks, National Institute of Information and Communications Technology, Suita, Osaka, 565-0871, Japan.

*

### Table of contents

**Experimental methods** • • • • • • • • • • • • • • • • • • • • • • • • • • • • • • • • • • • • • • • **4-8**

**Figure S1. Representative fluorescence probes used in this study** • • • • • • • • • • • • • • • • • • • • **9**

**Table S1. Selected probes for activity-based screening used in this study** • • • • • • • • • • • • • • • • • • • • • • • • • • • • • • • • • • • • • • • • • • **10**

**Table S2. Organ weights used to determine background scores** • • • • • • • • • • • • • • • • • • • • • **10**

**Table S3. Definition and nomenclature of background score notation** • • • • • • • • • • • • • • • • • **11**

**Figure S2. Representative time-dependent fluorescence images in the screening using surgically resected normal lung and lung cancer tissues** • • • • • • • • • • • • • • • • • • • • • • • • • • • • • • • • **12**

**Figure S3. ROC curves of HMRef-β-D-Glc and HMRef-α-L-Fuc for lung cancer detection** • • • • • • • **13**

**Figure S4. Fluorescence images of resected mouse organs after systemic administration of each fluorescent probe** • • • • • • • • • • • • • • • • • • • • • • • • • • • • • • • • • • • • • • • • **14**

**Figure S5. Fluorescence imaging of glycosidase activities using HMRef-βGal and HMRef-βGlcNAc in lung cancer cell lines** • • • • • • • • • • • • • • • • • • • • • • • • • • • • • • • • • • • • • • • • • **15**

**Figure S6. DEG assay using HMRef-βGlcNAc with surgically resected Ad specimen from human lung** • • • • • • • • • • • • • • • • • • • • • • • • • • • • • • • • • • • • • • • • • • • • • • • **16**

**Table S4. List of detected proteins in DEG assay of a human lung Ad surgical specimen** • • • • • **17-19**

**Figure S7. IHC analysis for GLB1 in lung normal, Ad and SCC tissues** • • • • • • • • • • • • • • • • **20**

**Figure S8. IHC analysis for HEXA and HEXB in lung normal, Ad and SCC tissues** • • • • • • • • • • **21**

**Figure S9. Evaluation of reactivity of SN38-βGal and SN38-βGlcNAc with target enzymes by LC-MS** • • • • • • • • • • • • • • • • • • • • • • • • • • • • • • • • • • • • • • • • • • • • • • • **22**

**Figure S10. Evaluation of enzyme activities in cultured cancer cells by live cell imaging** • • • • • • **22**

**Figure S11. Time-dependent fluorescence images of A549 cells in the presence of SN38 or CPT-11** • • **23**

**Figure S12. Time-dependent fluorescence images of A549 cells in the presence of SN38-βGal or SN38-βGlcNAc** • • • • • • • • • • • • • • • • • • • • • • • • • • • • • • • • • • • • • • • • • • **24**

**Figure S13. Time-dependent fluorescence images of A549 cells in the presence of SN38-βGlcNAc** • • **25**

**Figure S14. Comprehensive evaluation of concentration-dependent cell growth inhibition in JFCR39 cancer cell panel** • • • • • • • • • • • • • • • • • • • • • • • • • • • • • • • • • • • • • • • **26**

**Figure S15. Correlations between enzymatic activities or anticancer efficacy and produced SN38 for the developed prodrugs in the JFCR39 cancer cell panel** • • • • • • • • • • • • • • • • • • • • • • • **26**

**Table S5. Biochemical analysis in the MTD study** • • • • • • • • • • • • • • • • • • • • • • • • • • **27**

**Table S6. Biochemical analysis in the repeated-dose toxicity study • • • • • • • • • • • • • • • • • • • 27**

**Figure S16. Evaluation of the head movement trajectory in mice during the 15 min following intravenous injection of each drug • • • • • • • • • • • • • • • • • • • • • • • • • • • • • • • • • • • • 28**

**Table S7. Evaluation of $LD_{50}$ values of SN38-βGal and SN38-βGlcNAc • • • • • • • • • • • • • • • • • 29**

**Figure S17. Plasma concentration changes of developed prodrugs (logarithmic scale) • • • • • • • 30**

**Table S8. Pharmacokinetic parameters of SN38-βGal, SN38-βGlcNAc and released SN38 in common marmosets • • • • • • • • • • • • • • • • • • • • • • • • • • • • • • • • • • • • • • • • 31**

**Figure S18. Histological analysis of resected PDX tumors after treatment with SN38-βGal or SN38-βGlcNAc • • • • • • • • • • • • • • • • • • • • • • • • • • • • • • • • • • • • • • • • • 32**

**Figure S19. Histological and IHC analysis for GLB1 in the clinical specimen evaluated by GLB1-reactive fluorescence probe • • • • • • • • • • • • • • • • • • • • • • • • • • • • • • • • • • • 32**

**Figure S20. Histological and IHC analysis for HEXA and HEXB in the clinical specimen evaluated by HEX-reactive fluorescence probe • • • • • • • • • • • • • • • • • • • • • • • • • • • • • • • • • 33**

**Figure S21. Rapid and sensitive visualization of target glycosidase activities in clinical lung cancer specimens using fluorescence probes • • • • • • • • • • • • • • • • • • • • • • • • • • • • • 34**

**Figure S22. Reaction scheme for 2'Me SPiDER-Red-βGal • • • • • • • • • • • • • • • • • • • • • • 35**

**Figure S23. Correlations between -log $GI_{50}$ of developed prodrugs and SLFN11 expression • • • • • • • • • • • • • • • • • • • • • • • • • • • • • • • • • • • • • • • • • • 36**

**Organic synthesis and characterization of compounds • • • • • • • • • • • • • • • • • • • • • • 37-50**

**References • • • • • • • • • • • • • • • • • • • • • • • • • • • • • • • • • • • • • • • • • • 51-52**

**Reagents**. All organic solvents and reagents were commercial products of guaranteed grade, and were used without further purification. Water was doubly distilled and deionized by a Milli-Q water system before use.

**Cell lines and culture conditions.** Established cell lines, A549, A549-Luc, NCI-H226, SHP-77, HEK/lacZ and LLC were used in this study. NCI-H226 SHP-77 and LLC were cultured in RPMI-1640 (with L-glutamine and phenol red) and maintained at 37°C in a humidified incubator under 5% $CO_2$ in air. A549, A549-Luc and HEK/lacZ were cultured in DMEM (high glucose with L-glutamine and phenol red) and maintained under the same conditions. All media were supplemented with 10% fetal bovine serum and 1% penicillin/streptomycin. A549 was obtained from RIKEN Cell Bank. NCI-H226 was obtained from American Type Culture Collection (ATCC). A549-Luc and LLC were obtained from Japanese Collection of Research Bioresources (JCRB) Cell Bank. SHP-77 was obtained from European Collection of Authenticated Cell Cultures (ECACC). HEK/lacZ was obtained from InvivoGen.

**Evaluation of stability of fluorescence probes in human serum.** Fluorescence probes (1 μM) were incubated with 10% human serum (Human serum, Pooled, Product number: 2930149, Lot number: U1123080905, MP Biomedicals) (*v/v*) containing PBS (-) solution in 384-well black plate (4514, Corning) at 37 °C. Fluorescence intensity was measured at 1 h, 2 h, 3 h and 6 h using an Envision plate reader (PerkinElmer). Filter settings were FITC 485 (485/14 nm) for excitation and FITC 535 (535/25 nm) for emission.

***Ex-vivo* fluorescence imaging of enzyme activities in resected CDX and PDX tumors**. Tumors (~1000 $cm^3$) were resected from subcutaneous A549 tumor or H226 tumor-bearing BALB/cAJcl-*nu*/*nu* mice or PDX tumor-bearing NOD.Cg-Prkdc<scid＞Il2rg<tmlSug>/ShiJic (NOG) mice. Resected tumors were cut into pieces a few mm in size. A 50 μM solution of fluorescence probe (200 μL) in PBS (-) with or without 500 μM enzyme inhibitor was added to an 8-well chamber (μ-Slide 8 well; Ibidi) containing tumor tissues so that the tissues were completely soaked with probe solution. The fluorescence images of the tumors were evaluated using the Maestro *in-vivo* imaging system (CRi Inc.) and recorded. Exposure time was set at 100-50 msec depending on the level of fluorescence intensity from specimens. The stage and lamp were both set at position 1 or 2.

**Evaluation of reactivity of fluorescence probes with recombinant proteins.** PBS (-) solutions containing 1 μM HMRef-βGal and 10 ng/mL GLB1 (6464-GH-020, R and D Systems) with or without 10 μM N-(n-nonyl)deoxygalactonoijirimycin were incubated for 1 h at 37 °C. PBS (-) solutions containing 1 μM HMRef-βGlcNAc and 10 ng/mL HEXA (637-GH-020, R and D Systems) or HEXB (8907-GH-020, R and D Systems) with or without 10 μM PUGNAc were incubated for 1 h at 37 °C. Fluorescence intensity was measured over 1 h using an Envision plate reader (PerkinElmer). Filter settings were FITC 485 (485/14 nm) for excitation and FITC 535 (535/25 nm) for emission.

**Immunohistochemical analysis of target enzyme expression levels.** For the analysis of GLB1 expression, sections were deparaffinized in Histo-Clear, sequentially washed in 100%, 90%, 80% and 70% ethanol, and then washed in PBS (-). After heat-induced antigen retrieval (citrate buffer, pH 6) using microwave irradiation, each slide was pre-incubated in 3% $H_2O_2$ for 20 min, reacted with primary antibodies in 5% skim milk for 90 min, and then with secondary antibodies (TaKaRa POD Conjugate Anti Rabbit, For Tissue, Product number: MK205, Lot: AJ92256A, TaKaRa) for 30 min at room temperature. Each slide was visualized with a 3,3'-diaminobenzidine tetrahydrochloride (DAB) detection kit (Product number: MK210, TaKaRa), and counterstained with hematoxylin. Anti-GLB1 (rabbit polyclonal, Product number: HPA069503, Lot: A114940, Sigma-Aldrich) was used as the primary antibody after 1/50 dilution. DAB reaction time was 5 min. For the analysis of HEXA and HEXB expression levels, immunoperoxidase staining for anti-HEXA (rabbit polyclonal, Product number: HPA054583, Lot: A104822, SIGMA-ALDRICH) and anti-HEXB (rabbit polyclonal, Product number: HPA055409, Lot: R74092, SIGMA-ALDRICH) was performed using a Ventana Benchmark XT (Ventana Medical Systems) automated slide staining system. Sections were deparaffinized, pretreated with Cell Conditioning 1 (CC1, Ventana Medicals Systems), reacted with the primary antibody for 32 min at room temperature, visualized by Ventana's iView DAB detection kit and counterstained with hematoxylin and bluing reagent (Ventana Medical Systems). HEXA and HEXB antibodies were diluted to 1/20, and used with the iVIEW DAB detection kit (Ventana Medical Systems) and endogenous biotin blocking kit (Ventana Medical Systems). Human liver tissues were used as a positive control of HEXA immunostaining, and for the negative control, REAL Antibody Diluent (Dako) was added instead of the primary antibody.

**Evaluation of reactivity of SN38-βGal and SN38-βGlcNAc with target enzymes.** A 100 mM $CH_3COOH/CH_3COONa$ buffer (pH 5.5) containing 40 μM prodrug and recombinant target enzyme was incubated for 10 min, 30 min or 1 h at 37 °C. MeOH (80 μL) was added to this solution (20 μL) to quench the enzyme reaction. The samples were analyzed by LC-MS (Aquity UPLC H-Class Plus, Waters). GLB1 (200 μg/mL) (6464-GH-020, R and D Systems) was added to SN38-βGal. HEXA (66.7 μg/mL) (637-GH-020, R and D Systems) or HEXB (20 μg/mL) (8907-GH-020, R and D Systems) was added to SN38-βGlcNAc.

**Evaluation of SN38 production rates in SHP-77 and HEK/lacZ cells.** SHP-77 or HEK/lacZ ($5.0 \times 10^5$ cells) in 100 μL/well of OptiMEM (Thermo Fisher Scientific) were seeded in a 96-well plate (3915, Costar) SN38-βGal or SN38-βGlcNAc in 100 μL of OptiMEM was added to each well at a final concentration of 20 μM, and the plate was incubated at 37°C under 5% $CO_2$ in humidified air for 24 h. After incubation, 50 μL/well of medium was collected and mixed with 50 μL of MeOH/MeCN (1:1, *v*/*v*) containing 40 μM camptothecin as an internal standard. The sample was filtered by centrifugation at 12,000 × g for 5 min at 4°C using a Cosmo Spin Filter G (06549-44, Nacalai Tesque). The filtrate (50 μL) was diluted with Milli-Q water (50 μL), and the SN38 production rate was quantified by LC-MS (Aquity UPLC H-Class Plus, Waters).

**JFCR39 cell lines.** JFCR39 cell lines were maintained at the Japanese Foundation for Cancer Research. Cells were cultured in RPMI-1640 (with L-glutamine and phenol red) containing 5% fetal bovine serum and maintained at 37°C in a humidified incubator under 5% $CO_2$ in air.

**Fluorescence evaluation of enzyme activities in the JFCR39 cancer cell panel.** Cells (7,500 cells/well) and fluorescence probe (10 μM) in 25 μL of RPMI-1640 medium supplemented with 5% heat-inactivated human serum (H3667-100ML, Sigma-Aldrich) containing 0.005% CHAPS were placed in a 384-well black plate. The plate was incubated at 37°C in a humidified incubator under 5% $CO_2$ in air. After 3 h and 24 h incubation, fluorescence intensity was measured using Envision plate reader (PerkinElmer). Filter settings were FITC 485 (485/14 nm) for excitation and FITC 535 (535/25 nm) for emission. CVR of probes in each cell line were calculated by dividing the fluorescence intensity by that of the resulting HMRef fluorophore at the same measurement time.

**Evaluation of SN38 production rates in the JFCR39 cancer cell panel.** Cells ($6.0 \times 10^4$ cells) were seeded in a 96-well plate in 100 μL of RPMI-1640 medium supplemented with 5% heat-inactivated human serum (H3667-100ML, Sigma-Aldrich). SN38-βGal or SN38-βGlcNAc in 10 μL of medium was added to each well at a final concentration of 20 μM, and the plate was incubated at 37°C in a humidified incubator under 5% $CO_2$ in air for 24 h. Then, 40 μL of medium from each well was collected, mixed with 40 μL of MeOH/MeCN (1:1, *v*/*v*) and stored at -20°C until analysis. Subsequently, 80 μL of MeOH/MeCN (1:1, *v*/*v*) containing 40 μM camptothecin as an internal standard was added, and centrifuged at 1,700 × g for 15 min at 4°C. The supernatant was diluted with 50 μL of Milli-Q water, and the SN38 production rate was quantified by LC-MS (Aquity UPLC H-Class Plus, Waters).

**Maximum tolerated dose toxicity studies.** Female 7-week-old female Jcl:ICR mice with free access to food and water before experiments were intravenously administered SN38-βGal and SN38-βGlcNAc via the tail vein: 10, 40, 80, and 120 mg/kg for SN38-βGal, and 10, 40, 80 and 200 mg/kg for SN38-βGlcNAc, three times per week for one week (n = 4 for each group). Body weight changes were monitored for two weeks following the initial administration. On day 14, all mice were sacrificed, and the liver, kidney, spleen, intestine and colon were collected for pathological examination. The liver, kidney and spleen were weighed. Blood samples for biochemical analysis were collected from the inferior vena cava under medetomidine/midazolam/butorphanol anesthesia.

**Acute toxicity studies.** A single intravenous dose of SN38-βGal or SN38-βGlcNAc was administered via the tail vein to 7-week-old female Jcl:ICR mice. The dosing regimens were 200 and 300 mg/kg for SN38-βGal and 200 and 500 mg/kg for SN38-βGlcNAc (n = 10 for each group). Mortality was monitored over a 14-day period. The maximum administrable doses were 300 mg/kg for SN38-βGal and 500 mg/kg for SN38-βGlcNAc owing to solubility limitations. The dosing solutions contained DMSO as a cosolvent at volumes of less than 50 μL for SN38-βGal and less than 55 μL for SN38-βGlcNAc.

**QMM calculations.** To the crystal structure of hAChE (from *Homo sapiens*, PDB ID: 6O5R), protons at pH 7.4 were assigned to each amino acid using the PROPKA server.[1] The structure of each drug molecule (CPT-11 or SN38-sugar conjugates) was appended so that the binding mode is the same as in the cocrystal structure of CPT-11 and hAChE (from *Tetronarce californica*, PDB ID: 1U65). With this structure, MD sampling (GROMACS 2023.1.[2] (B) with the AMBER03 parameter set) was performed with fixed coordinates of the hAChE-drug conjugate. Sampling settings were determined according to the literature.[3] Solvent water molecules at distances greater than 15 Å from the drug were removed from the NPT ensemble. The hybrid structure (hAChE+drug+solvent) was optimized by the 2-layer ONIOM method with Gaussian16.[4] The calculations were performed at the level of ONIOM(APFD/6-311+g(2d,p):UFF)//ONIOM(B3LYP+D3/6-31G(d,p):UFF) with electronic embedding. The high layer included the drug molecule and the side chain of 286Trp which has a π-π interaction with the drug. For the low layer, the atomic charges of the drug were calculated at the same level as used for the high layer. In the structural optimization, only atoms within 10 Å of the high layer atoms (except the hydrogen atoms of water) were allowed to relax. The difference between the energy calculation results above and the sum of the results for the drug alone (APFD/6-311+g(2d,p), in water (iefpcm)) and hAChE without the drug was calculated as the binding stabilization energy.

**Comparison of therapeutic efficacy of developed prodrugs with CPT-11 in A549 lung orthotopic tumor-bearing mice.** Female 7-week-old BALB/cAJcl-*nu*/*nu* mice with free access to food and water before experiments were used. For tumor implantation, luciferase-expressing A549 cells ($1.0 \times 10^6$ cells) were suspended in 50 μL of PBS (-) containing 10% Matrigel and injected into the left lung of mice under anesthesia with a combination of medetomidine, midazolam and butorphanol. Before treatment, tumor implantation was confirmed by luciferin-luciferase bioluminescence imaging using an IVIS *in vivo* imager (PerkinElmer). Seventeen days after tumor implantation, SN38-βGal, SN38-βGlcNAc and CPT-11 were intravenously administered to mice via the tail vein at a dose of 40 mg/kg three times per week for 4 weeks. The dosing solutions were prepared in PBS (-) containing less than 9.0 μL of DMSO as a cosolvent, and were administered at a total volume of 100 μL. Lung tumor growth was monitored by bioluminescence imaging under isoflurane anesthesia after intraperitoneal administration of luciferin (3.0 mg) dissolved in 200 μL PBS (-). Weight changes were monitored over 4 weeks after initial administration of prodrugs. The statistical significance of differences in anticancer efficacy was analyzed by one-way ANOVA with Tukey's multiple comparisons test.

**Combination of SN38-βGlcNAc with anti-PD-1.** Female 7-week-old C57BL/6 mice with free access to food and water before experiments were used. LLC cells ($2.0 \times 10^6$ cells) in 50 μL of PBS (-) were injected into the shaved right flank and tumor growth was monitored until the median size reached about 100 $mm^3$. SN38-βGlcNAc was administered at a dose of 150 mg/kg via retro-orbital intravenous injection under isoflurane anesthesia three times per week for 2 weeks. The dosing solutions were prepared in PBS (-) containing less than 15 µL of DMSO as a cosolvent (total volume 100 µL). Anti-mouse CD279 (PD-1) (Clone RMP1-14)-Purified *in vivo* GOLD Functional Grade (100 µg) (Monoclonal, Product number: P362, Lot: 0923L620, LET Leinco Technologies, Inc.) was intraperitoneally administered two times per week for 2 weeks. The dosing solutions were prepared in PBS (-) and were administered at a total volume of 100 µL. Tumor volumes and weight changes were monitored for 3 weeks after initial administration. The statistical significance of differences in anticancer efficacy was analyzed by one-way ANOVA with Tukey's multiple comparisons test.

***Ex-vivo* fluorescence imaging of human lung surgical specimens containing both cancer and normal tissues**. A 50 µM solution of fluorescence probe (1-2 mL) in PBS (-) containing 0.5% (*v/v*) DMSO as a co-solvent was added to a 3.5 or 5.0 cm dish containing a human surgical specimen so that the tissue was completely soaked with probe solution. Fluorescence images of lung specimens were evaluated using the Maestro *in-vivo* imaging system as described above. The green filter setting (Ex/Em = 490 nm/550 nm long-pass) was used for HMRef-βGal and HMRef-βGlcNAc. The red filter setting (Ex/Em = 570 nm/610 nm long-pass) was used for 2'Me-SPiDER-Red-βGal. Exposure time was set at 100-20 msec depending on the level of fluorescence intensity from specimens. The stage and lamp were both set at position 1 or 2.

**Statistical analysis.** ROC curves were drawn with EZR software (Saitama Medical Centre, Jichi Medical University).[5] Statistical comparisons were made with KaleidaGraph Version 5.0 software (HULINKS).

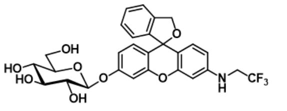

**1** (HMRef-β-D-Glc)

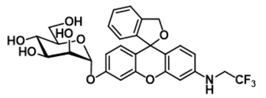

**5** (HMRef-α-D-Man)
HMRef-αMan

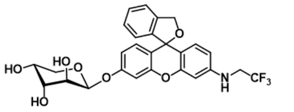

**9** (HMRef-α-D-Ara)

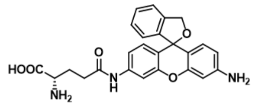

gGlu-HMRG

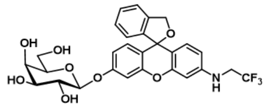

**2** (HMRef-β-D-Gal)
HMRef-βGal

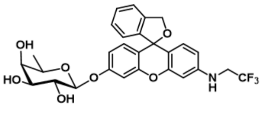

**6** (HMRef-β-D-Fuc)

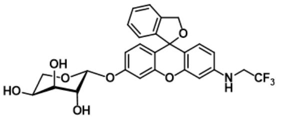

**10** (HMRef-α-L-Ara)

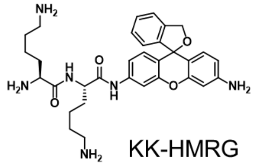

KK-HMRG

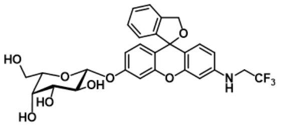

**3** (HMRef-β-L-Gal)

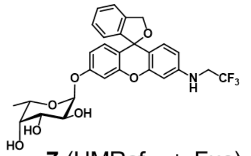

**7** (HMRef-α-L-Fuc)

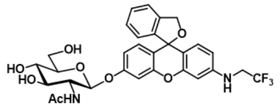

**11** (HMRef-β-D-GlcNAc)
HMRef-βGlcNAc

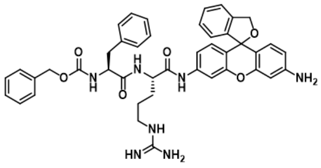

ZFR-HMRG
(Z-Phe-Arg-HMRG)

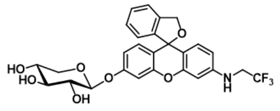

**4** (HMRef-β-D-Xyl)

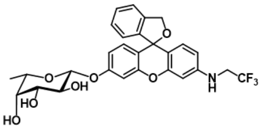

**8** (HMRef-β-L-Fuc)

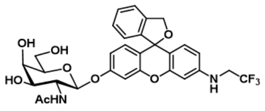

**12** (HMRef-β-D-GalNAc)

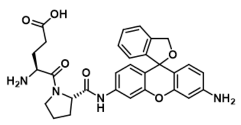

EP-HMRG

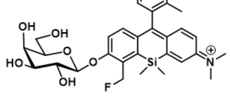

2’Me-SPiDER-Red-βGal
(2’Me-4CH$_2$F-Sirhodol-βGal)

**Figure S1.** Representative fluorescence probes used in this study.

**Table S1. Selected probes for activity-based screening used in this study.** Upper hit peptidase-reactive fluorescence probes and glycosidase-reactive fluorescence probes from our previous studies[6-14] were selected and evaluated in the activity-based screening for the prodrug development.

| Probes | Reported targets | References |
|---|---|---|
| KK-HMRG | PSA | Upper hit probes in NSCLC (Ref. 6-8) |
| GP-HMRG | DPP-IV | |
| PP-HMRG | - | |
| KR-HMRG | - | |
| KA-HMRG | - | |
| QA-HMRG | PSA, DPP-IV | |
| AY-HMRG | - | |
| GL-HMRG | - | |
| GY-HMRG | - | |
| SF-HMRG | - | |
| gGlu-HMRG | GGT | |
| EP-HMRG | DPP-IV | Upper hit probes in esophageal cancer (Ref. 9) |
| PR-HMRG | CAPN1 | Upper hit probes in glioma (Ref. 10) |
| GY-HMRG | - | |
| GH-HMRG | - | |
| KH-HMRG | APN | Upper hit probes in stomach cancer cancer (Ref. 6) |
| PM-HMRG | PSA | Upper hit probes in cholangiocarcinoma (Ref. 11) |
| SL-HMRG | - | |
| RK-HMRG | - | Upper hit probes in colorectal cancer lung metastasis (Ref. 12) |
| ZFR-HMRG | CATs | Cathepsins-reactive probes (Ref. 13) |
| ZRR-HMRG | CATs | |
| 1, HMRef-β-D-Glc | β-glucosidase | Glycosidase-reactive probes (Ref. 14) |
| 2, HMRef-β-D-Gal | β-galactosidase | |
| 3, HMRef-β-L-Gal | - | |
| 4, HMRef-β-D-Xyl | - | |
| 5, HMRef-α-D-Man | α-mannosidase | |
| 6, HMRef-β-D-Fuc | - | |
| 7, HMRef-α-L-Fuc | α-fucosidase | |
| 8, HMRef-β-L-Fuc | - | |
| 9, HMRef-α-L-Ara | - | |
| 10, HMRef-α-D-Ara | - | |
| 11, HMRef-β-D-GlcNAc | β-hexosaminidase | |
| 12, HMRef-β-D-GalNAc | - | |

**Table S2. Organ weights used to determine background scores.** Organ weights (g) were evaluated from 7-week-old healthy female ICR mice (n = 3).

| | Body weight | Brain | Heart | Thymus | Lung | Liver | Stomach | Pancreas | Spleen | Kidney | Intestine | Colon | Uterus |
|---|---|---|---|---|---|---|---|---|---|---|---|---|---|
| Mouse 1 | 31.6 | 0.491 | 0.146 | 0.083 | 0.203 | 1.784 | 0.259 | 0.229 | 0.146 | 0.410 | 1.565 | 0.420 | 0.095 |
| Mouse 2 | 31.6 | 0.531 | 0.147 | 0.085 | 0.182 | 1.760 | 0.248 | 0.322 | 0.137 | 0.446 | 1.435 | 0.345 | 0.145 |
| Mouse 3 | 30.9 | 0.521 | 0.128 | 0.076 | 0.173 | 1.749 | 0.289 | 0.439 | 0.160 | 0.382 | 1.790 | 0.273 | 0.128 |
| Average | 31.4 | 0.515 | 0.140 | 0.081 | 0.186 | 1.764 | 0.265 | 0.330 | 0.147 | 0.413 | 1.597 | 0.346 | 0.123 |
| s.d. | 0.3 | 0.017 | 0.009 | 0.004 | 0.013 | 0.015 | 0.017 | 0.086 | 0.009 | 0.026 | 0.146 | 0.060 | 0.021 |

7-week-old female Jcl:ICR mice (n=3)

**Table S3. Definition and nomenclature of background score notation.** Background scores were defined to quantify systemic and organ-specific probe activation. The organ background score ($B_{org}$) was calculated for each organ, and the total background score ($B_{tot}$) was defined as the sum of $B_{org}$ across all examined organs. *In vitro* and *in vivo* background scores were defined based on the probe conversion rate (CVR) or fluorescence intensity (a.u.), respectively, each multiplied by organ weight. Organ-specific notations follow standardized three-letter abbreviations derived from physiologically relevant terminology (e.g., Cer (cerebral), Hep (hepatic), Ren (renal)).

| Category | Notation | Definition |
|---|---|---|
| **General notations** | | |
| Organ background score | $B_{org}$ | Background score for each organ |
| Total background score | $B_{tot}$ | Sum of $B_{org}$ across all examined organs ($B_{tot} = \Sigma B_{org}$) |
| *In vitro* organ background score | $B_{org,vitro}$ | Background score in each organ measured *in vitro*, defined as the product of CVR of fluorescence probe in each organ and organ weight |
| *In vivo* organ background score | $B_{org,vivo}$ | Background score in each organ measured *in vivo*, defined as the product of fluorescence intensity (a.u.) in each organ and organ weight |
| *In vitro* total background score | $B_{tot,vitro}$ | Sum of $B_{org,vitro}$ across all examined organs ($B_{tot,vitro} = \Sigma B_{org,vitro}$) |
| *In vivo* total background score | $B_{tot,vivo}$ | Sum of $B_{org,vivo}$ across all examined organs ($B_{tot,vivo} = \Sigma B_{org,vivo}$) |
| **Organ-specific notations** | | |
| Brain | $B_{Cer}$ | Background score in brain |
| Heart | $B_{Car}$ | Background score in heart |
| Thymus | $B_{Thy}$ | Background score in thymus |
| Lung | $B_{Pul}$ | Background score in lung |
| Liver | $B_{Hep}$ | Background score in liver |
| Stomach | $B_{Gas}$ | Background score in stomach |
| Pancreas | $B_{Pan}$ | Background score in pancreas |
| Spleen | $B_{Spl}$ | Background score in spleen |
| Kidney | $B_{Ren}$ | Background score in kidney |
| Intestine | $B_{Int}$ | Background score in intestine |
| Colon | $B_{Col}$ | Background score in colon |
| Uterus | $B_{Ute}$ | Background score in uterus |

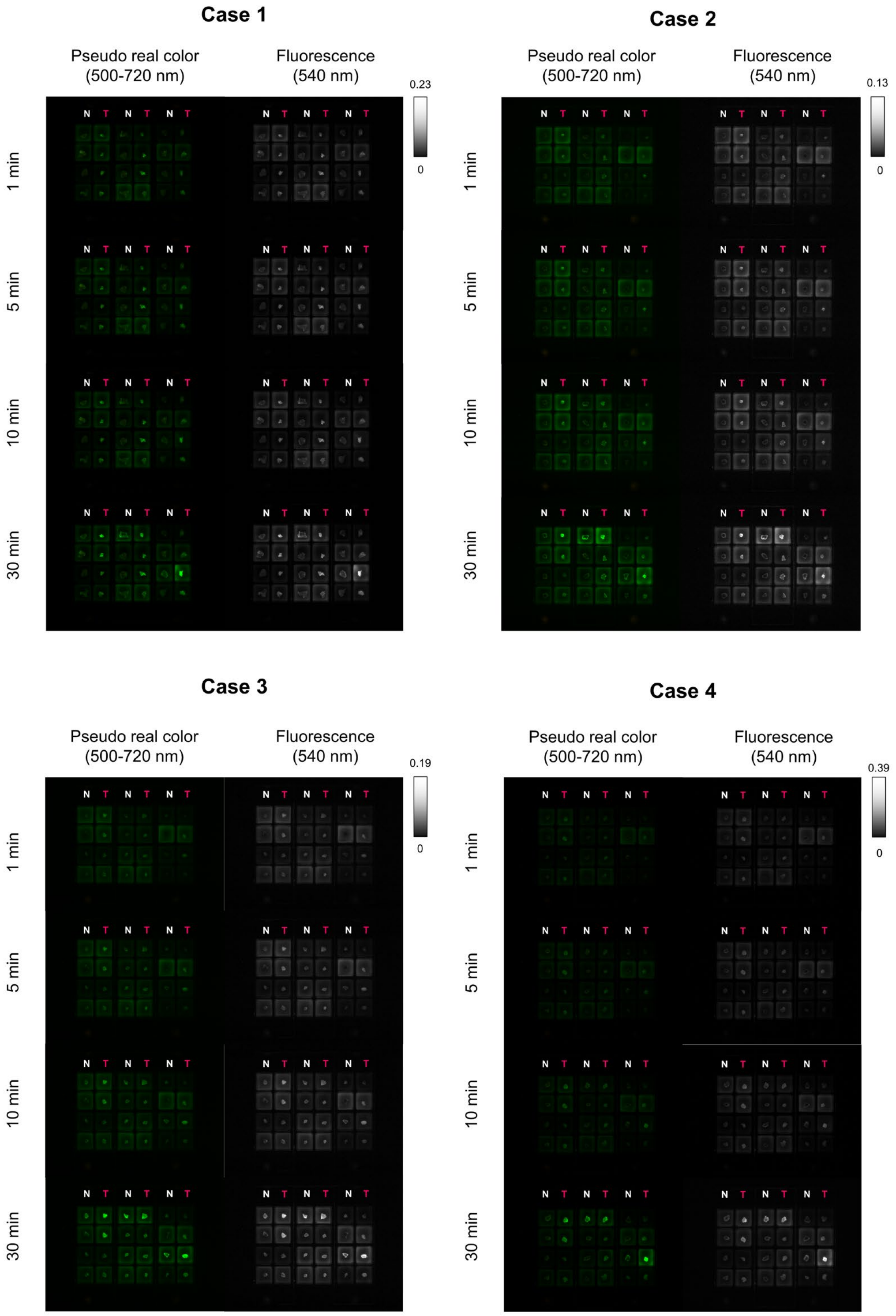


**Figure S2. Representative time-dependent fluorescence images in the screening using surgically resected normal lung and lung cancer tissues.** Cases 1-4 represent screening using specimens from 4 different lung cancer patients. Exposure time = 100 msec. [Fluorescent probe] = 50 μM. Scale bar, 2 cm.

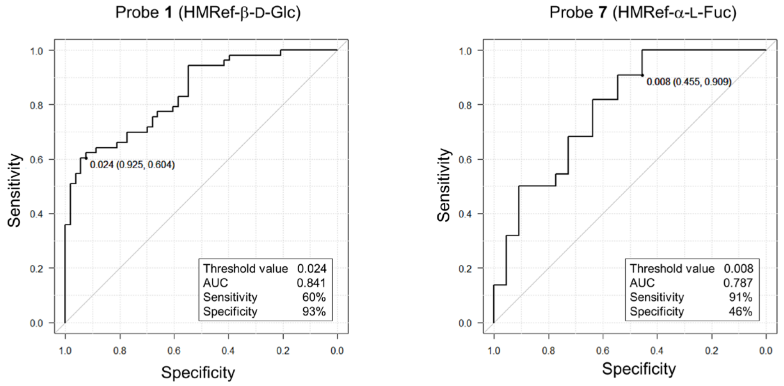


**Figure S3. ROC curves of HMRef-β-D-Glc and HMRef-α-L-Fuc for lung cancer detection.** Threshold value, sensitivity, specificity and AUC of each probe were evaluated from the ROC curve. Normal (N = 53), Ad (N = 35) and SCC (N = 18) tissues were examined with HMRef-β-D-Glc. Normal (N = 22), Ad (N = 10) and SCC (N = 12) tissues were examined with HMRef-α-L-Fuc.

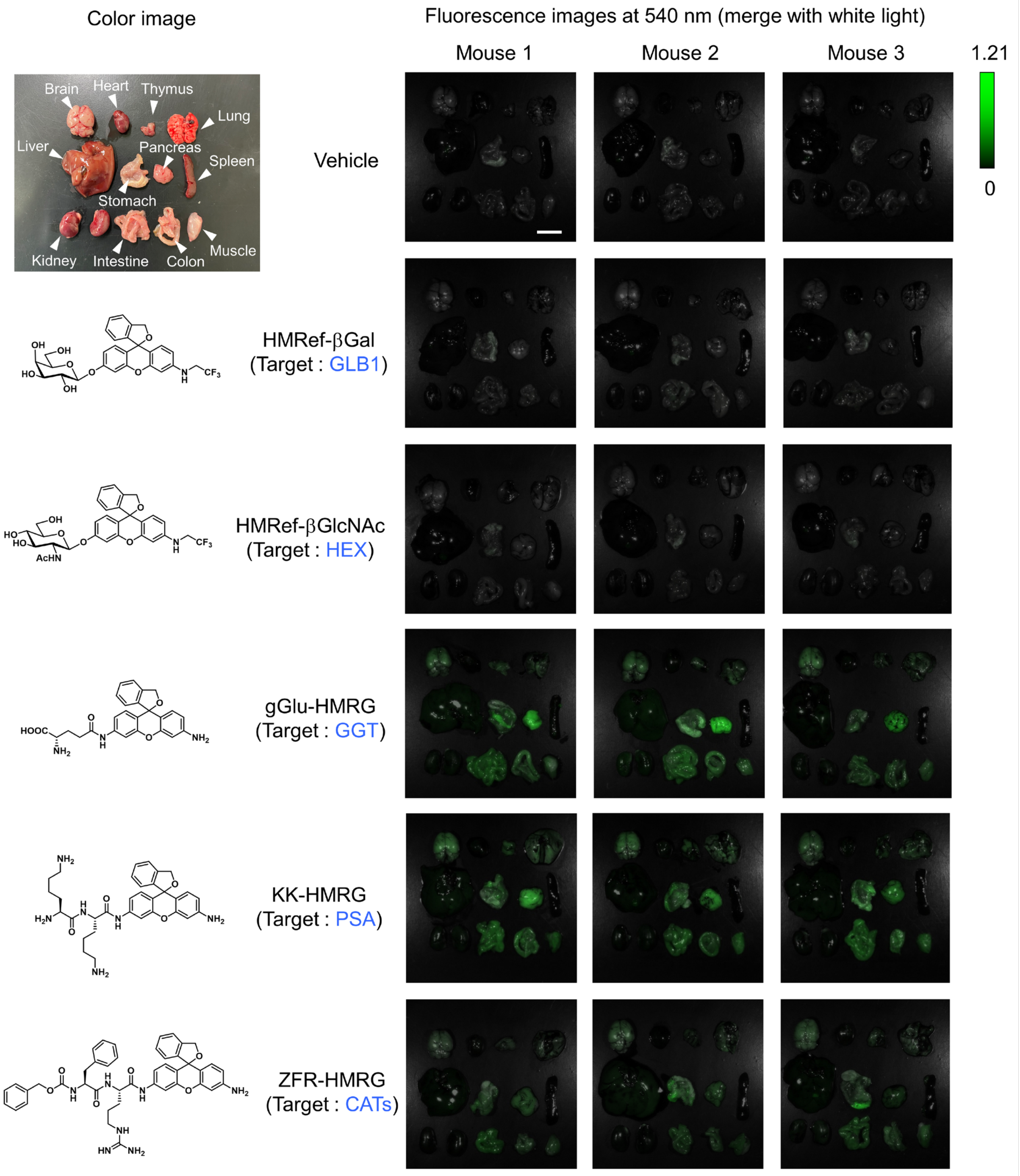


**Figure S4. Fluorescence images of resected mouse organs after systemic administration of each fluorescent probe.** Organs (brain, heart, lung, thymus, liver, stomach, pancreas, spleen, kidney, intestine, colon and muscle) were resected 20 min after systemic administration of 5 mg/kg fluorescent probe. ICR mice were used (n = 3). Scale bar, 1 cm.

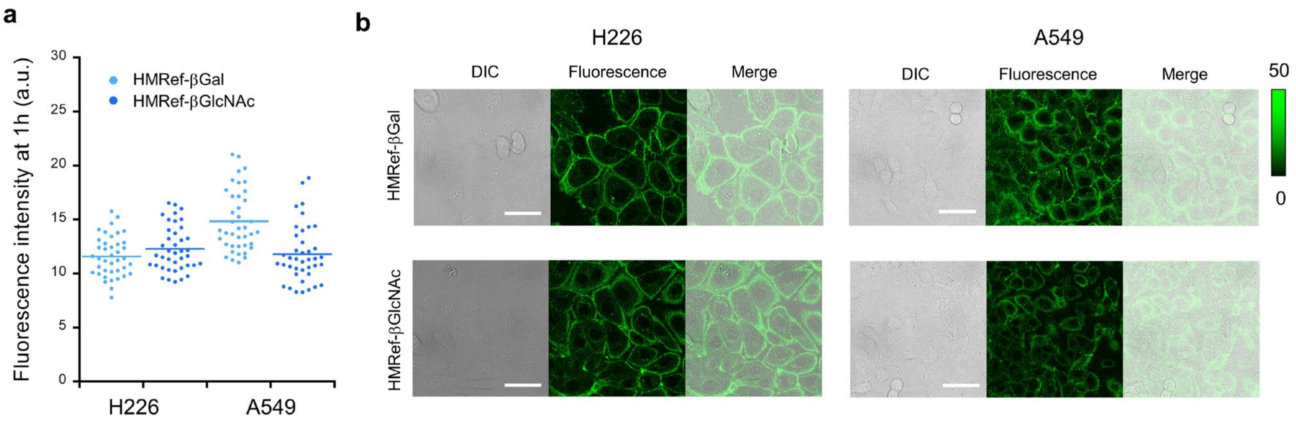


**Figure S5. Fluorescence imaging of glycosidase activities using HMRef-βGal and HMRef-βGlcNAc in lung cancer cell lines.** (**a**) Dot-plot diagram of single-cell fluorescence intensities at 1 h after addition of HMRef-βGal or HMRef-βGlcNAc (n = 40 for each group). The line in the diagram represents the average. (**b**) Confocal images of fluorescence intensity in NCI-H226 and A549 lung cancer cells treated with HMRef-βGal or HMRef-βGlcNAc. Cells were incubated with 20 μM fluorescence probe for 1 h at 37 °C, and fluorescence images were obtained. Ex/Em = 498 nm/505-600 nm. [Fluorescent probes] = 20 μM. Scale bars, 50 μm.

a

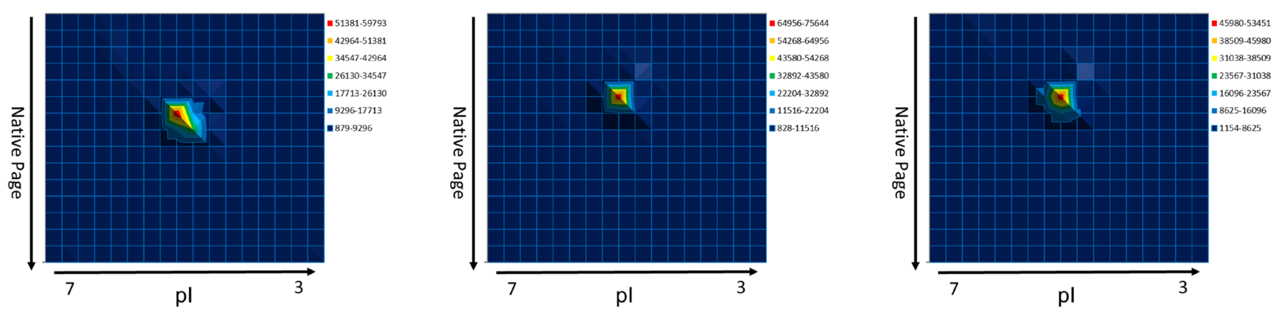


b

**P06865 (100%), 60,704.0 Da**
**Beta-hexosaminidase subunit alpha OS=Homo sapiens OX=9606 GN=HEXA PE=1 SV=2**
**3 exclusive unique peptides, 3 exclusive unique spectra, 4 total spectra, 41/529 amino acids (8% coverage)**

| | | | | |
|---|---|---|---|---|
| MTSSRLWFSL | LLAAAFAGRA | TALWPWPQNF | QTSDQRYVLY | PNNFQFQYDV |
| SSAAQPGCSV | LDEAFQRYRD | LLFGSGSWPR | PYLTGKRHTL | EKNVLVVSVV |
| TPGCNQLPTL | ESVENYTLTI | NDDQCLLLSE | TVWGALR**GLE** | **TFSQLVWK**SA |
| EGTFFINKTE | IEDFPRFPHR | **GLLLDTSR**HY | LPLSSILDTL | DVMAYNKLNV |
| FHWHLVDDPS | FPYESFTFPE | LMRKGSYNPV | THIYTAQDVK | EVIEYARLRG |
| IRVLAEFDTP | GHTLSWGPGI | PGLLTPCYSG | SEPSGTFGPV | NPSLNNTYEF |
| MSTFFLEVSS | VFPDFYLHLG | GDEVDFTCWK | SNPEIQDFMR | KKGFGEDFKQ |
| LESFYIQTLL | DIVSSYGKGY | VVWQEVFDNK | VKIQPDTIIQ | VWREDIPVNY |
| MKELELVTKA | GFR**ALLSAPW** | **YLNR**ISYGPD | WKDFYIVEPL | AFEGTPEQKA |
| LVIGGEACMW | GEYVDNTNLV | PRLWPRAGAV | AERLWSNK**LT** | **SDLTFAYER**L |
| SHFRCELLRR | GVQAQPLNVG | FCEQEFEQT | | |

**P07686 (100%), 63,113.3 Da**
**Beta-hexosaminidase subunit beta OS=Homo sapiens OX=9606 GN=HEXB PE=1 SV=3**
**2 exclusive unique peptides, 2 exclusive unique spectra, 3 total spectra, 39/556 amino acids (7% coverage)**

| | | | | |
|---|---|---|---|---|
| MELCGLGLPR | PPMLLALLLA | TLLAAMLALL | TQVALVVQVA | EAARAPSVSA |
| KPGPALWPLP | LSVKMTPNLL | HLAPENFYIS | HSPNSTAGPS | CTLLEEAFRR |
| YHGYIFGFYK | WHHEPAEFQA | KTQVQQLLVS | ITLQSECDAF | PNISSDESYT |
| LLVKEPVAVL | KANRVWGALR | GLETFSQLVY | QDSYGTFTIN | ESTIIDSPRF |
| SHR**GILIDTS** | **R**HYLPVKIIL | KTLDAMAFNK | FNVLHWHIVD | DQSFPYQSIT |
| FPELSNKGSY | SLSHVYTPND | VRMVIEYARL | RGIRVLPEFD | TPGHTLSWGK |
| GQKDLLTPCY | SRQNKLDSFG | PINPTLNTTY | SFLTTFFKEI | SEVFPDQFIH |
| LGGDEVEFKC | WESNPKIQDF | MRQKGFGTDF | KKLESFYIQK | **VLDIIATINK** |
| GSIVWQEVFD | DKAK**LAPGTI** | **VEVWKDSAYP** | **EELSR**VTASG | FPVILSAPWY |
| LDLISYGQDW | RKYYKVEPLD | FGGTQKQKQL | FIGGEACLWG | EYVDATNLTP |
| RLWPRASAVG | ERLWSSKDVR | DMDDAYDRLT | RHRCRMVERG | IAAQPLYAGY |
| CNHENM | | | | |

**Figure S6. DEG assay using HMRef-βGlcNAc with surgically resected Ad specimen from human lung.** (**a**) 2D gel assay. The assays were carried out as reported. A single fluorescent spot was observed on 2D gels and HEXA and HEXB were identified by peptide mass fingerprinting analysis. Each lysate sample was evaluated on four gels. Gels were incubated with the probe at 37°C for 12 h. Proteins detected by peptide mass fingerprinting analysis are summarized in **Tables S3**. [HMRef-βGlcNAc] = 1 μM. (**b**) Detected peptide sequences in the peptide mass fingerprinting analysis. Unique peptides of HEXA and HEXB were detected from the fluorescent spot in DEG assay.

**Table S4. List of detected proteins in DEG assay of a human lung Ad surgical specimen.** 186 proteins were identified by peptide mass fingerprinting analysis. β-Hexosaminidase subunit α (HEXA) and β-hexosaminidase subunit β (HEXB) are highlighted in yellow.

| | Identified proteins | Molecular Weight | Protein identification probability |
|---|---|---|---|
| 1 | Cluster of Keratin, type II cytoskeletal 1 | 66 kDa | 100% |
| 2 | Alpha-actinin-4 | 105 kDa | 100% |
| 3 | Heat shock protein HSP 90-alpha | 85 kDa | 100% |
| 4 | Alpha-actinin-1 | 103 kDa | 100% |
| 5 | Endoplasmin | 92 kDa | 100% |
| 6 | Cluster of Vimentin | 54 kDa | 100% |
| 7 | UDP-glucose:glycoprotein glucosyltransferase 1 | 177 kDa | 100% |
| 8 | Aldehyde dehydrogenase, mitochondrial | 56 kDa | 100% |
| 9 | Heat shock cognate 71 kDa protein | 71 kDa | 100% |
| 10 | Serotransferrin | 77 kDa | 100% |
| 11 | Heat shock 70 kDa protein 1A | 70 kDa | 100% |
| 12 | Keratin, type I cytoskeletal 10 | 59 kDa | 100% |
| 13 | Keratin, type I cytoskeletal 18 | 48 kDa | 100% |
| 14 | Keratin, type I cytoskeletal 9 | 62 kDa | 100% |
| 15 | X-ray repair cross-complementing protein 5 | 83 kDa | 100% |
| 16 | Cluster of Actin, cytoplasmic 1 | 42 kDa | 100% |
| 17 | Spectrin alpha chain, non-erythrocytic 1 | 285 kDa | 100% |
| 18 | Keratin, type I cytoskeletal 19 | 44 kDa | 100% |
| 19 | Keratin, type II cytoskeletal 7 | 51 kDa | 100% |
| 20 | Neutral alpha-glucosidase AB | 107 kDa | 100% |
| 21 | Ubiquitin-like modifier-activating enzyme 1 | 118 kDa | 100% |
| 22 | Heat shock 70 kDa protein 4 | 94 kDa | 100% |
| 23 | Albumin OS=Homo sapiens | 69 kDa | 100% |
| 24 | Calpain-1 catalytic subunit | 82 kDa | 100% |
| 25 | Interleukin enhancer-binding factor 3 | 95 kDa | 100% |
| 26 | Inter-alpha-trypsin inhibitor heavy chain H2 | 106 kDa | 100% |
| 27 | Stress-70 protein, mitochondrial | 74 kDa | 100% |
| 28 | Endoplasmic reticulum chaperone BiP | 72 kDa | 100% |
| 29 | X-ray repair cross-complementing protein 6 | 70 kDa | 100% |
| 30 | Tryptophan--tRNA ligase, cytoplasmic | 53 kDa | 100% |
| 31 | Alpha-2-macroglobulin | 163 kDa | 100% |
| 32 | Thioredoxin reductase 1, cytoplasmic | 71 kDa | 100% |
| 33 | Heat shock protein 105 kDa | 97 kDa | 100% |
| 34 | Isocitrate dehydrogenase [NADP] cytoplasmic | 47 kDa | 100% |
| 35 | Heat shock protein HSP 90-beta | 83 kDa | 100% |
| 36 | Galectin-3-binding protein | 65 kDa | 100% |
| 37 | Ceruloplasmin | 122 kDa | 100% |
| 38 | Complement C3 | 187 kDa | 100% |
| 39 | Glyceraldehyde-3-phosphate dehydrogenase | 36 kDa | 100% |
| 40 | Nucleolin | 77 kDa | 100% |
| 41 | Dihydropyrimidinase-related protein 2 | 62 kDa | 100% |
| 42 | Cluster of Keratin, type I cytoskeletal 14 | 52 kDa | 100% |
| 43 | Prolyl 4-hydroxylase subunit alpha-1 | 61 kDa | 100% |
| 44 | Inter-alpha-trypsin inhibitor heavy chain H1 | 101 kDa | 100% |
| 45 | Peroxiredoxin-6 | 25 kDa | 100% |
| 46 | Aldehyde dehydrogenase 1A1 | 55 kDa | 100% |
| 47 | Protein disulfide-isomerase | 57 kDa | 100% |
| 48 | Cluster of Polyadenylate-binding protein 1 | 71 kDa | 100% |
| 49 | Adenosylhomocysteinase | 48 kDa | 100% |
| 50 | ATP-dependent RNA helicase A | 141 kDa | 100% |
| 51 | Translin | 26 kDa | 100% |
| 52 | Ubiquitin carboxyl-terminal hydrolase 7 | 128 kDa | 100% |
| 53 | 4-trimethylaminobutyraldehyde dehydrogenase | 54 kDa | 100% |
| 54 | Interleukin enhancer-binding factor 2 | 43 kDa | 100% |
| 55 | 60S acidic ribosomal protein P0 | 34 kDa | 100% |
| 56 | Eukaryotic initiation factor 4A-I | 46 kDa | 100% |
| 57 | Hemoglobin subunit beta | 16 kDa | 100% |
| 58 | 60 kDa heat shock protein, mitochondrial | 61 kDa | 100% |
| 59 | Elongation factor 1-gamma | 50 kDa | 100% |
| 60 | Transaldolase | 38 kDa | 100% |
| 61 | Hemoglobin subunit alpha | 15 kDa | 100% |
| 62 | Heterogeneous nuclear ribonucleoprotein U-like protein 1 | 96 kDa | 100% |

| | | | |
|---|---|---|---|
| 63 | L-lactate dehydrogenase B chain | 37 kDa | 100% |
| 64 | Lamin-B1 | 66 kDa | 100% |
| 65 | Citrate synthase, mitochondrial | 52 kDa | 100% |
| 66 | Carbonyl reductase [NADPH] 1 | 30 kDa | 100% |
| 67 | Aldo-keto reductase family 1 member C1 | 37 kDa | 100% |
| 68 | Insulin-degrading enzyme | 118 kDa | 100% |
| 69 | Elongation factor 1-alpha 1 | 50 kDa | 100% |
| 70 | Translin-associated protein X | 33 kDa | 100% |
| 71 | Ferritin heavy chain | 21 kDa | 100% |
| 72 | Tubulin beta chain | 50 kDa | 100% |
| 73 | Major vault protein | 99 kDa | 100% |
| 74 | Gelsolin OS=Homo sapiens | 86 kDa | 100% |
| 75 | Phosphoglycerate mutase 1 | 29 kDa | 100% |
| 76 | Ras-related C3 botulinum toxin substrate 1 | 21 kDa | 100% |
| 77 | Hornerin | 282 kDa | 100% |
| 78 | Ribosome-binding protein 1 | 152 kDa | 100% |
| 79 | L-lactate dehydrogenase A chain | 37 kDa | 100% |
| 80 | Ubiquitin carboxyl-terminal hydrolase isozyme L1 | 25 kDa | 100% |
| 81 | Triosephosphate isomerase | 27 kDa | 100% |
| 82 | Peptidyl-prolyl cis-trans isomerase FKBP4 | 52 kDa | 100% |
| 83 | Cell division control protein 42 homolog | 21 kDa | 100% |
| 84 | Immunoglobulin heavy constant alpha 1 | 38 kDa | 100% |
| 85 | Polyubiquitin-B | 26 kDa | 100% |
| 86 | 60S ribosomal protein L12 | 18 kDa | 100% |
| 87 | Methanethiol oxidase | 52 kDa | 100% |
| 88 | SUMO-activating enzyme subunit 2 | 71 kDa | 100% |
| 89 | Proteasome activator complex subunit 3 | 30 kDa | 100% |
| 90 | Replication protein A 70 kDa DNA-binding subunit | 68 kDa | 100% |
| 91 | Heterogeneous nuclear ribonucleoprotein H | 49 kDa | 100% |
| 92 | Glycerol-3-phosphate phosphatase | 34 kDa | 100% |
| 93 | Protein AMBP | 39 kDa | 100% |
| 94 | Beta-hexosaminidase subunit alpha | 61 kDa | 100% |
| 95 | Glucosidase 2 subunit beta | 59 kDa | 100% |
| 96 | Transgelin-2 | 22 kDa | 100% |
| 97 | Dihydropyrimidinase-related protein 3 | 62 kDa | 100% |
| 98 | Heterogeneous nuclear ribonucleoprotein R | 71 kDa | 100% |
| 99 | Annexin A2 | 39 kDa | 100% |
| 100 | Lysosome-associated membrane glycoprotein 2 | 45 kDa | 100% |
| 101 | Protein disulfide-isomerase A6 | 48 kDa | 100% |
| 102 | Cluster of Peroxiredoxin-1 | 22 kDa | 100% |
| 103 | Heterogeneous nuclear ribonucleoprotein D0 | 38 kDa | 100% |
| 104 | Beta-hexosaminidase subunit beta | 63 kDa | 100% |
| 105 | Cluster of Plastin-2 | 70 kDa | 100% |
| 106 | Tubulin alpha-1B chain | 50 kDa | 100% |
| 107 | Hemopexin | 52 kDa | 100% |
| 108 | TBC1 domain family member 4 | 147 kDa | 100% |
| 109 | 6-phosphogluconate dehydrogenase, decarboxylating | 53 kDa | 100% |
| 110 | Importin subunit beta-1 | 97 kDa | 100% |
| 111 | Glutathione S-transferase omega-1 | 28 kDa | 100% |
| 112 | Hsc70-interacting protein | 41 kDa | 100% |
| 113 | Chloride intracellular channel protein 1 | 27 kDa | 100% |
| 114 | 5'-3' exoribonuclease 2 | 109 kDa | 98% |
| 115 | Heterogeneous nuclear ribonucleoprotein U | 91 kDa | 100% |
| 116 | Importin-5 | 124 kDa | 100% |
| 117 | Prolyl 4-hydroxylase subunit alpha-2 | 61 kDa | 100% |
| 118 | Alpha-galactosidase A | 49 kDa | 100% |
| 119 | Bifunctional purine biosynthesis protein ATIC | 65 kDa | 100% |
| 120 | Serine/threonine-protein phosphatase 2B catalytic subunit alpha isoform | 59 kDa | 100% |
| 121 | Chloride intracellular channel protein 6 | 73 kDa | 100% |
| 122 | NIF3-like protein 1 | 42 kDa | 100% |
| 123 | Bleomycin hydrolase | 53 kDa | 100% |
| 124 | F-actin-capping protein subunit beta | 31 kDa | 100% |

| | | | |
|---|---|---|---|
| 125 | Heterogeneous nuclear ribonucleoprotein A/B | 36 kDa | 100% |
| 126 | General vesicular transport factor p115 | 108 kDa | 100% |
| 127 | Transcription elongation factor SPT6 | 199 kDa | 100% |
| 128 | Protein transport protein Sec31A | 133 kDa | 100% |
| 129 | Probable serine carboxypeptidase CPVL | 54 kDa | 100% |
| 130 | GTP-binding nuclear protein Ran | 24 kDa | 100% |
| 131 | Junction plakoglobin | 82 kDa | 100% |
| 132 | E3 UFM1-protein ligase 1 | 90 kDa | 100% |
| 133 | Ezrin | 69 kDa | 100% |
| 134 | U6 snRNA-associated Sm-like protein LSm4 | 15 kDa | 100% |
| 135 | Vinculin | 124 kDa | 100% |
| 136 | Protein disulfide-isomerase A3 | 57 kDa | 100% |
| 137 | Short-chain specific acyl-CoA dehydrogenase, mitochondrial | 44 kDa | 100% |
| 138 | Importin-9 | 116 kDa | 100% |
| 139 | Calpain small subunit 1 | 28 kDa | 100% |
| 140 | Acyl-protein thioesterase 1 | 25 kDa | 100% |
| 141 | Serine/threonine-protein phosphatase 2A 56 kDa regulatory subunit epsilon isoform | 55 kDa | 100% |
| 142 | Lysozyme C | 17 kDa | 95% |
| 143 | Heterogeneous nuclear ribonucleoprotein U-like protein 2 | 85 kDa | 41% |
| 144 | 14-3-3 protein theta | 28 kDa | 97% |
| 145 | 14-3-3 protein beta/alpha | 28 kDa | 97% |
| 146 | 14-3-3 protein gamma | 28 kDa | 97% |
| 147 | 14-3-3 protein zeta/delta | 28 kDa | 97% |
| 148 | Puromycin-sensitive aminopeptidase-like protein | 54 kDa | 95% |
| 149 | Immunoglobulin lambda-like polypeptide 5 | 23 kDa | 95% |
| 150 | Phosphoribosylformylglycinamidine synthase | 145 kDa | 95% |
| 151 | Selenoprotein F | 18 kDa | 95% |
| 152 | Cytosolic 10-formyltetrahydrofolate dehydrogenase | 99 kDa | 95% |
| 153 | Phosphoglycerate kinase 1 | 45 kDa | 95% |
| 154 | Carcinoembryonic antigen-related cell adhesion molecule 5 | 77 kDa | 95% |
| 155 | Alpha-enolase | 47 kDa | 95% |
| 156 | Tropomyosin alpha-3 chain | 33 kDa | 95% |
| 157 | 40S ribosomal protein SA | 33 kDa | 95% |
| 158 | Beta-galactosidase | 76 kDa | 95% |
| 159 | Cytosol aminopeptidase | 56 kDa | 95% |
| 160 | Protein S100-A11 | 12 kDa | 95% |
| 161 | Carcinoembryonic antigen-related cell adhesion molecule 6 | 37 kDa | 95% |
| 162 | Ras GTPase-activating-like protein IQGAP1 | 189 kDa | 95% |
| 163 | F-actin-capping protein subunit alpha-2 | 33 kDa | 95% |
| 164 | Rho GDP-dissociation inhibitor 1 | 23 kDa | 95% |
| 165 | Dermcidin OS=Homo sapiens | 11 kDa | 95% |
| 166 | Nicotinate-nucleotide pyrophosphorylase [carboxylating] | 31 kDa | 95% |
| 167 | 2,4-dienoyl-CoA reductase [(3E)-enoyl-CoA-producing], mitochondrial | 36 kDa | 95% |
| 168 | Keratinocyte proline-rich protein | 64 kDa | 95% |
| 169 | Scavenger receptor cysteine-rich type 1 protein M130 | 125 kDa | 95% |
| 170 | Protein TFG OS=Homo sapiens | 43 kDa | 95% |
| 171 | Coronin-1B OS=Homo sapiens | 54 kDa | 95% |
| 172 | N-alpha-acetyltransferase 15, NatA auxiliary subunit | 101 kDa | 95% |
| 173 | Prefoldin subunit 4 OS=Homo sapiens | 15 kDa | 95% |
| 174 | Nuclear migration protein nudC | 38 kDa | 95% |
| 175 | Hypoxia up-regulated protein 1 | 111 kDa | 95% |
| 176 | Histone H2A type 1-B/E OS=Homo sapiens | 14 kDa | 94% |
| 177 | Glutathione S-transferase P | 23 kDa | 94% |
| 178 | Immunoglobulin heavy constant gamma 1 | 36 kDa | 93% |
| 179 | Histone H2B type 2-E1 | 13 kDa | 91% |
| 180 | Glycogenin-1 | 39 kDa | 89% |
| 181 | Transmembrane glycoprotein NMB | 64 kDa | 88% |
| 182 | Prefoldin subunit 2 | 17 kDa | 87% |
| 183 | Clusterin | 52 kDa | 86% |
| 184 | Protein S100-A6 | 10 kDa | 84% |
| 185 | Voltage-dependent calcium channel gamma-6 subunit | 28 kDa | 32% |
| 186 | Myosin light chain kinase, smooth muscle | 211 kDa | 5% |

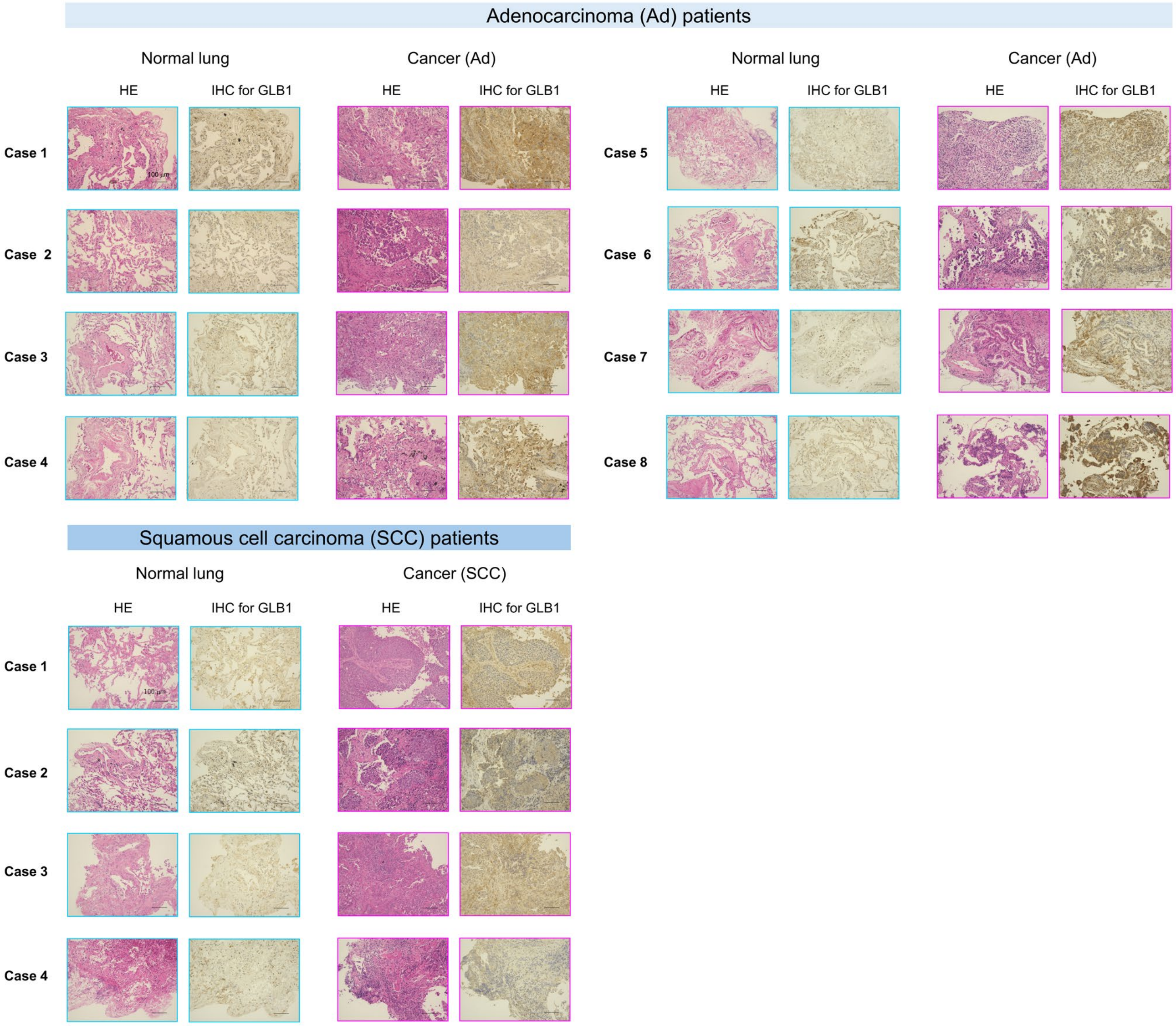


**Figure S7. IHC analysis for GLB1 in lung normal, Ad and SCC tissues.** Eight patients were evaluated for Ad (Cases 1-8). Four patients were evaluated for SCC (Cases 1-4). Scale bar = 100 µm.

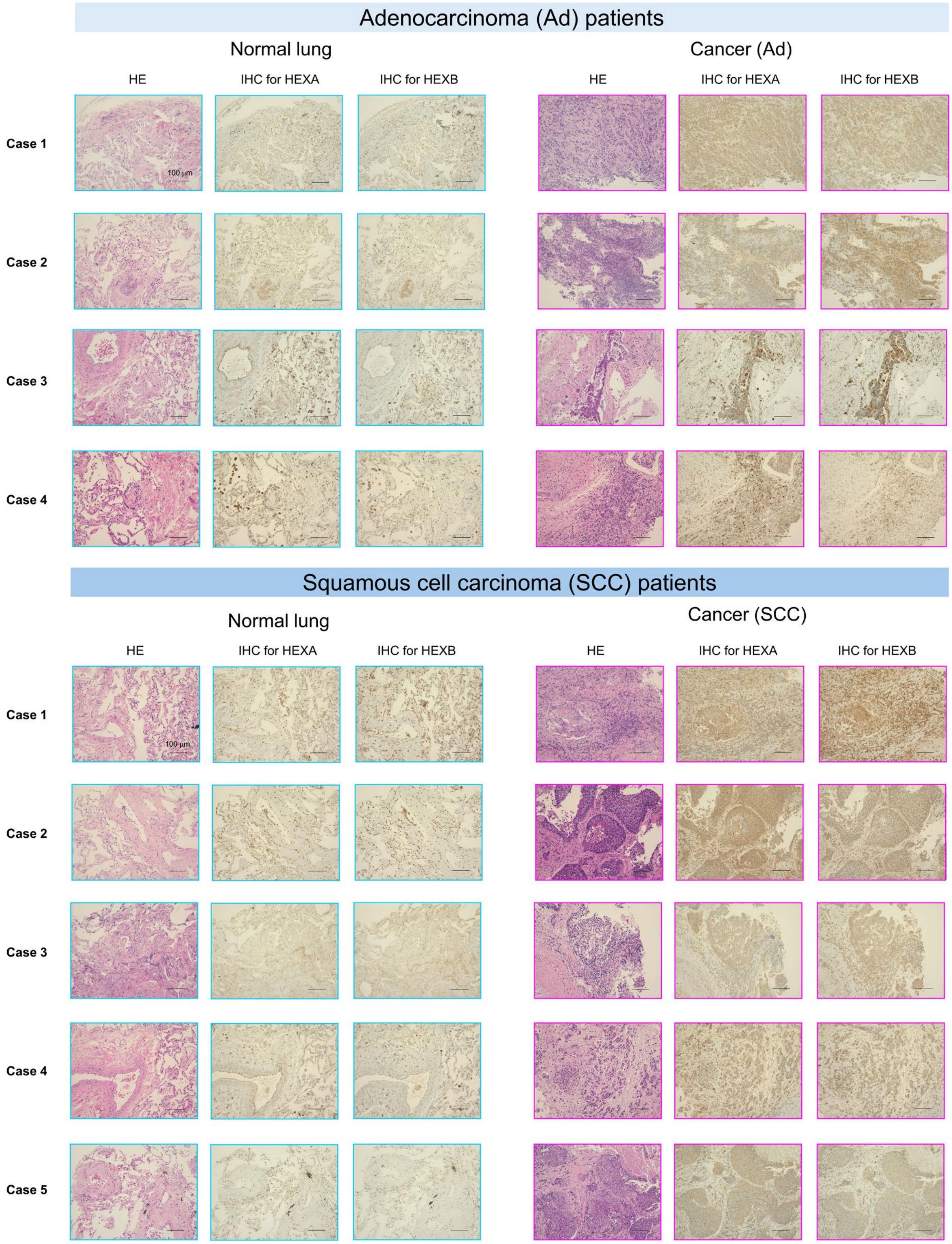


**Figure S8. IHC analysis for HEXA and HEXB in lung normal, Ad and SCC tissues.** Four patients were evaluated for Ad (Cases 1-4). Five patients were evaluated for SCC (Cases 1-5). Scale bar = 100 μm.

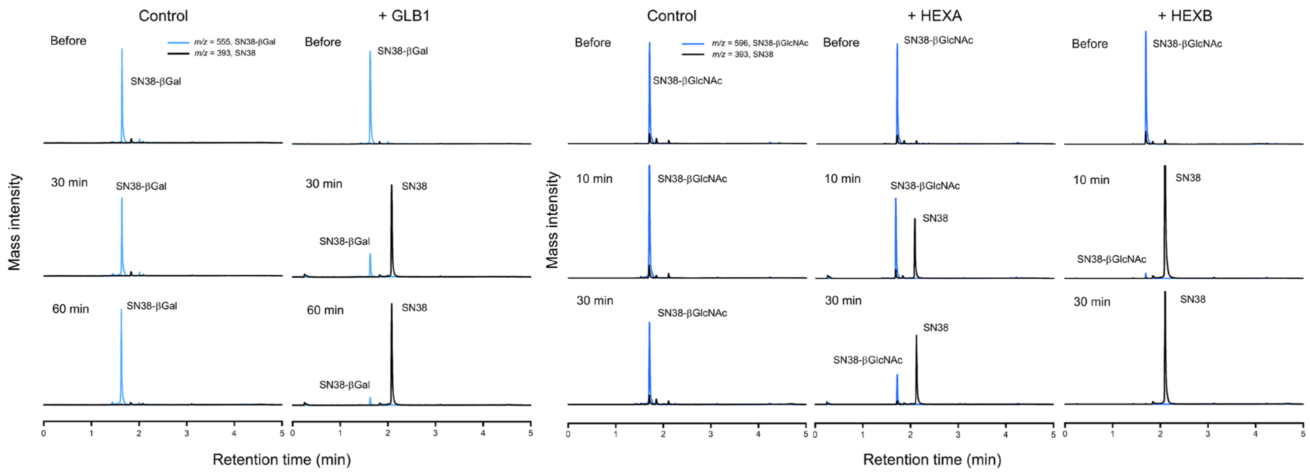


**Figure S9. Evaluation of reactivity of SN38-βGal and SN38-βGlcNAc with target enzymes by LC-MS.** LC-MS chromatograms of SN38-βGal in the presence and absence of GLB1. Production of SN38 was observed in the presence of GLB1 (left). LC-MS chromatograms of SN38-βGlcNAc in the presence and absence of HEXA or HEXB (right). Production of SN38 was observed in the presence of HEXA or HEXB.

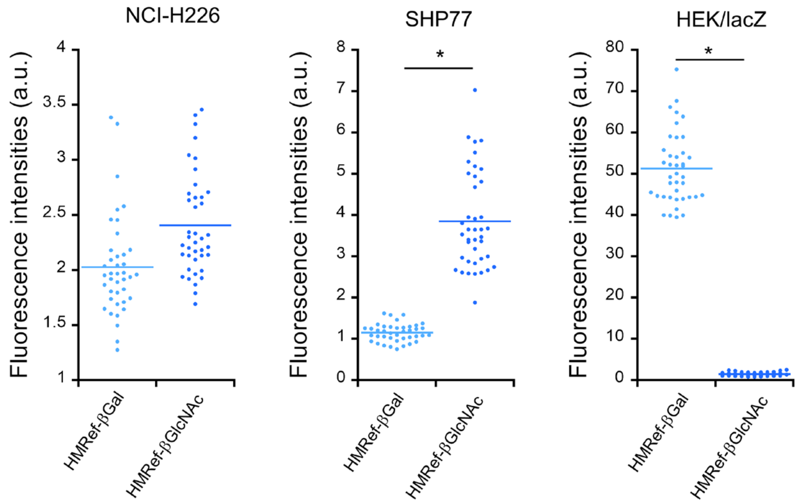


**Figure S10. Evaluation of enzyme activities in cultured cancer cells by live cell imaging.** Fluorescence images were captured 1 h after administration of each probe. Single-cell fluorescence intensities were quantified by drawing ROIs (n = 40). *$P$<0.01 by Welch's $t$-test. [Fluorescence probes] = 20 μM.

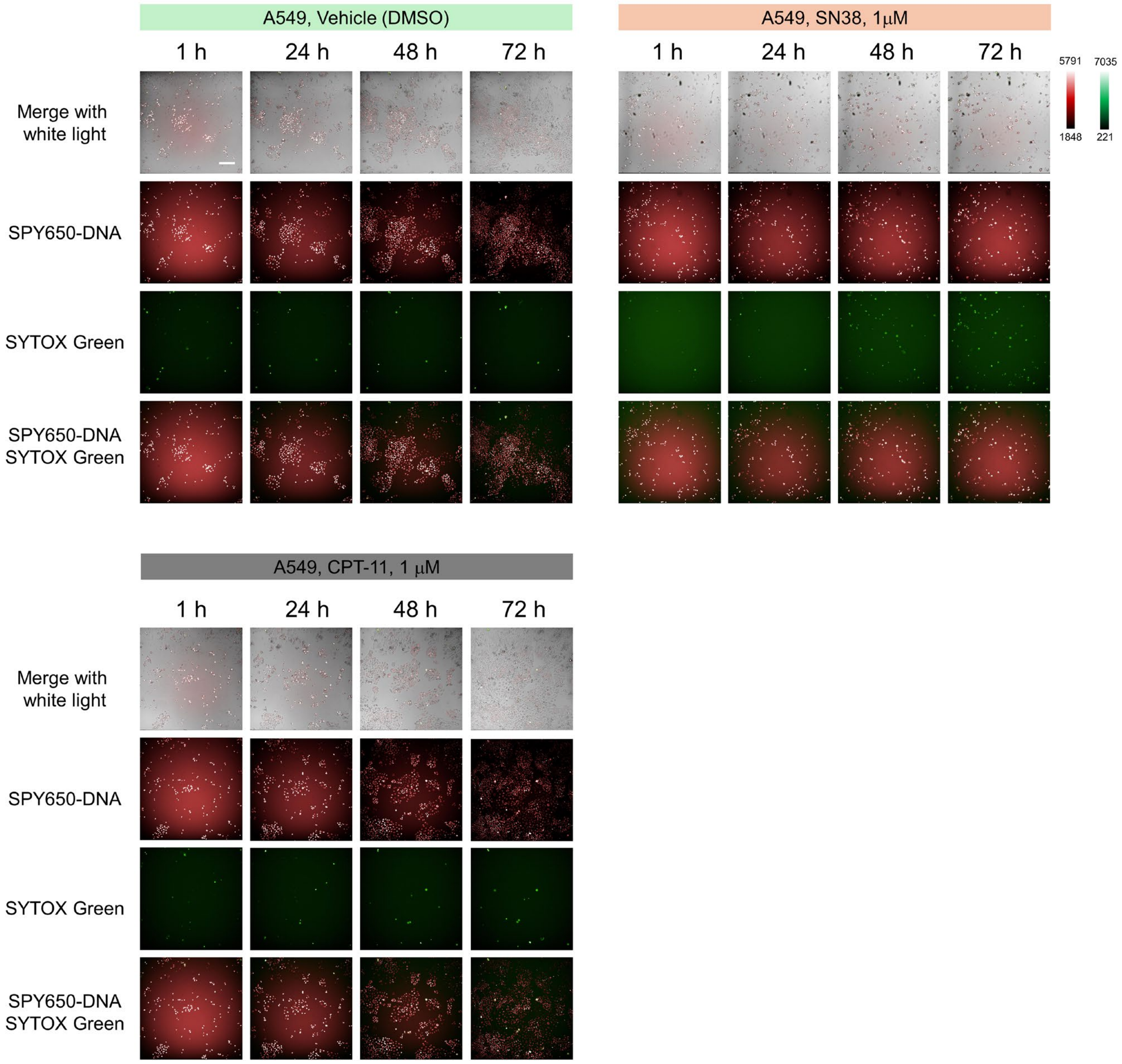


**Figure S11. Time-dependent fluorescence images of A549 cells in the presence of SN38 or CPT-11.** Nuclei were stained by SPY650-DNA. Dead cells were stained by SYTOX Green. SN38 significantly suppressed cell growth and increased SYTOX Green-positive dead cells over 72 h. CPT-11 did not show significant suppression of cell growth at the concentration of 1 μM. [drugs] = 1 μM. Scale bar, 200 μm.

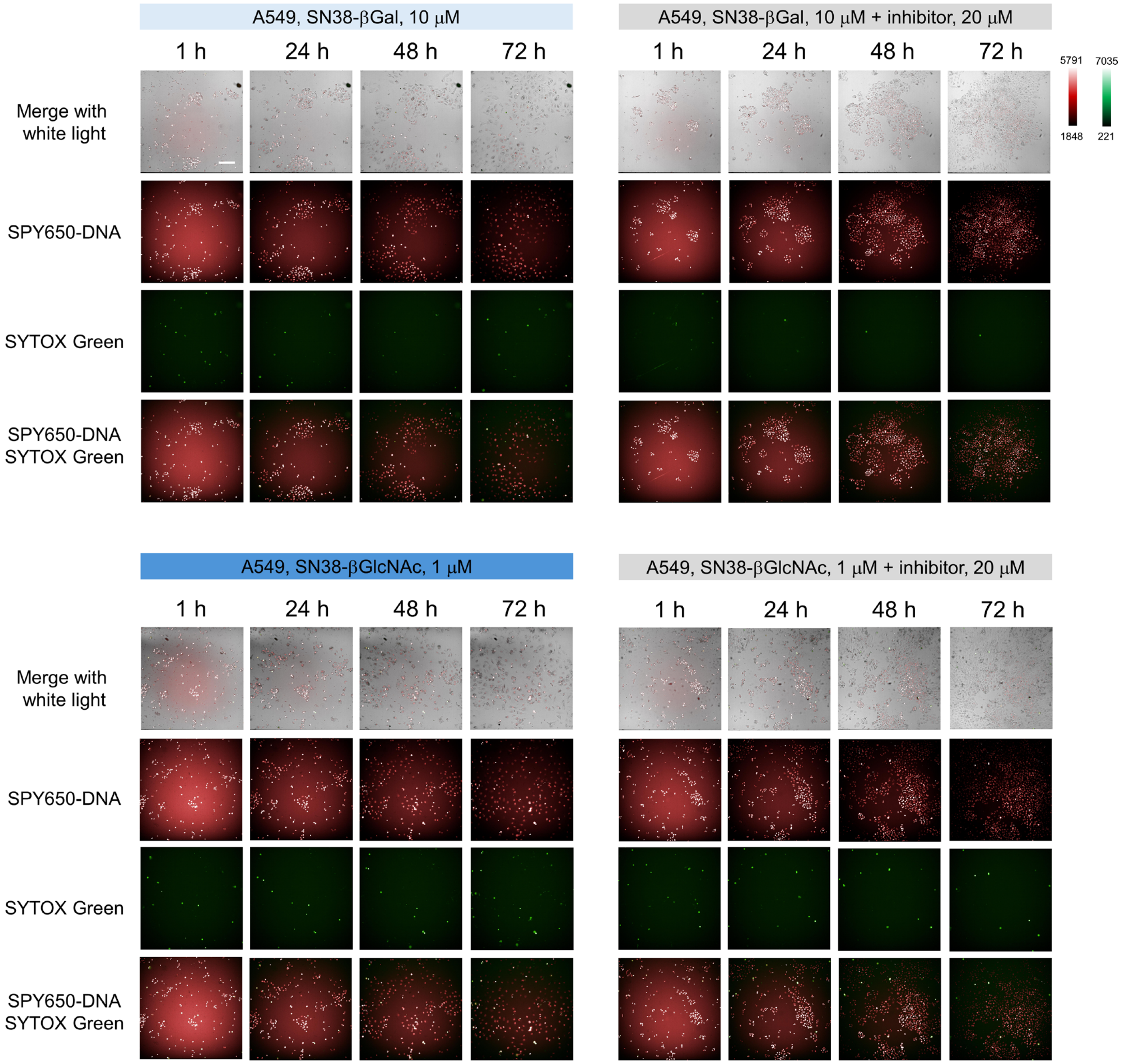


**Figure S12. Time-dependent fluorescence images of A549 cells in the presence of SN38-βGal or SN38-βGlcNAc.** Nuclei were stained by SPY650-DNA. Dead cells were stained by SYTOX Green. SN38-βGal and SN38-βGlcNAc inhibited A549 cell growth and their efficacy was suppressed in the presence of the corresponding glycosidase inhibitor. [SN38-βGal] = 10 μM, [SN38-βGlcNAc] = 1 μM, [inhibitor] = 20 μM. N-(n-Nonyl)deoxygalactonoijirimycin was used as an inhibitor for SN38-βGal. PUGNAc was used as an inhibitor for SN38-βGlcNAc. Scale bar, 200 μm.

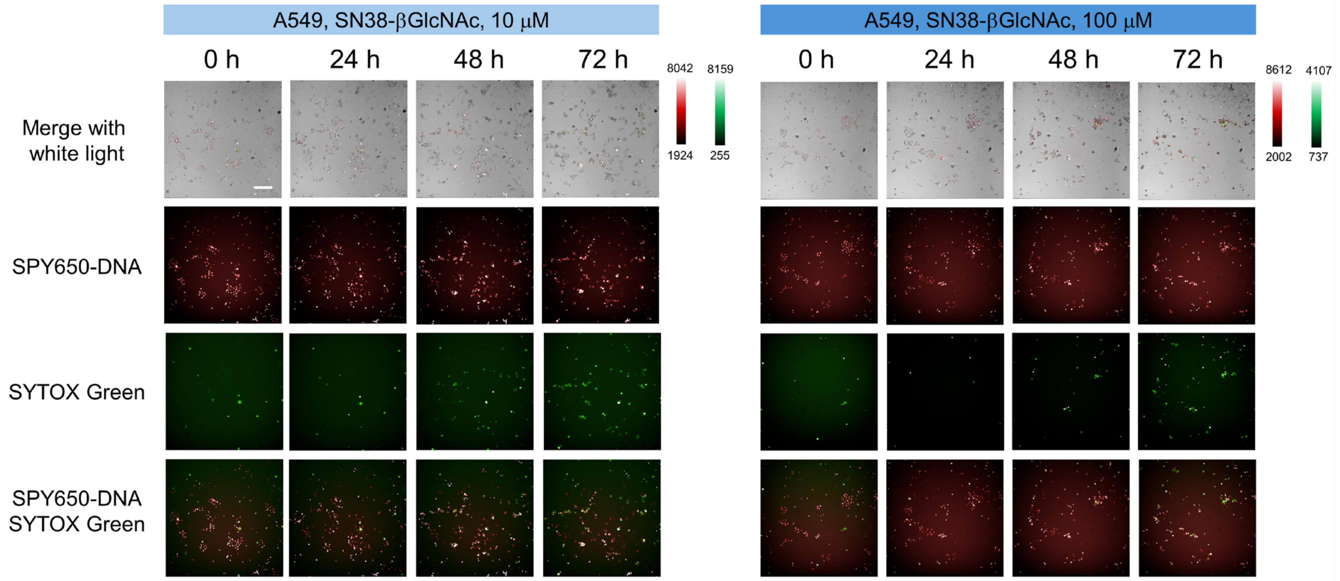


**Figure S13. Time-dependent fluorescence images of A549 cells in the presence of SN38-βGlcNAc.** Nuclei were stained by SPY650-DNA. Dead cells were stained by SYTOX Green. SN38-βGlcNAc significantly suppressed cell growth and increased SYTOX Green-positive dead cells over 72 h. [SN38-βGlcNAc] = 10 μM or 100 μM. Scale bar, 200 μm.

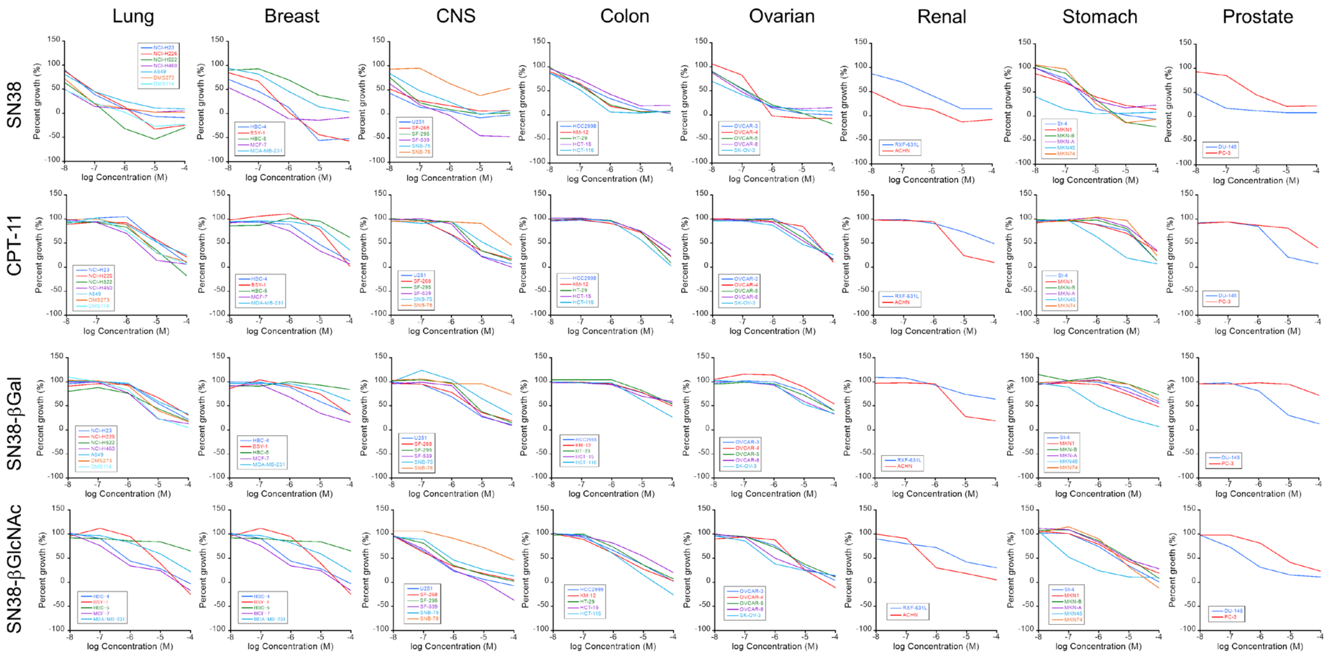


**Figure S14. Comprehensive evaluation of concentration-dependent cell growth inhibition in JFCR39 cancer cell panel.** Cell growth inhibition induced by SN38, CPT-11, SN38-βGal and SN38-βGlcNAc was examined in the JFCR39 cancer cell panel, excluding LOX-IMVI.

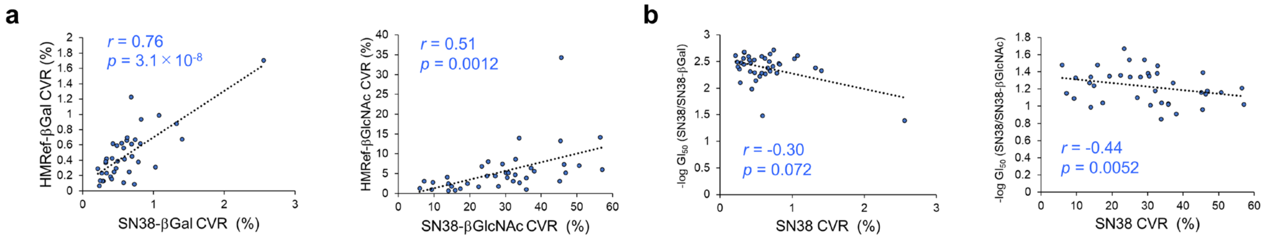


**Figure S15. Correlations between enzymatic activities or anticancer efficacy and produced SN38 for the developed prodrugs in the JFCR39 cancer cell panel.** (**a**) Correlations between CVR of HMRef-βGal to HMRef and CVR of SN38-βGal to SN38 (left), and correlations between CVR of HMRef-βGlcNAc to HMRef and CVR of SN38-βGlcNAc to SN38 (right). Cells with higher enzymatic activities tended to exhibit higher CVR of the prodrug to SN38. CVR values of fluorescence probes were measured after 3 h incubation with each cell line. CVR values of prodrugs were measured after 24 h incubation with each cell line. (**b**) Correlations between -log $GI_{50}$ (SN38/SN38-βGal) and CVR of SN38-βGal to SN38 (left), and correlations between -log $GI_{50}$ (SN38/SN38-βGlcNAc) and CVR of SN38-βGlcNAc to SN38 (right). Cells with higher CVR of the prodrug to SN38 tended to exhibit $GI_{50}$ values closer to those of SN38. These analyses were performed in JFCR39 panel cell lines, excluding LOX-IMVI.

### Table S5. Biochemical analysis in the MTD study.

| | | | Vehicle (DMSO) | | SN38-βGal 40 mg/kg × 3 | | SN38-βGal 120 mg/kg × 3 | | SN38-βGlcNAc 40 mg/kg × 3 | | SN38-βGlcNAc 200 mg/kg × 3 | |
|---|---|---|---|---|---|---|---|---|---|---|---|---|
| | | unit | Average | s.d. | Average | s.d. | Average | s.d. | Average | s.d. | Average | s.d. |
| Total protein | TP | g/dL | 4.13 | 0.15 | 4.50 | 0.38 | 4.25 | 0.21 | 4.50 | 0.08 | 4.48 | 0.15 |
| Albumin | ALB | g/dL | 2.98 | 0.10 | 3.20 | 0.16 | 3.03 | 0.10 | 3.13 | 0.05 | 3.15 | 0.10 |
| Albumin/Globulin ratio | A/G | | 2.60 | 0.21 | 2.53 | 0.47 | 2.48 | 0.19 | 2.27 | 0.08 | 2.38 | 0.01 |
| Total bilirubin | T-Bil | mg/dL | 0.1> | - | 0.1> | - | 0.1> | - | 0.1> | - | 0.1> | - |
| Aspartate aminotransferase | AST(GOT) | U/L | 446 | 155 | 334 | 271 | 520 | 88 | 69 | 20 | 222 | 186 |
| Alanine aminotransferase | ALT(GPT) | U/L | 66.0 | 24.3 | 70.8 | 24.5 | 86.5 | 41.3 | 18.5 | 1.3 | 31.8 | 13.6 |
| Lactate dehydrogenase | LDH | U/L | 1682 | 574 | 1396 | 985 | 1522 | 395 | 359 | 200 | 769 | 496 |
| Alkaline phosphatase | ALP | U/L | 107.3 | 12.1 | 97.8 | 8.6 | 120.5 | 27.3 | 95.8 | 8.7 | 79.5 | 16.3 |
| Amylase | AMY | U/L | 2463 | 280 | 2593 | 204 | 2571 | 453 | 2635 | 184 | 2597 | 322 |
| Lipase | LIP | U/L | 95.8 | 54.3 | 57.0 | 29.8 | 86.0 | 59.5 | 68.8 | 29.3 | 41.8 | 15.0 |
| Blood urea nitrogen | BUN | mg/dL | 20.48 | 1.97 | 20.1 | 2.6 | 22.2 | 5.2 | 18.9 | 3.3 | 17.0 | 1.2 |
| Creatinine | CRE | mg/dL | 0.13 | 0.02 | 0.1-0.15> | - | 0.1-0.12> | - | 0.1-0.15> | - | 0.1-0.15> | - |
| Total cholesterol | TC | mg/dL | 74.8 | 12.3 | 93.5 | 25.9 | 94.3 | 18.7 | 91.5 | 7.7 | 101.0 | 11.9 |
| Triglycerides | TG | mg/dL | 72.3 | 9.7 | 88.5 | 26.2 | 102.0 | 26.4 | 118.5 | 43.1 | 136.5 | 8.3 |
| Na | | mEq/L | 145.3 | 2.6 | 146.3 | 3.3 | 145.3 | 2.4 | 146.3 | 1.7 | 146.5 | 2.4 |
| K | | mEq/L | 6.75 | 0.83 | 5.7 | 1.9 | 7.1 | 1.6 | 4.3 | 0.4 | 4.7 | 0.6 |
| Cl | | mEq/L | 112.5 | 1.9 | 112.0 | 2.4 | 112.3 | 3.0 | 108.8 | 1.5 | 110.5 | 2.4 |
| Ca | | mg/dL | 9.10 | 0.34 | 9.35 | 0.49 | 9.58 | 0.39 | 9.5 | 0.5 | 9.40 | 0.47 |
| Inorganic phosphate | IP | mg/dL | 12.3 | 1.8 | 11.0 | 0.8 | 13.1 | 1.1 | 10.9 | 1.7 | 10.3 | 1.8 |
| Blood glucose | BG | mg/dL | 406 | 58 | 348 | 77 | 401 | 86 | 406 | 88 | 390 | 71 |
| Total bile acids | TBA | μmol/L | 8.40 | 5.62 | 4.70 | 4.60 | 6.65 | 7.62 | 1.5 | 0.5 | 1.73 | 0.50 |

n=4, s.d. represent standard devision.

### Table S6. Biochemical analysis in the repeated-dose toxicity study.

| | | | Vehicle (DMSO) | | SN38-βGal | | SN38-βGlcNAc | | CPT-11 | |
|---|---|---|---|---|---|---|---|---|---|---|
| | | unit | Average | s.d. | Average | s.d. | Average | s.d. | Average | s.d. |
| Total protein | TP | g/dL | 4.23 | 0.33 | 4.38 | 0.21 | 3.93 | 0.17 | 4.08 | 0.44 |
| Albumin | ALB | g/dL | 3.35 | 0.31 | 3.33 | 0.21 | 3.18 | 0.10 | 2.83 | 0.54 |
| Albumin/Globulin ratio | A/G | | 4.30 | 1.74 | 3.18 | 0.29 | 4.53 | 1.41 | 2.35 | 0.76 |
| Total bilirubin | T-Bil | mg/dL | 0.1> | - | 0.1> | - | 0.1> | - | 0.1> | - |
| Aspartate aminotransferase | AST(GOT) | U/L | 1253 | 779 | 821 | 142 | 1262 | 187 | 929 | 214 |
| Alanine aminotransferase | ALT(GPT) | U/L | 302 | 211 | 120 | 27 | 264 | 131 | 148 | 74 |
| Lactate dehydrogenase | LDH | U/L | >5000 | - | around 5000 | - | >5000 | - | around 5000 | - |
| Alkaline phosphatase | ALP | U/L | 86.3 | 24.3 | 85.3 | 17.1 | 80.5 | 12.5 | 56.5 | 41.9 |
| Amylase | AMY | U/L | 2054 | 313 | 2282 | 380 | 2263 | 69 | 2063 | 521 |
| Lipase | LIP | U/L | 60.5 | 24.8 | 67.0 | 20.8 | 72.0 | 18.3 | 61.3 | 21.8 |
| Blood urea nitrogen | BUN | mg/dL | 25.5 | 1.5 | 22.1 | 4.1 | 31.8 | 6.4 | 30.4 | 14.6 |
| Creatinine | CRE | mg/dL | 0.15 | 0.02 | 0.14 | 0.02 | 0.19 | 0.04 | 0.18 | 0.02 |
| Total cholesterol | TC | mg/dL | 81.0 | 8.4 | 71.8 | 11.0 | 67.0 | 12.4 | 74.0 | 20.1 |
| Triglycerides | TG | mg/dL | 63.25 | 8.18 | 57.75 | 11.93 | 62.25 | 11.03 | 50.25 | 8.18 |
| Na | | mEq/L | 143.8 | 2.2 | 146.3 | 0.5 | 140.5 | 1.7 | 142.8 | 8.7 |
| K | | mEq/L | 10.9 | 3.5 | 8.7 | 0.8 | 11.6 | 1.3 | 10.6 | 2.6 |
| Cl | | mEq/L | 101.3 | 1.5 | 108.3 | 2.6 | 101.0 | 0.8 | 106.3 | 2.1 |
| Ca | | mg/dL | 102.00 | 1.83 | 108.25 | 2.63 | 101.00 | 0.82 | 103.75 | 5.44 |
| Inorganic phosphate | IP | mg/dL | 11.05 | 2.27 | 12.98 | 1.12 | 13.55 | 0.66 | 12.48 | 0.81 |
| Blood glucose | BG | mg/dL | 300.5 | 14.6 | 279.0 | 102.2 | 338.8 | 60.8 | 263.3 | 49.0 |
| Total bile acids | TBA | μmol/L | 6.35 | 2.69 | 4.78 | 0.69 | 6.40 | 2.03 | 6.35 | 3.54 |

n=4, s.d. represent standard devision.

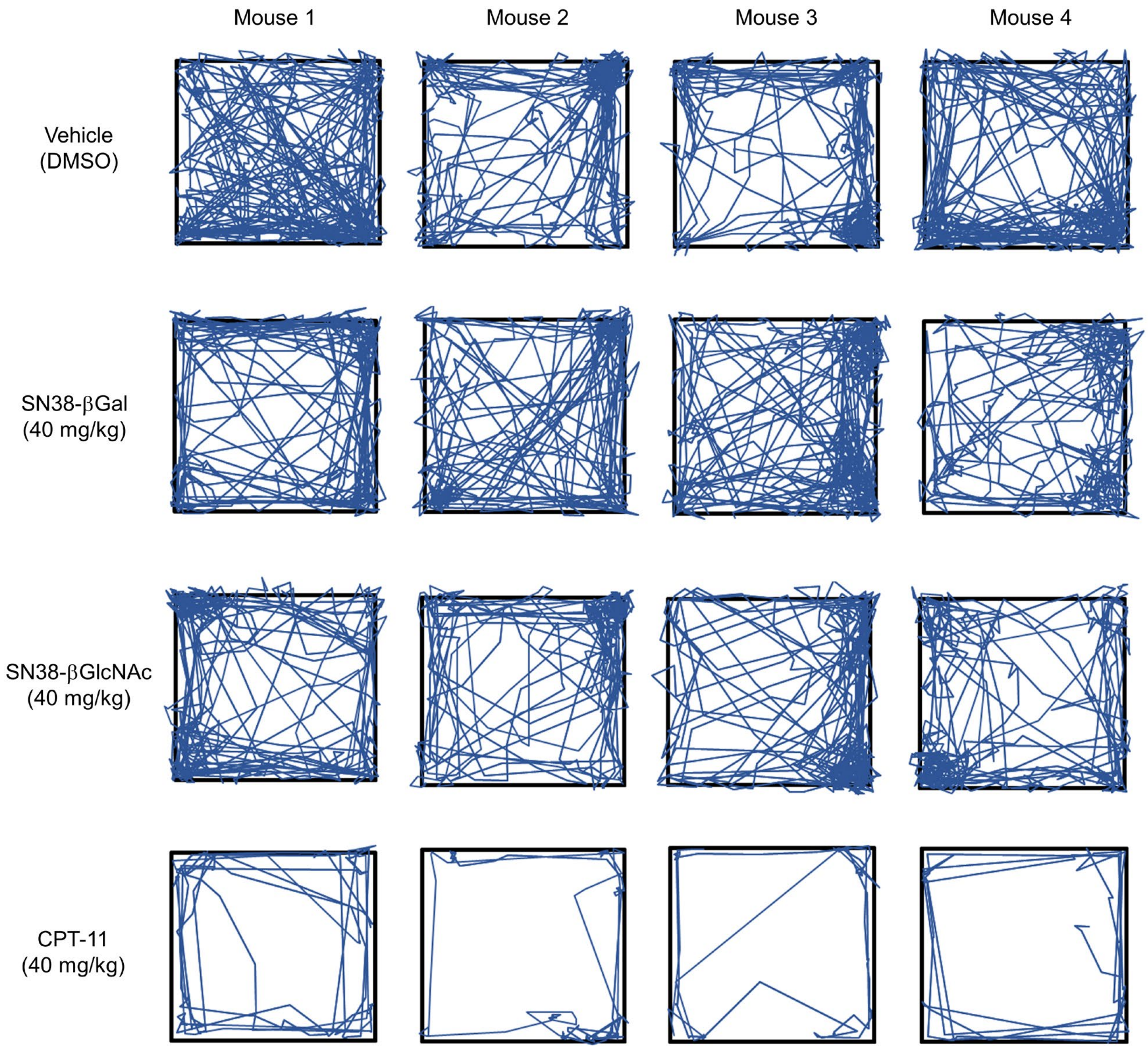


**Figure S16. Evaluation of the head movement trajectories in mice during 15 min following intravenous injection of each drug.** A significant decrease in locomotion was observed following the administration of CPT-11, but not SN38-βGal or SN38-βGlcNAc (n = 4 for each group).

**Table S7. Evaluation of $LD_{50}$ values of SN38-βGal and SN38-βGlcNAc.** The intravenous $LD_{50}$ values were determined to be over 300 mg/kg for SN38-βGal and over 500 mg/kg for SN38-βGlcNAc. The intravenous $LD_{50}$ values for CPT-11 were taken from the manufacturers' toxicology reports for irinotecan hydrochloride (Yakult Honsha Co., Ltd., 2022, version 14; Sawai Pharmaceutical Co., Ltd., 2022, version 9). Lethality after a single intravenous dose of 111 mg/kg in mice was also reported in the FDA review for irinotecan hydrochloride (NDA 20-571).

| Prodrugs (intravenous injection) | mouse | n | amount (mg/kg) | Survival rate (%) (just after administration) | Survival rate (%) (after 14 days) | $LD_{50}$ (mg/kg) |
|---|---|---|---|---|---|---|
| SN38-βGal | Jcl:ICR (female) | 10 | 200 | 100 | 100 | |
| | Jcl:ICR (female) | 10 | 300 | 100 | 100 | >300 |
| SN38-βGlcNAc | Jcl:ICR (female) | 10 | 200 | 100 | 100 | |
| | Jcl:ICR (female) | 10 | 500 | 100 | 100 | >500 |
| CPT-11 | female mouse | - | - | - | - | 132-133 |

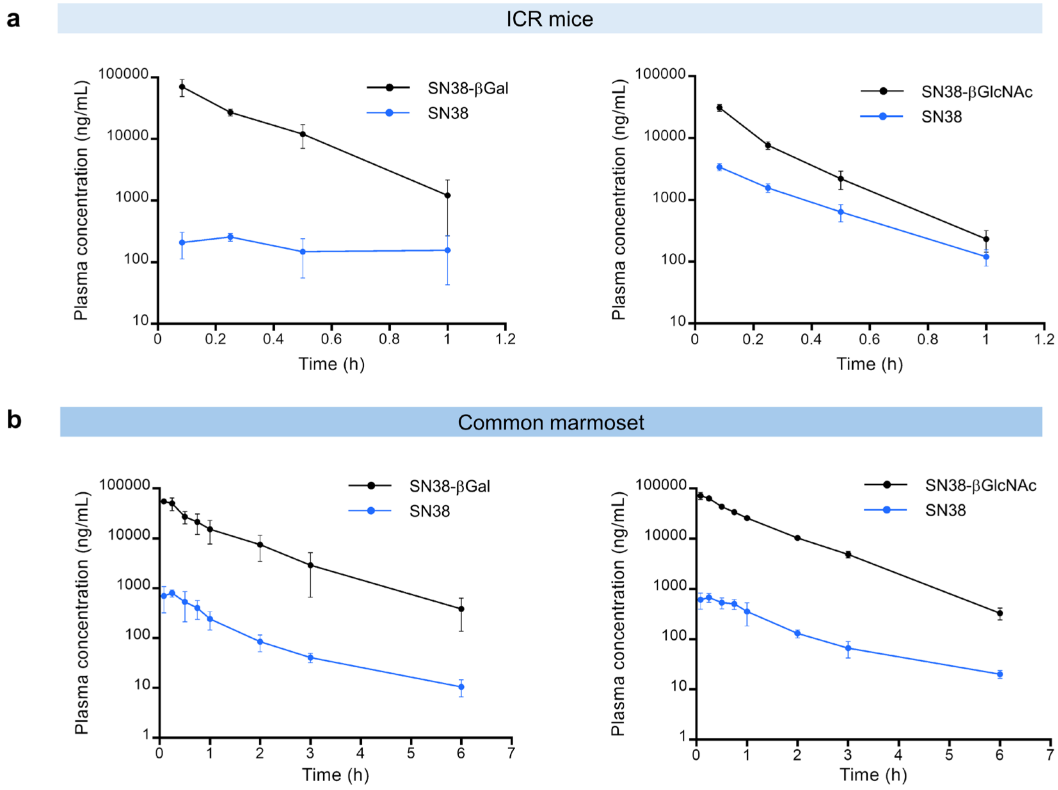


**Figure S17. Plasma concentration changes of developed prodrugs (logarithmic scale).** (**a**) Plasma concentrations of SN38-βGal (left), SN38-βGlcNAc (right) and released SN38 after intravenous injection of 40 mg/kg SN38-βGal or SN38-βGlcNAc to female ICR mice. Each data point represents the mean of three mice. Error bars represent s.d. (**b**) Plasma concentrations of SN38-βGal (left), SN38-βGlcNAc (right) and released SN38 after 11 min intravenous infusion of 10 mg/kg SN38-βGal or SN38-βGlcNAc to female common marmoset. Each data point represents the mean of three marmosets. Error bars represent s.d.

**Table S8. Pharmacokinetic parameters of SN38-βGal, SN38-βGlcNAc and released SN38 in common marmosets.** Pharmacokinetic parameters were calculated by non-compartmental analysis (NCA).

Pharmacokinetic parameters of SN38-βGal and released SN38.

| SN38-βGal | Age (Month) | Body weight (g) | $t_{1/2}$ (h) | $AUC_{0\text{-}6h}$ (μg・h/mL) | $AUC_{0\text{-}\infty}$ (μg・h/mL) | $CL_{tot}$ (mL/min/kg) | $V_d$ (L/kg) |
|---|---|---|---|---|---|---|---|
| Marmoset 1 | 39 | 365 | 0.832 | 48.57 | 48.85 | 3.4 | 0.205 |
| Marmoset 2 | 39 | 370 | 0.979 | 75.13 | 76.08 | 2.2 | 0.169 |
| Marmoset 3 | 39 | 365 | 1.105 | 34.47 | 34.87 | 4.8 | 0.271 |
| Average | | | 0.972 | 52.73 | 53.27 | 3.5 | 0.215 |
| s.d. | | | 0.137 | 20.65 | 20.96 | 1.3 | 0.052 |

| SN38 | Age (Month) | Body weight (g) | $t_{1/2}$ (h) | $AUC_{0\text{-}6h}$ (μg・h/mL) | $AUC_{0\text{-}\infty}$ (μg・h/mL) | $CL_{tot}$ (mL/min/kg) | $V_d$ (L/kg) |
|---|---|---|---|---|---|---|---|
| Marmoset 1 | 39 | 365 | 1.138 | 1.124 | 1.140 | | |
| Marmoset 2 | 39 | 370 | 1.818 | 0.637 | 0.677 | | |
| Marmoset 3 | 39 | 365 | 1.301 | 0.698 | 0.712 | | |
| Average | | | 1.419 | 0.820 | 0.843 | | |
| s.d. | | | 0.355 | 0.265 | 0.258 | | |

Pharmacokinetic parameters of SN38-βGlcNAc and released SN38.

| SN38-βGlcNAc | Age (Month) | Body weight (g) | $t_{1/2}$ (h) | $AUC_{0\text{-}6h}$ (μg・h/mL) | $AUC_{0\text{-}\infty}$ (μg・h/mL) | $CL_{tot}$ (mL/min/kg) | $V_d$ (L/kg) |
|---|---|---|---|---|---|---|---|
| Marmoset 4 | 39 | 400 | 0.732 | 73.51 | 73.76 | 2.3 | 0.149 |
| Marmoset 5 | 39 | 350 | 0.828 | 84.34 | 84.81 | 2.0 | 0.134 |
| Marmoset 6 | 39 | 365 | 0.821 | 76.92 | 77.34 | 2.2 | 0.143 |
| Average | | | 0.794 | 78.26 | 78.64 | 2.2 | 0.142 |
| s.d. | | | 0.054 | 5.54 | 5.64 | 0.2 | 0.008 |

| SN38 | Age (Month) | Body weight (g) | $t_{1/2}$ (h) | $AUC_{0\text{-}6h}$ (μg・h/mL) | $AUC_{0\text{-}\infty}$ (μg・h/mL) | $CL_{tot}$ (mL/min/kg) | $V_d$ (L/kg) |
|---|---|---|---|---|---|---|---|
| Marmoset 4 | 39 | 400 | 1.543 | 0.987 | 1.032 | | |
| Marmoset 5 | 39 | 350 | 1.514 | 1.061 | 1.107 | | |
| Marmoset 6 | 39 | 365 | 1.614 | 0.680 | 0.717 | | |
| Average | | | 1.557 | 0.909 | 0.952 | | |
| s.d. | | | 0.051 | 0.202 | 0.207 | | |

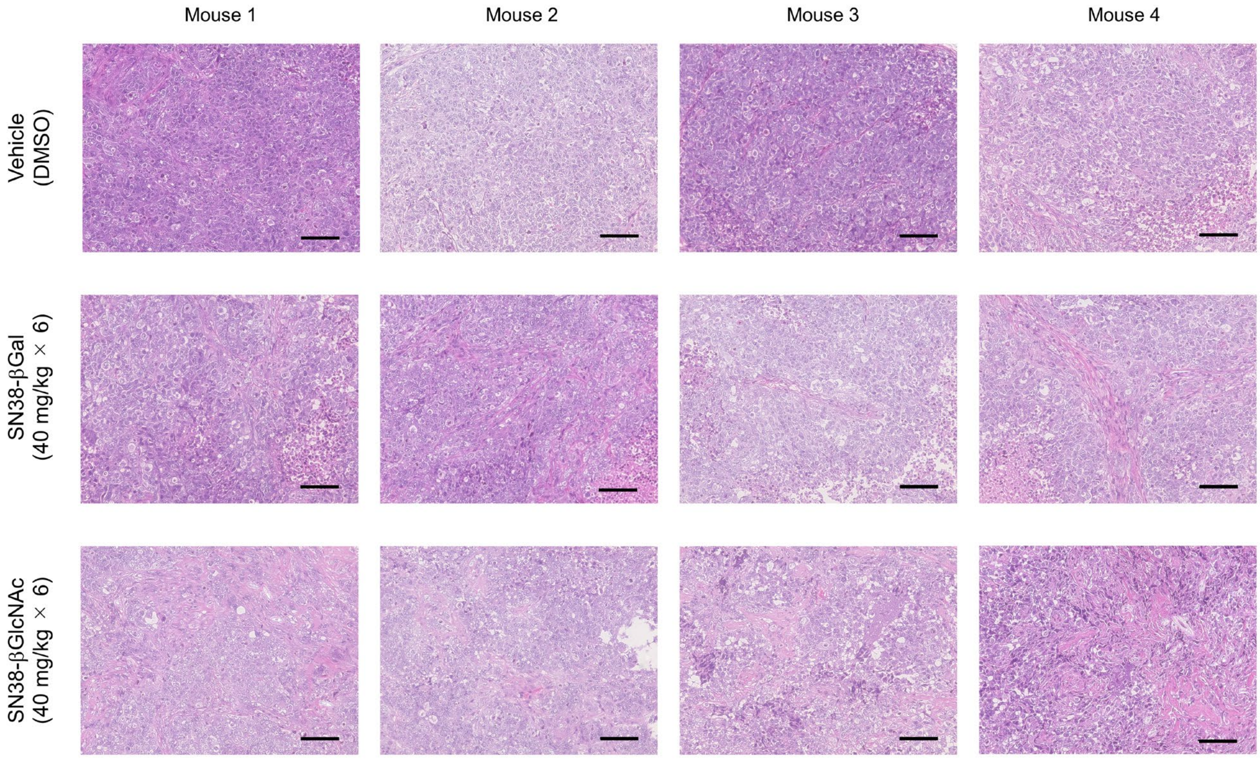


**Figure S18. Histological analysis of resected PDX tumors after treatment with SN38-βGal or SN38-βGlcNAc.** The SN38-βGal and SN38-βGlcNAc (40 mg/kg × 6)-treated groups in **Fig. 5l** exhibited chemotherapeutic damage such as cytoplasmic and nuclear swelling and fibrosis. Scale bars, 100 μm.

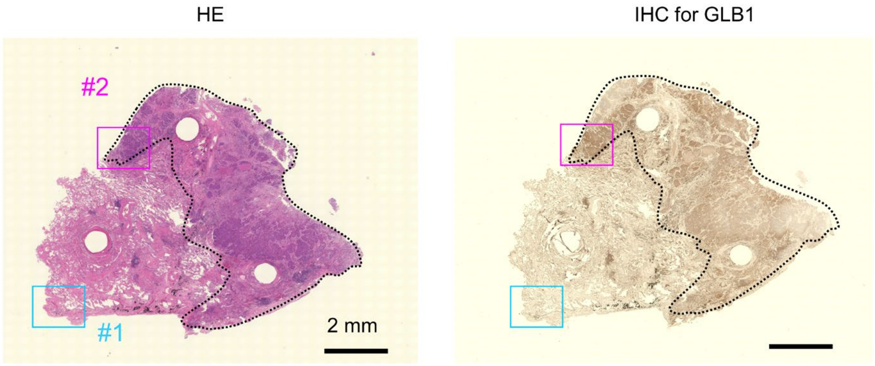


**Figure S19. Histological and IHC analysis for GLB1 in the clinical specimen evaluated by GLB1-reactive fluorescence probe.** Histological and IHC images of the evaluated specimen from **Extended Fig. 8a**. Cancer tissues (enclosed with dotted line) showed overexpression of GLB1. Scale bars, 2 mm.

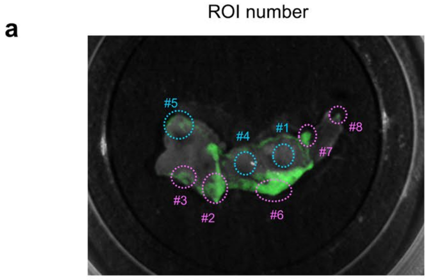


**b**

| ROI | Fluorescence increase (a.u.) | Cancer existence | HEXA level | HEXB level |
|---|---|---|---|---|
| #1 | 0.287 | - | - | - |
| #2 | 0.586 | + | +++ | +++ |
| #3 | 0.464 | + | +++ | +++ |
| #4 | 0.241 | - | - (+ in macrophage) | - (+ in macrophage) |
| #5 | 0.363 | - | - (+ in macrophage) | - (+ in macrophage) |
| #6 | 0.587 | + | +++ | +++ |
| #7 | 0.544 | + | ++ | +++ |
| #8 | 0.489 | + | + | + |

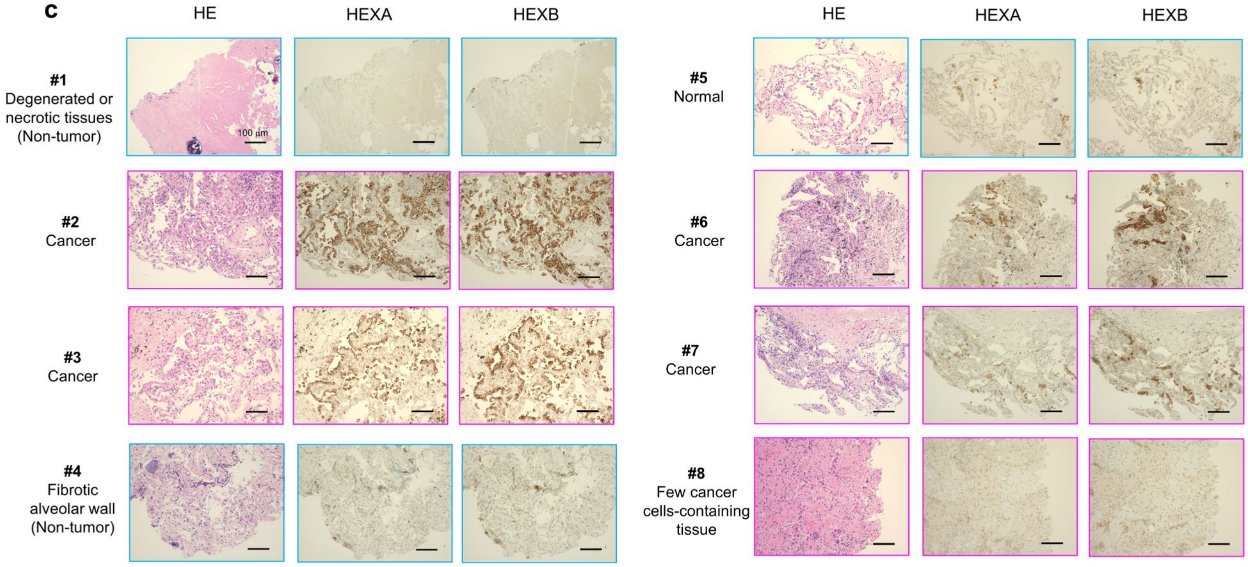


**Figure S20. Histological and IHC analysis for HEXA and HEXB in the clinical specimen evaluated by HEX-reactive fluorescence probe.** Histological and IHC images of the evaluated specimen from **Extended Fig. 8b**. (**a**) Evaluated ROI number, #1-8. (**b**) Result of pathological analysis of each ROI. Cancer tissues showed overexpression of HEXA and HEXB. Normal tissues showed slight expression of HEXA and HEXB in macrophages. (**c**) Histological and IHC analyses of fluorescent and non-fluorescent regions in Case 2. Scale bars, 100 µm.

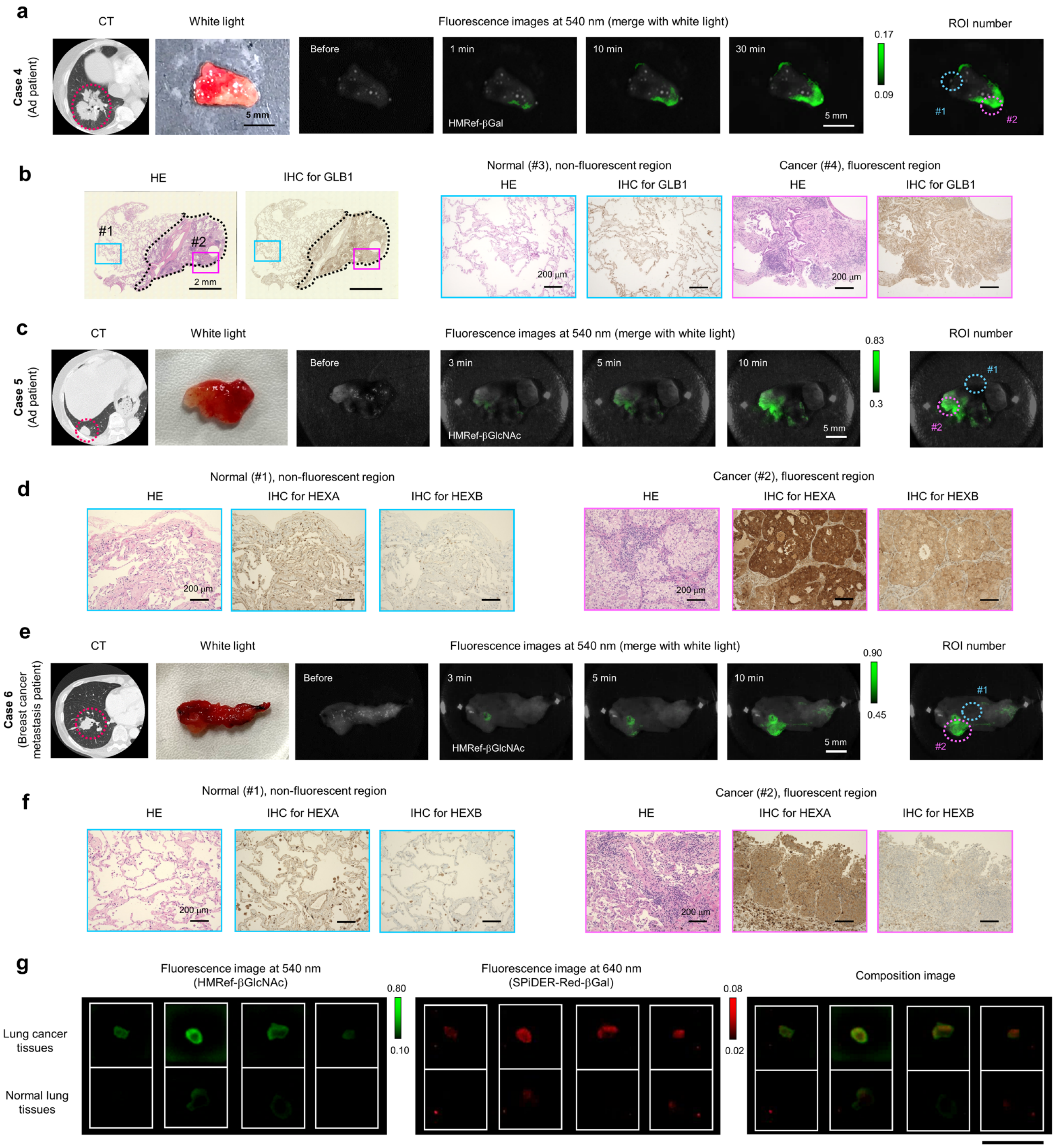


**Figure S21. Rapid and sensitive visualization of target glycosidase activities in clinical lung cancer specimens using fluorescence probes.** (**a**) CT image of Ad patient (left), corresponding white light image (middle) and time-dependent fluorescence images at 540 nm (right) of surgically resected fresh lung specimens containing both normal and Ad tissues after administration of HMRef-βGal (Case 4). [HMRef-βGal] = 50 μM. Scale bars, 5 mm. (**b**) Histological analysis and IHC analysis of GLB1 in boxed regions with no fluorescence activation (blue box, #1) or with strong fluorescence activation (pink box, #2) in (**a**). Fluorescent regions were well matched with pathologically identified cancer regions and GLB1-overexpressing regions. Scale bars, 200 μm. (**c**) CT image of Ad patient (left), corresponding white light image (middle) and time-dependent fluorescence images at 540 nm of surgically resected fresh lung specimens containing both normal and Ad tissues after administration of HMRef-βGlcNAc (Case 5). [HMRef-βGlcNAc] = 50 μM. Scale bar, 5 mm. (**d**) Histological analysis and IHC analyses for HEXA and HEXB in boxed regions with no fluorescence activation (blue box, #1) or with strong fluorescence activation (pink box, #2) in (**c**). Fluorescence regions were well matched with pathologically confirmed Ad regions and HEXA and HEXB-overexpressing regions. Scale bars, 200 μm. (**e**) CT image of breast cancer lung metastasis patient (left), corresponding white light image (middle) and time-dependent fluorescence images at 540 nm (right) of surgically resected fresh lung specimens containing both normal and cancer tissues after administration of HMRef-βGlcNAc (Case 6). Scale bar, 5 mm. [HMRef-βGlcNAc] = 50 μM. (**f**) Histological analysis and IHC analysis of HEXA and HEXB in boxed regions with no fluorescence activation (blue box, #1) or with strong fluorescence activation (pink box, #2) in (**e**). Fluorescence regions were well matched with pathologically confirmed cancer regions and HEXA and HEXB-overexpressing regions. Scale bars, 200 μm. (**f**) Fluorescence images at 540 nm and 640 nm (right) of pairs of surgically resected normal lung and lung cancer specimens from 4 different patients. Fluorescence images were captured 30 min after administration of HMRef-βGlcNAc and 2'Me-SPiDER-Red-βGal. [HMRef-βGlcNAc] = 50 μM, [2'Me-SPiDER-Red-βGal] = 50 μM. Scale bar, 1 cm.

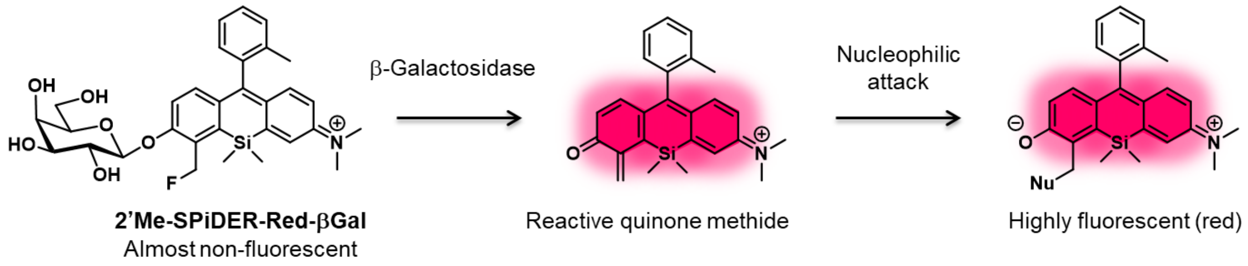


**Figure S22. Reaction scheme for 2'Me-SPiDER-Red-βGal.** Red fluorescence probe 2'Me-SPiDER-Red-βGal (2'Me-4$CH_2$F-Sirhodol-βGal) reacts with β-galactosidase and immediately produces the reactive quinone methide intermediate, which reacts with intracellular nucleophiles such as GSH, $H_2O$ and proteins.[15]

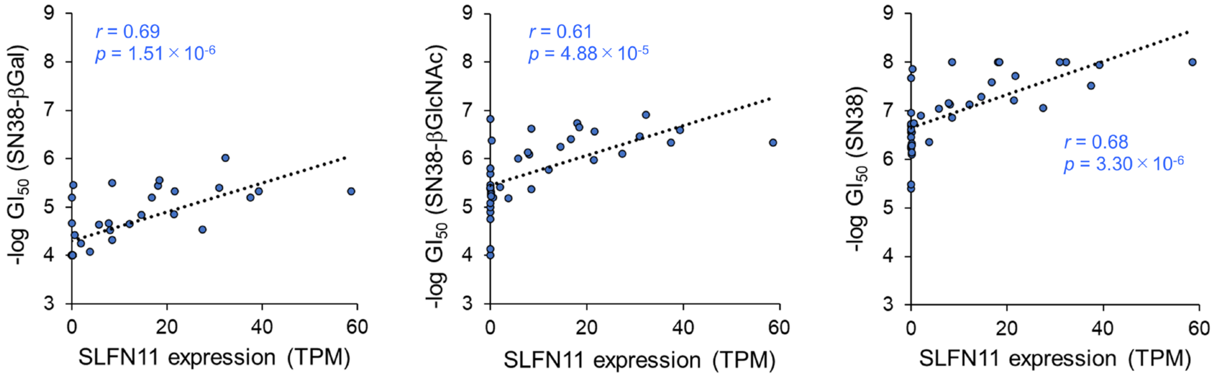


**Figure S23. Correlations between -log $GI_{50}$ of developed prodrugs and SLFN11 expression.** Correlations between SN38-βGal (left), SN38-βGlcNAc (middle) or SN38 (right) and SLFN11 expression (TPM) in JFCR39 cancer cell panel. A good correlation was found for each compound. The LOX-IMVI was excluded from the analyses of SN38-βGal and SN38-βGlcNAc.

**Procedure for the synthesis of glycosidase-reactive probes based on HMRef scaffold**. Probes 1-12 were synthesized according to the reported procedures.[14,16]

**Procedure for the synthesis of 2'Me-SPiDER-Red-βGal**. 2'Me-SPiDER-Red-βGal was synthesized according to the reported procedure.[15]

**Procedure for the synthesis of aminopeptidase-reactive probes based on the HMRG scaffold**. gGlu-HMRG was synthesized according to the reported procedure.[17] GP-HMRG, PP-HMRG, KR-HMRG and KA-HMRG were synthesized according to the reported procedures.[6] EP-HMRG was synthesized according to the reported procedure.[9,18] ZFR-HMRG was synthesized according to the reported procedure.[13]

**Scheme 1.** Synthesis of Boc-Lys(Boc)Lys(Boc)-OMe.

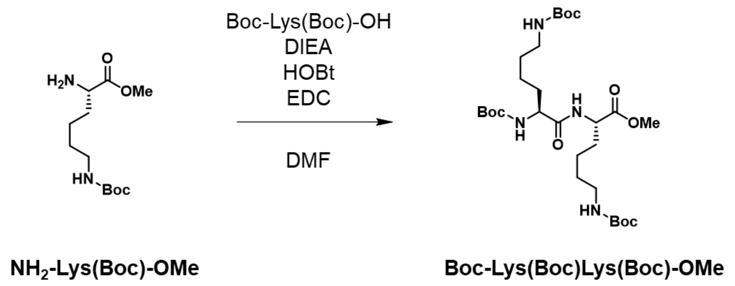


**Procedure for the synthesis of Boc-Lys(Boc)Lys(Boc)-OMe.** To a mixture of Boc-Lys(Boc)-OH (355 mg, 1.02 mmol), and anhydrous *N*,*N*-dimethylformamide (5.0 mL) were added diisopropylethylamine (389 mg, 3.01 mmol), $NH_2$-Lys(Boc)-OMe HCl (304 mg, 1.02 mmol), anhydrous HOBt (162 mg, 1.20 mmol), and EDC (235 mg, 1.23 mmol) at room temperature under an argon atmosphere. The mixture was stirred for 17.5 h at the same temperature, the concentrated under reduced pressure. Dichloromethane (30.0 mL) was added and the mixture was washed with 5 % citric acid solution (3 × 20.0 mL), aqueous saturated sodium hydrogen carbonate (3 × 20.0 mL) and brine, dried over sodium sulfate and filtered. The filtrate was concentrated under reduced pressure to give **Boc-Lys(Boc)Lys(Boc)-OMe** (382 mg, 0.649 mmol) as a colorless oil in 63.6 % yield. The identity of the product was confirmed by means of $^1H$ NMR and ESI-HRMS. $^1H$ NMR (400 MHz, $CD_3OD$) δ 4.39 (dd, *J* = 8.9, 4.8 Hz, 1H), 4.03 (dd, *J* = 8.2, 5.5 Hz, 1H), 3.70 (s, 3H), 3.05–3.00 (m, 4H), 1.89–1.55 (m, 4H), 1.50–1.38 (m, 35H). ESI-HRMS($ESI^+$): calcd for $[M+Na]^+$, 611.3626; found, 611.3643.

**Scheme 2.** Synthesis of Boc-Lys(Boc)Lys(Boc)-OH.

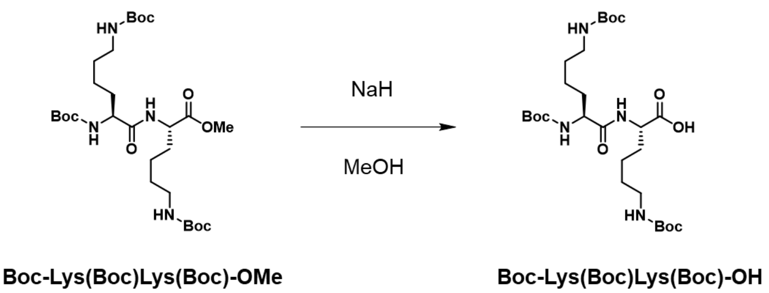


**Procedure for the synthesis of Boc-Lys(Boc)Lys(Boc)-OH.** To a mixture of Boc-Lys(Boc)Lys(Boc)-OMe (374 mg, 0.635 mmol) and methanol (4.0 mL) was added 1 M sodium hydroxide solution (1.5 mL) at room temperature. The mixture was stirred for 2.5 h at the same temperature, then 0.1 M hydrochloric acid solution (16.0 mL) was added. The mixture was extracted with dichloromethane (3 × 20.0 mL). The combined organic extract was washed with brine, dried over sodium sulfate and filtered. The filtrate was concentrated under reduced pressure to give **Boc-Lys(Boc)Lys(Boc)-OH** (195 mg, 0.339 mmol) as a colorless solid in 53.3 % yield. The identity of the product was confirmed by means of $^1$H NMR and ESI-HRMS. $^1$H NMR (400 MHz, $CD_3OD$) δ 4.35 (dd, *J* = 8.5, 4.8 Hz, 1H), 4.03 (dd, *J* = 8.2, 5.0 Hz, 1H), 3.02 (t, *J* = 6.6 Hz, 4H), 1.91–1.55 (m, 4H), 1.52–1.38 (m, 35H). ESI-HRMS(ESI$^+$): calcd for $[M+Na]^+$, 597.3470; found, 597.3496.

**Scheme 3.** Synthesis of KK-HMRG.

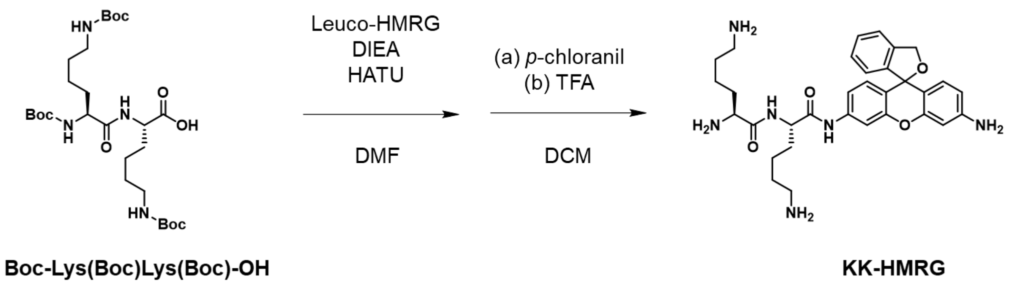


**Procedure for the synthesis of KK-HMRG.** To a mixture of Leuco-HMRG (40.0 mg, 0.13 mmol), Boc-Lys(Boc)Lys(Boc)-OH (80.0 mg, 0.14 mmol) and anhydrous *N*,*N*-dimethylformamide (1.0 mL) were added diisopropylethylamine (48.0 mg, 0.37 mmol), and HATU (53.0 mg, 0.14 mmol) at room temperature under an argon atmosphere. The mixture was stirred for 4 h at the same temperature, then water (2.0 mL) was added and the mixture was extracted with ethyl acetate. The combined organic extract was washed with water and brine, dried over sodium sulfate and filtered. The filtrate was concentrated under reduced pressure. To a mixture of the crude product and dichloromethane (2.0 mL) was added *p*-chloranil (34.0 mg, 0.14 mmol) at room temperature. The mixture was stirred for 50 min at the same temperature, then trifluoroacetic acid (2.0 mL) was added and stirring was continued for 1.5 h at the same temperature. The mixture was concentrated under reduced pressure. The residue was purified by HPLC to afford **KK-HMRG** (22.8 mg, 0.0332 mmol) as the trifluoroacetate salt as a red solid in 25.6 % yield. The identity of the product was confirmed by means of $^1$H NMR and ESI-HRMS. $^1$H NMR (400 MHz, $CD_3OD$ + $K_2CO_3$) δ 7.62–7.60 (m, 1H), 7.43–7.36 (m, 2H), 7.27 (dd, *J* = 6.9, 6.9 Hz, 1H), 7.11 (dd, *J* = 8.7, 2.3 Hz, 1H), 6.85–6.80 (m, 2H), 6.64 (d, *J* = 8.7 Hz, 1H), 6.50 (d, *J* = 2.3 Hz, 1H), 6.41 (dd, *J* = 8.5, 2.1 Hz, 1H), 5.25 (s, 2H), 4.47–4.43 (m, 1H), 3.37 (t, *J* = 6.6 Hz, 1H), 2.66–2.57 (m, 4H), 1.92–1.32 (m, 12H). ESI-HRMS($ESI^+$): calcd for $[M+H]^+$, 573.3184; found, 573.3204.

**Scheme 4.** Synthesis of SN38-βGal.

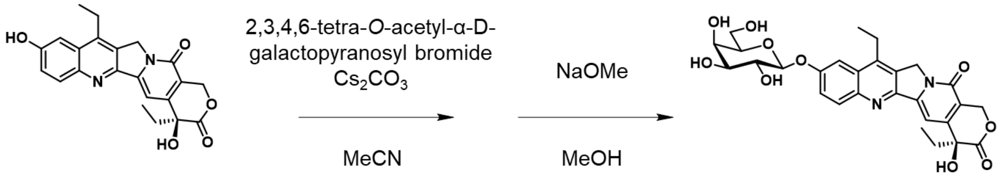


**SN38** **SN38-βGal**

**Procedure for the synthesis of SN38-βGal.** A mixture of SN38 (1.50 g, 3.82 mmol), 2,3,4,6-tetra-*O*-acetyl-α-D-galactopyranosyl bromide (7.00 g, 17.0 mmol), and $Cs_2CO_3$ (5.00 g, 15.3 mmol) in 80.0 mL dried MeCN was stirred at room temperature for 3 h, then concentrated under reduced pressure. The residue was purified by flash column chromatography ($CH_2Cl_2$ : MeOH = 95 : 5) to obtain the crude 2,3,4,6-tetra-*O*-acetyl-β-D-galactopyranosylated derivative, which was dissolved in 70.0 mL dried MeOH. To this solution, NaOMe (1.50 g, 27.8 mmol) dissolved in 30.0 mL dried MeOH was added dropwise, and the reaction mixture was stirred at room temperature for 1 h then concentrated under reduced pressure. The residue was purified by HPLC to afford **SN38-βGal** (1.10 g, 1.99 mmol) as a yellow powder in 52.1 % yield. The identity of the product was confirmed by means of $^1$H NMR, $^{13}$C NMR and ESI-HRMS. $^1$H NMR (400 MHz, DMSO-d6): δ 8.10 (d, *J* = 9.6 Hz, 1H), 7.74 (s, 1H), 7.58 (dd, *J* = 1.2 Hz, 9.2 Hz, 1H), 7.28 (s, 1H), 5.43 (s, 2H), 5.30 (s, 2H), 5.10 (d, *J* = 7.6 Hz, 1H, overlaps with HOD), 3.75-3.68 (m, 3H), 3.60-3.20 (m, 2H), 3.50 (dd, *J* = 2.4 Hz, 9.2 Hz, 1H), 3.21-3.20 (m, 2H), 1.95-1.80 (m, 2H), 1.31 (t, *J* = 7.2 Hz, 3H), 0.88 (t, *J* = 7.2 Hz, 3H). $^{13}$C NMR (101 MHz, DMSO-d6): δ 172.5, 156.8, 156.1, 150.1, 150.0, 146.2, 144.7, 144.2, 131.2, 128.3, 127.6, 122.5, 118.4, 106.6, 101.1, 96.1, 75.9, 73.3, 72.4, 70.2, 68.2, 65.2, 60.5, 49.5, 30.2, 22.2, 13.5, 7.7. ESI-HRMS (ESI +) m/z calcd. for $[M+Na]^+$, 539.05247; found, 539.05233.

**Scheme 5.** Synthesis of SN38-βGlcNAc.

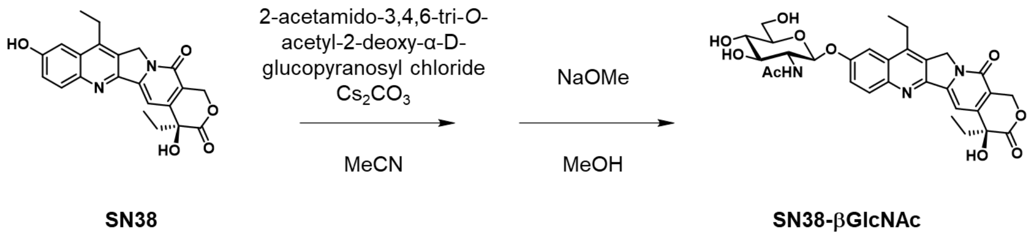


**Procedure for the synthesis of SN38-βGlcNAc.** A mixture of SN38 (1.00 g, 2.55 mmol), 2-acetamido-3,4,6-tri-*O*-acetyl-2-deoxy-α-D-glucopyranosyl chloride (4.00 g, 10.9 mmol), and $Cs_2CO_3$ (3.20 g, 9.82 mmol) in 100 mL dried MeCN was stirred at room temperature for 3 h, then concentrated under reduced pressure. The residue was purified by flash column chromatography ($CH_2Cl_2$ : MeOH = 95 : 5) to obtain the crude 2-acetamido-3,4,6-tri-*O*-acetyl-2-deoxy-β-D-glucopyranosylated derivative, which was dissolved in 60.0 mL dried MeOH. To this solution, NaOMe (700 mg, 13.0 mmol) dissolved in 20.0 mL dried MeOH was added dropwise, and the reaction mixture was stirred at room temperature for 1 h then concentrated under reduced pressure. The residue was purified by HPLC to afford **SN38-βGlcNAc** (947 mg, 1.59 mmol) as a yellow powder in 62.6 % yield. The identity of the product was confirmed by means of $^1H$ NMR, $^{13}C$ NMR and ESI-HRMS. $^1H$ NMR (400 MHz, $CD_3OD$): δ 7.57 (d, *J* = 9.2 Hz, 1H), 7.38 (s, 1H), 7.24 (s, 1H), 7.20 (d, *J* = 2.4 Hz, 1H), 5.39 (d, *J* = 16 Hz, 1H), 5.17 (d, *J* = 16 Hz, 1H), 5.13 (d, *J* = 8.4 Hz, 1H), 3.99-3.93 (m, 2H), 3.75 (dd, *J* = 6.8 Hz, 12 Hz, 1H), 3.62 (t, *J* = 8.8 Hz, 1H), 3.55-3.51 (m, 1H), 3.39 (t, *J* = 9.2 Hz, 1H), 2.98-2.94 (m, 2H), 1.98 (s, 3H), 1.86-1.78 (m, 2H), 1.26 (t, *J* = 7.6 Hz, 3H), 0.90 (t, *J* = 7.6 Hz, 3H). $^{13}C$ NMR (101 MHz, $CD_3OD$): δ 174.8, 174.0, 158.8, 157.7, 152.6, 160.6, 147.4, 146.5, 145.8, 131.8, 129.0, 128.7, 123.6, 119.8, 108.1, 100.8, 99.1, 79.0, 75.8, 74.1, 72.2, 66.6, 62.8, 57.4, 40.4, 32.2, 23.9, 23.2, 14.1, 8.2. ESI-HRMS (ESI +) m/z calcd. for $[M+Na]^+$, 618.20582; found, 618.20633.

**Scheme 6.** Synthesis of compound 1.

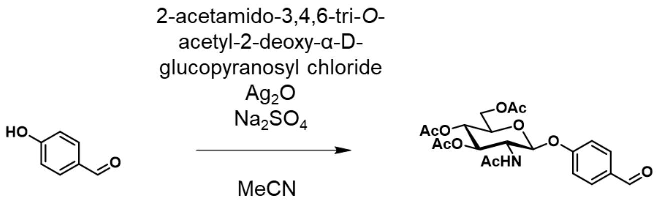


***p*-hydroxybenzaldehyde** **1**

**Procedure for the synthesis of compound 1.** A mixture of *p*-hydroxybenzaldehyde (3.00 g, 24.6 mmol), 2-acetamido-3,4,6-tri-O-acetyl-2-deoxy-α-D-glucopyranosyl chloride (9.00 g, 24.6 mmol) and $Ag_2O$ (17.0 g, 73.4 mmol) in 80 mL dried MeCN was stirred at room temperature for 20 h. The reaction mixture was filtered through a Celite pad, then concentrated under reduced pressure. The residue was purified by flash column chromatography ($CH_2Cl_2$ : MeOH = 100 : 0 to 70 : 30) to afford compound **1** (10.53 g, 23.3 mmol) as a white powder in 95.0 % yield. The identity of the product was confirmed by means of $^1H$ NMR, $^{13}C$ NMR and ESI-HRMS. $^1H$ NMR (400 MHz, MeOD): δ 9.90 (s, 1H), 7.91 (d, *J* = 8.8 Hz, 2H), 7.21 (d, *J* = 8.8 Hz, 2H), 5.49 (d, *J* = 8.4 Hz, 1H), 5.39 (t, *J* = 9.2 Hz, 1H), 5.11 (t, *J* = 10 Hz, 1H), 4.34 (dd, *J* = 5.2 Hz, 12.4 Hz, 1H), 4.20-4.10 (m, 3H), 2.07 (s, 3H), 2.06 (s, 3H), 2.04 (s, 3H), 1.93 (s, 3H). $^{13}C$ NMR (101 MHz, MeOD): δ 192.8, 173.7, 172.3, 171.8, 171.3, 163.2, 133.1, 132.9, 117.8, 98.9, 73.7, 73.3, 69.9, 63.1, 55.4, 22.7, 20.6, 20.6, 20.5. ESI-HRMS (ESI +) m/z calcd. for $[M+Na]^+$, 474.13707; found, 474.13640.

**Scheme 7.** Synthesis of compound **2**.

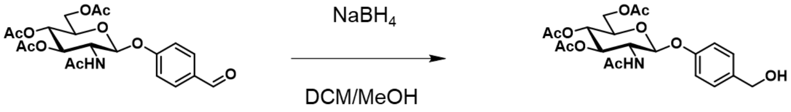


**1** **2**

**Procedure for the synthesis of compound 2.** A mixture of compound 1 (750 mg, 1.66 mmol) and $NaBH_4$ (50.0 mg, 1.32 mmol) in 10 mL solution ($CH_2Cl_2$ : MeOH = 1:1) was stirred at room temperature for 1.5 h. After addition of water and $CH_2Cl_2$, the organic layer was washed with water and brine, dried over $Na_2SO_4$ and concentrated to afford compound **2** (665 mg, 1.46 mmol) as a white powder in 88.3 % yield. The identity of the product was confirmed by means of $^1H$ NMR and ESI-HRMS. $^1H$ NMR (400 MHz, $CD_2Cl_2$): δ 7.33 (d, *J* = 8.4 Hz, 2H), 7.02 (d, *J* = 8.4 Hz, 2H), 5.70 (d, *J* = 8.4 Hz, 1H), 5.43 (t, *J* = 10 Hz, 1H), 5.31 (d, *J* = 8.0 Hz, 1H), 5.13 (t, *J* = 9.6 Hz, 1H), 4.30 (dd,

*J* = 5.6 Hz, 12.0 Hz, 1H), 4.18-4.00 (m, 2H), 3.95-3.85 (m, 1H), 2.09-2.07 (m, 9H), 1.95 (s, 3H). ESI-HRMS (ESI +) m/z calcd. for $[M+Na]^+$, 476.15272; found, 476.15320.

**Scheme 8.** Synthesis of compound **3**.

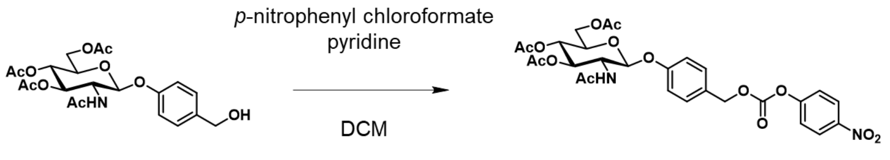


2 3

**Procedure for the synthesis of compound 3.** A mixture of compound 2 (278 mg, 0.614 mmol), *p*-nitrophenyl chloroformate (368 mg, 1.84 mmol) and pyridine (148 μL) in 30 mL dried DCM was stirred at room temperature under an Ar atmosphere for 3 h. The organic layer was washed with sat $NaHCO_3$ aq., water, and brine, dried over $Na_2SO_4$ and concentrated. The residue was purified by flash column chromatography (AcOEt : hexane = 0 : 100 to 100 : 0) to afford compound **3** (267 mg, 0.424 mmol) as a white powder in 69.1 % yield. The identity of the product was confirmed by means of $^1$H NMR and ESI-HRMS. $^1$H NMR (400 MHz, $CD_2Cl_2$): δ 8.30 (d, *J* = 9.2 Hz, 2H), 7.44-7.41 (m, 4H), 7.07 (d, *J* = 8.8 Hz, 2H), 5.78 (d, *J* = 8.8 Hz, 1H), 5.48 (t, *J* = 9.6 Hz, 1H), 5.39 (d, *J* = 8.4 Hz, 1H), 5.27 (s, 2H), 5.14 (t, *J* = 9.6 Hz, 1H), 4.31 (dd, *J* = 5.6 Hz, 12.4 Hz, 1H), 4.19-4.07 (m, 2H), 3.97-3.92 (m, 1H), 2.08-2.07 (m, 9H), 1.95 (s, 3H). ESI-HRMS (ESI +) m/z calcd. for $[M+Na]^+$, 641.15892; found, 641.16136.

**Scheme 9.** Synthesis of DOX-βGlcNAc.

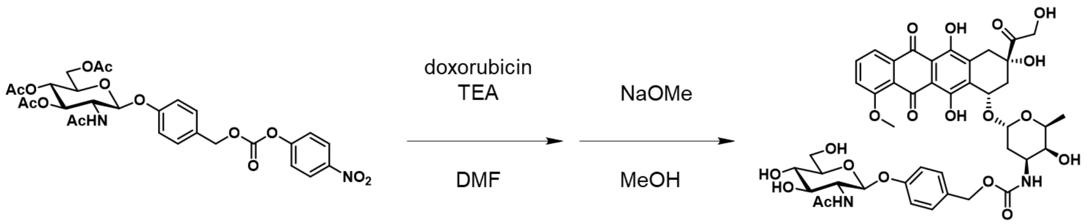


3 **DOX-βGlcNAc**

**Procedure for the synthesis of DOX-βGlcNAc.** A mixture of compound **3** (170 mg, 0.275 mmol), doxorubicin (150 mg, 0.276 mmol), triethylamine (600 μL) in 8.0 mL dried DMF was stirred at room temperature under an Ar atmosphere for 1 h, then concentrated under reduced pressure. After addition of $Et_2O$, the red precipitate was collected by filtration. This red solid was dissolved in 20.0 mL dried MeOH. To this solution, NaOMe (149 mg, 2.76 mmol) was added, and the reaction mixture was stirred at room temperature for 1 h. After quenching with

Amberlite IR120 H (300 mg), the solution was filtered and concentrated under reduced pressure. The residue was purified by flash column chromatography ($CH_2Cl_2$ : MeOH = 100 : 0 to 75 : 25) to afford **DOX-βGlcNAc** (69.0 mg, 0.0770 mmol) as a red powder in 30.0 % yield. The identity of the product was confirmed by means of $^1$H NMR, $^{13}$C NMR and ESI-HRMS. $^1$H NMR (400 MHz, MeOD): δ 7.83 (d, *J* = 7.2 Hz, 1H), 7.75 (t, *J* = 8.0 Hz, 1H), 7.47 (d, *J* = 8.0 Hz, 1H), 7.22 (d, *J* = 8.4 Hz, 2H), 6.93 (d, *J* = 8.4 Hz, 2H), 5.38 (s, 1H), 5.20-4.94 (m, overlaps with HOD), 4.74 (s, 1H), 4.31-4.24 (m, 1H), 3.98 (s, 3H), 3.90-3.81 (m, 4H), 3.74-3.51 (m, 9H), 4.77 (d, *J* = 4.4 Hz, 3H), 3.08-3.03 (m, 1H), 2.93-2.87 (m, 1H), 2.37-2.33 (m, 1H), 2.15-2.13 (m, 1H), 2.05-1.94 (m, 4H), 1.76-1.73 (m, 1H), 1.26 (d, *J* = 6.4 Hz, 3H). $^{13}$C NMR (101 MHz, MeOD): δ 214.0, 187.8, 187.5, 173.9, 162.3, 158.9, 158.0, 157.3, 156.1, 137.1, 136.2, 135.6, 135.0, 132.3, 130.6, 129.5, 121.3, 120.5, 120.2, 117.6, 112.3, 112.1, 102.2, 100.8, 78.2, 77.4, 75.8, 71.9, 71.8, 70.2, 68.7, 67.1, 65.8, 62.6, 57.4, 57.1, 37.3, 34.0, 30.9, 23.0, 17.3. ESI-HRMS (ESI +) m/z calcd. for $[M+Na]^+$, 919.27435; found, 919.27385.

**Scheme 10.** Synthesis of MMAE-βGlcNAc.

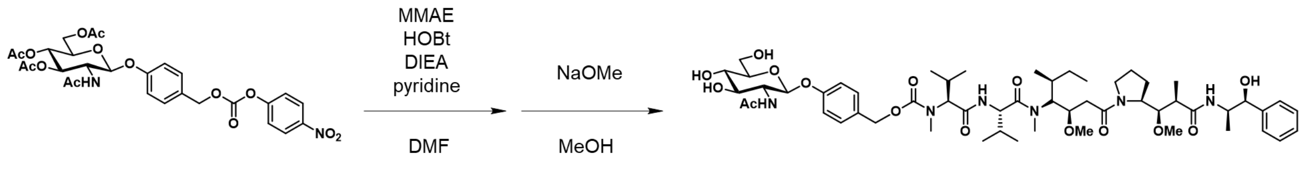


**Procedure for the synthesis of MMAE-βGlcNAc.** A mixture of compound **3** (70.0 mg, 0.113 mmol), monomethyl auristatin E (MMAE) (79.7 mg, 0.113 mmol), 1-hydroxybenzotriazole (HOBt) (15.3 mg, 0.113 mmol), *N,N*-diisopropylethylamine (DIEA) (30 μL), pyridine (1.5 mL) in 6.0 mL dried DMF was stirred at room temperature under an Ar atmosphere for 20 h, then concentrated under reduced pressure. The residue was purified by HPLC to obtain the crude 2-acetamido-3,4,6-tri-*O*-acetyl-2-deoxy-β-D-glucopyranosylated derivative, which was dissolved in 10.0 mL dried MeOH. To this solution, NaOMe (61.0 mg, 1.13 mmol) was added, and the reaction mixture was stirred at room temperature for 1 h. After quenching with Amberlite IR120 H (300 mg), the solution was filtered and concentrated under reduced pressure. The residue was purified by HPLC to afford the **MMAE-βGlcNAc** (115 mg, 0.107 mmol) as a colorless solid in 94.9 % yield. The identity of the product was confirmed by means of $^1$H NMR and ESI-HRMS. $^1$H NMR (400 MHz, MeOD): δ 7.43-7.30 (m, 6H), 7.25-7.21 (m, 1H), 7.04 (d, *J* = 8.0 Hz, 2H), 5.60-5.40 (m, overlaps with HOD), 5.19-5.07 (m, overlaps with HOD), 4.72-4.66 (m, 1H), 4.65-4.58 (m, 1H), 4.28-4.21 (m, 3H), 4.150-4.09 (m, 1H), 3.98-3.89 (m, 3H), 3.80-3.74 (m, 2H), 3.67-3.62 (m, 1H), 3.60-3.50 (m, 1H), 3.48 (d, *J* = 5.2 Hz, 3H), 3.43-3.29 (m, 10H), 3.25-3.15 (m, 1H), 3.12 (s, 1H), 2.99-2.92 (m, 3H), 2.53-2.48 (m, 2H), 2.30-2.10 (m, 3H), 2.00 (d, *J* = 4.0 Hz, 4H), 1.91-1.84 (m, 2H), 1.75-1.71 (m, 1H), 1.64-1.59 (m, 1H), 1.50-1.35

(m, 2H), 1.21-1.14 (m, 6H), 1.03-0.75 (m, 19H). ESI-HRMS (ESI +) m/z calcd. for $[M+Na]^+$, 1093.60434; found, 1093.60145.

**Scheme 11.** Synthesis of MEL-βGlcNAc.

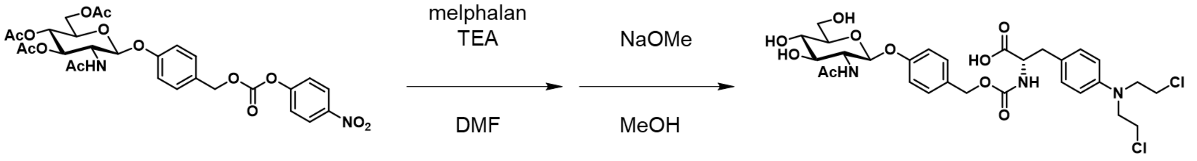


3

MEL-βGlcNAc

**Procedure for the synthesis of MEL-βGlcNAc.** A mixture of compound **3** (203 mg, 0.328 mmol), melphalan (100 mg, 0.328 mmol) and triethylamine (400 μL) in 15.0 mL dried DMF was stirred at room temperature under an Ar atmosphere for 1.5 h, then concentrated under reduced pressure. The residue was dissolved in 20.0 mL dried MeOH. To this solution, NaOMe (177 mg, 3.28 mmol) was added, and the reaction mixture was stirred at room temperature for 1 h. After quenching with Amberlite IR120 H (300 mg), the solution was filtered and concentrated under reduced pressure. The residue was purified by HPLC to afford the **MEL-βGlcNAc** (92.6 mg, 0.141 mmol) as a white solid in 43.0 % yield. The identity of the product was confirmed by means of $^1$H NMR and ESI-HRMS. $^1$H NMR (400 MHz, MeOD): δ 7.24 (d, *J* = 8.8 Hz, 2H), 7.08 (d, *J* = 8.4 Hz, 2H), 7.00 (d, *J* = 8.4 Hz, 2H), 6.65 (d, *J* = 8.4 Hz, 2H), 5.06 (d, *J* = 8.4 Hz, 1H), 4.33-4.30 (m, 1H), 3.98-3.91 (m, 2H), 3.76-3.57 (m, 10H), 3.47 (s, 2H), 3.33 (t, *J* = 1.2 Hz, 1H), 3.08 (dd, *J* = 4.8 Hz, *J* = 14 Hz, 1H), 3.08 (dd, *J* = 4.8 Hz, *J* = 14 Hz, 1H), 2.85 (dd, *J* = 8.4 Hz, *J* = 14 Hz, 1H), 2.66 (s, 3H), 2.00 (s, 2H). $^{13}$C NMR (101 MHz, MeOD): δ 176.5, 174.0, 158.8, 158.3, 146.5, 132.3, 131.6, 130.5, 127.3, 118.0, 117.6, 113.3, 100.8, 78.2, 75.8, 71.8, 67.2, 62.5, 57.7, 57.3, 54.5, 41.8, 40.5, 37.9, 23.1. ESI-HRMS (ESI +) m/z calcd. for $[M+Na]^+$, 680.17482; found, 680.17122.

**Scheme 12.** Synthesis of compound **4**.

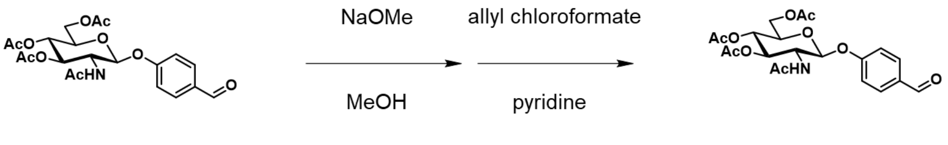


**3** **4**

**Procedure for the synthesis of compound 4.** Compound **1** (5.05 g, 11.2 mmol) was dissolved in 10.0 mL dried MeOH. To this solution, NaOMe (2.00 g, 37.0 mmol) was added, and the reaction mixture was stirred at room temperature for 1 h. After quenching with Amberlite IR120 H (2.00 g), the solution was filtered and concentrated under reduced pressure. The residue was purified by flash column chromatography ($CH_2Cl_2$ : MeOH = 100 : 0 to 0 : 100) to obtain the deprotected derivative, which was dissolved in 15.0 mL pyridine. To this solution, allyl chloroformate (13.0 g, 108 mmol) was added at 0 °C and the mixture was stirred at room temperature under an Ar atmosphere for 20 h. After addition of DCM, the organic layer was washed with sat $NaHCO_3$ aq., water, and brine, dried over $Na_2SO_4$ and concentrated. The residue was purified by flash column chromatography (AcOEt : hexane = 0 : 100 to 100 : 0) to afford compound **4** (144 mg, 0.249 mmol) as a colorless solid in 2.2 % yield. The identity of the product was confirmed by means of $^{1}$H NMR, $^{13}$C NMR and ESI-HRMS. $^{1}$H NMR (400 MHz, $CD_2Cl_2$): δ 8.29 (d, *J* = 9.2 Hz, 2H), 7.41 (d, *J* = 9.2 Hz, 4H), 7.10 (d, *J* = 8.8 Hz, 2H), 6.33 (d, *J* = 8.4 Hz, 1H), 6.01-5.91 (m, 3H), 5.59 (d, *J* = 8.4 Hz, 1H), 5.52 (t, *J* = 9.2 Hz, 1H), 5.40-5.26 (m, 8H), 5.01 (t, *J* = 10 Hz, 1H), 4.67-4.46 (m, 4H), 4.62 (d, *J* = 5.6 Hz, 1H), 4.43-4.39 (m, 1H), 4.35-4.31 (m, 1H), 4.15-4.00 (m, 2H),1.94 (s, 3H). $^{13}$C NMR (101 MHz, $CD_2Cl_2$): δ 171.1, 157.8, 156.0, 155.1, 154.9, 154.3, 152.8, 145.8, 132.0, 131.7, 131.7, 130.8, 129.5, 125.6, 122.3, 119.4, 119.2, 119.0, 117.4, 98.3, 75.9, 73.0, 71.8, 70.9, 69.6, 69.4, 69.1, 65.9, 55.5, 23.5. ESI-HRMS (ESI +) m/z calcd. for $[M+Na]^+$, 767.19062; found, 767.19413.

**Scheme 13.** Synthesis of compound **5**.

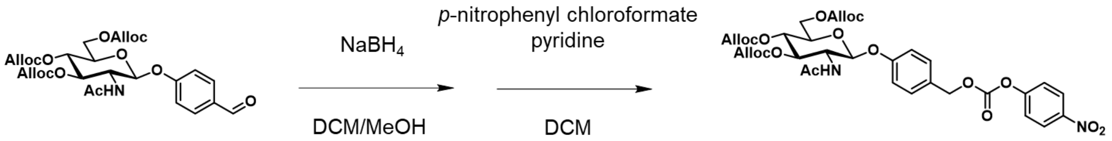


4

5

**Procedure for the synthesis of compound 5.** A mixture of compound **4** (144 mg, 0,249 mmol) and $NaBH_4$ (7.50 mg, 0.198 mmol) in 3.0 mL dried DCM was stirred at room temperature under an Ar atmosphere for 30 min. After addition of water, the mixture was extracted with DCM. The organic layer was washed with brine, dried over $Na_2SO_4$ and concentrated. The residue was dissolved in 20.0 mL dried DCM, and *p*-nitrophenyl chloroformate (150 mg, 0.750 mmol) and pyridine (60.3 μL) were added. The reaction mixture was stirred at room temperature under an Ar atmosphere for 1 h, then concentrated under reduced pressure. The residue was purified by flash column chromatography (AcOEt : hexane = 0 : 100 to 100 : 0) to afford compound **5** (89.3 mg, 0.120 mmol) as a colorless solid in 47.9 % yield. The identity of the product was confirmed by means of $^{1}H$ NMR, $^{13}C$ NMR and ESI-HRMS. $^{1}H$ NMR (400 MHz, $CD_2Cl_2$): δ 9.874 (s, 1H), 7. 88 (d, *J* = 8.8 Hz, 2H), 7.22 (d, *J* = 8.8 Hz, 2H), 5.99-5.92 (m, 3H), 5.66 (d, *J* = 8.4 Hz, 2H), 5.41-5.24 (m, 9H), 5.04-4.99 (m, 1H), 4.64-4.63 (m, 4H), 4.59 (d, *J* = 5.6 Hz, 1H), 4.40-4.30 (m, 1H), 4.30-4.20 (m, 2H), 1.92 (s, 3H). $^{13}C$ NMR (101 MHz, $CD_2Cl_2$): δ 192.9, 173.8, 163.2, 156.1, 156.1, 155.5, 133.1, 133.1, 133.0, 133.0, 132.9, 119.4, 119.1, 119.0, 117.9, 98.6, 77.4, 74.0, 72.8, 70.3, 70.0, 69.8, 66.8, 55.7, 23.0. ESI-HRMS (ESI +) m/z calcd. for $[M+Na]^+$, 600.16876; found, 600.16612.

**Scheme 14.** Synthesis of PTX-βGlcNAc.

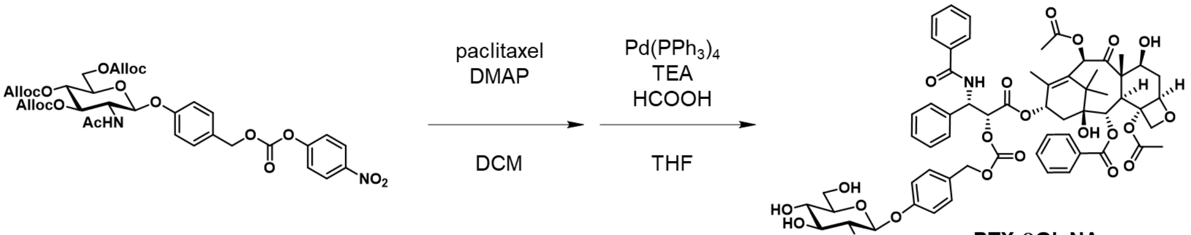


**Procedure for the synthesis of PTX-βGlcNAc.** A mixture of compound 5 (67.2 mg, 0.0900 mmol), paclitaxel (100 mg, 0.117 mmol) and 4-dimethylaminopyridine (DMAP) (14.3 mg, 0.117 mmol) in 5.0 mL dried DCM was stirred at room temperature under an Ar atmosphere for 15 h. After evaporation, the residue was purified by flash column chromatography (AcOEt : hexane = 0 : 100 to 100 : 0) to obtain a white solid, which was dissolved in 8.0 mL dried THF. To this solution, HCOOH (8.1 µL) and triethylamine (45 µL) were added, and the reaction mixture was stirred at room temperature under an Ar atmosphere for 10 min. To this solution, tetrakis(triphenylphosphine)palladium (0) ($Pd(PPh_3)_4$) (24.9 mg, 0.0216 mmol) was added at 0 °C and stirring was continued at room temperature under an Ar atmosphere for 2 h. After evaporation, the residue was purified by flash column chromatography ($CH_2Cl_2$ : MeOH = 100 : 0 to 85 : 15) to afford **PTX-βGlcNAc** (82.0 mg, 0.249 mmol) as a white solid in 75.3 % yield. The identity of the product was confirmed by means of $^1$H NMR, $^{13}$C NMR and ESI-HRMS. $^1$H NMR (400 MHz, MeOD): δ 8.14 (d, *J* = 8.4 Hz, 2H), 7.79 (d, *J* = 8.4 Hz, 2H), 7.68 (t, *J* = 7.6 Hz, 2H), 7.61-7.57 (m, 3H), 7.55-7.50 (m, 3H), 7.47-7.41 (m, 4H), 7.32-7.26 (m, 3H), 7.02 (d, *J* = 8.4 Hz, 2H), 6.48 (s, 1H), 6.10 (t, *J* = 8.8 Hz, 1H), 5.86 (d, *J* = 6.4 Hz, 1H), 5.66 (d, *J* = 7.2 Hz, 1H), 5.50 (d, *J* = 6.4 Hz, 1H), 5.16 (d, *J* = 4.0 Hz, 2H), 5.01 (d, *J* = 9.2 Hz, 2H), 4.39-4.36 (m, 1H), 4.20 (s, 2H), 3.97-3.92 (m, 2H), 3.84 (d, *J* = 6.8 Hz, 1H), 3.75 (dd, *J* = 4.8 Hz, *J* = 12 Hz, 1H), 3.61 (t, *J* = 7.6 Hz, 1H), 3.47-3.46 (m, 2H), 3.37-3.33 (m, 1H), 2.43 (s, 3H), 2.19 (s, 3H), 2.00 (s, 3H), 1.95 (s, 2H), 1.89-1.80 (m, 3H), 1.68 (s, 3H), 1.17 (s, 3H), 1.15 (s, 3H). $^{13}$C NMR (101 MHz, MeOD): δ 205.2, 174.0, 171.7, 171.4, 170.5, 170.3, 167.7, 159.4, 155.8, 142.3, 138.2, 135.4, 135.0, 134.7, 133.0, 131.4, 131.2, 130.5, 130.2, 129.8, 129.6, 128.6, 117.8, 100.7, 85.9, 82.3, 79.1, 78.6, 78.3, 77.5, 76.8, 76.3, 75.8, 73.2, 72.4, 71.9, 71.3, 62.6, 59.3, 57.4, 55.4, 47.9, 44.6, 37.5, 36.5, 27.0, 23.3, 23.1, 22.5, 20.9, 15.0, 10.6. ESI-HRMS (ESI +) m/z calcd. for $[M+Na]^+$, 1229.43124; found, 1229.43200.

**Scheme 15.** Synthesis of compound **6**.

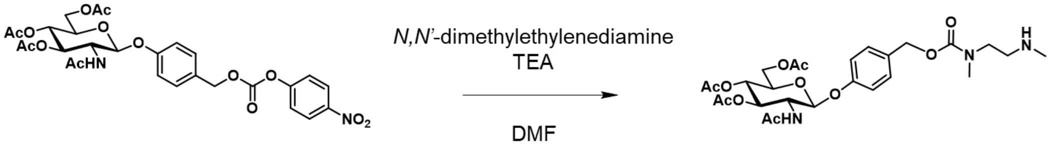


5 6

**Procedure for the synthesis of compound 6.** A mixture of compound **5** (200 mg, 0.324 mmol), *N,N'*-dimethylethylenediamine (114 mg, 1.29 mmol) and triethylamine (2.0 mL) in 5.0 mL DMF was stirred at room temperature under an Ar atmosphere for 2 h. After evaporation, the residue was purified by flash column chromatography ($CH_2Cl_2$ : MeOH = 100 : 0 to 0 : 100) to afford compound **6** (161 mg, 0.284 mmol) as a white solid in 87.7 % yield. The identity of the product was confirmed by means of $^1$H NMR, $^{13}$C NMR and ESI-HRMS. $^1$H NMR (400 MHz, MeOD): δ 7. 35(d, *J* = 8.4 Hz, 2H), 7.04 (d, *J* = 8.8 Hz, 2H), 5.40-5.33 (m, 2H), 5.11-5.07 (m, 3H), 4.34 (dd, *J* = 5.2 Hz, *J* = 12.4 Hz, 1H), 4.18-4.11 (m, 2H), 4.05-4.02 (m, 1H), 3.43 (s, 2H), 2.94 (s, 3H), 2.75-2.70 (m, 2H), 2.41 (s, 2H), 2.35 (s, 2H), 2.06-2.03 (m, 9H), 1.93 (s, 3H). $^{13}$C NMR (101 MHz, MeOD): δ 172.1, 170.8, 170.4, 169.8, 157.0, 131.3, 129.5, 129.2, 116.4, 98.4, 72.4, 71.6, 68.7, 66.5, 61.8, 54.1, 34.5, 33.9, 33.4, 21.3, 19.2, 19.2, 19.2. ESI-HRMS (ESI +) m/z calcd. for $[M+Na]^+$, 590.23203; found, 590.23179.

**Scheme 16.** Synthesis of compound 7.

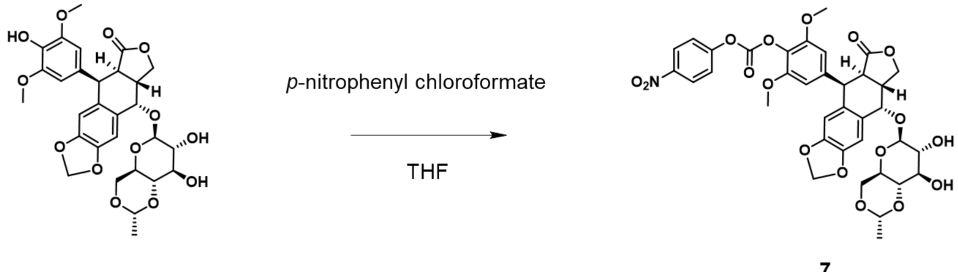


**Etoposide** **7**

**Procedure for the synthesis of compound 7.** A mixture of etoposide (590 mg, 1.00 mmol) and *p*-nitrophenyl chloroformate (200 mg, 1.00 mmol) in 30.0 mL dried THF was stirred at room temperature under an Ar atmosphere for 1 h. After filtration to remove the precipitate, the filtrate was evaporated. The residue was purified by flash column chromatography (AcOEt : hexane = 0 : 100 to 80 : 20) to afford compound **7** (410 mg, 0.544 mmol) as a colorless solid in 54.3 % yield. The identity of the product was confirmed by means of $^1$H NMR, $^{13}$C NMR and ESI-HRMS. $^1$H NMR (400 MHz, $CD_2Cl_2$): δ 8.18 (dd, *J* = 2.0 Hz, *J* = 6.8 Hz, 2H), 7.08 (dd, *J* = 2.4 Hz, *J* = 7.2 Hz, 2H), 6.78 (d, *J* = 10 Hz, 1H), 6.45 (d, *J* = 2.5 Hz, 1H), 6.27 (s, 2H), 5.91 (d, *J* = 8.4 Hz, 2H), 5.24 (s, 1H), 4.84 (d, *J* = 3.2 Hz,

1H), 4.64 (d, J = 5.2 Hz, 1H), 4.57-4.55 (m, 1H), 4.32-4.30 (m, 1H), 4.14 (t, J = 8.4 Hz, 1H), 4.08-4.05 (m, 1H), 3.66 (s, 6H), 3.54-3.44 (m, 3H), 3.28-3.15 (m, 4H), 3.04 (s, 1H), 2.84 (m, 1H), 1.24 (d, J = 5.2 Hz, 1H). $^{13}C$ NMR (101 MHz, $CD_2Cl_2$): δ 175.1, 156.0, 151.6, 150.7, 149.3, 147.8, 146.9, 145.9, 139.5, 132.4, 129.3, 128.3, 125.7, 123.1, 110.8, 109.6, 108.0, 102.6, 102.3, 100.1, 80.3, 74.9, 74.0, 73.5, 68.5, 68.4, 66.8, 56.6, 44.4, 41.3, 38.1, 21.2, 20.5, 14.1. ESI-HRMS (ESI +) m/z calcd. for $[M+Na]^+$, 776.17972; found, 776.17989.

**Scheme 17.** Synthesis of ETP-βGlcNAc.

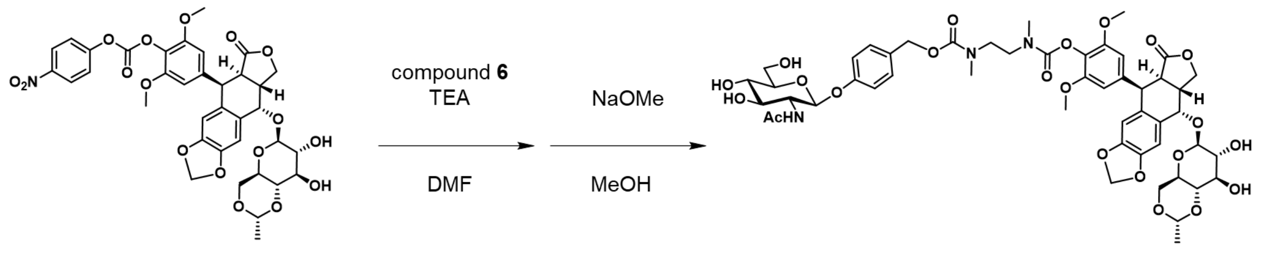


7

ETP-βGlcNAc

**Procedure for the synthesis of ETP-βGlcNAc.** A mixture of compound **7** (214 mg, 0.284 mmol), compound **6** (161 mg, 0.284 mmol) and triethylamine (2.0 mL) in 5.0 mL dried DMF was stirred at room temperature under an Ar atmosphere for 12 h. After evaporation, the residue was purified by flash column chromatography ($CH_2Cl_2$ : MeOH = 100 : 0 to 0 : 100) to obtain a white solid, which was dissolved in 10.0 mL dried MeOH. To this solution, NaOMe (107 mg, 1.98 mmol) was added, and the reaction mixture was stirred at room temperature for 1 h. After quenching with Amberlite IR120 H (300 mg), the solution was filtered and concentrated under reduced pressure. The residue was purified by HPLC to afford the **ETP-βGlcNAc** (192 mg, 0.182 mmol) as a white solid in 64.1 % yield. The identity of the product was confirmed by means of $^1H$ NMR, $^{13}C$ NMR and ESI-HRMS. $^1H$ NMR (400 MHz, MeOD): δ 7.40-7.24 (m, 2H), 7.15 (s, 1H), 7.01-6.91 (m, 2H), 6.61-6.57 (m, 3H), 5.92 (d, J = 6.8 Hz, 2H), 5.372 (m, 1H, overlaps with HOD), 4.84 (s, 1H), 4.73 (d, J = 4.8 Hz, 1H), 4.50-4.47 (m, 1H), 4.39 (s, 1H), 4.35-4.25 (m, 1H), 4.15-4.05 (m, 1H), 3.99-3.89 (m, 4H), 3.76-3.71 (m, 9H), 3.66-3.45 (m, 10H), 3.34-3.20 (m, 5H), 3.11-2.97 (m, 5H), 2.88 (s, 1H), 2.03 (s, 3H), 1.99 (s, 2H), 1.30 (d, J = 5.2 Hz, 1H). $^{13}C$ NMR (101 MHz, MeOD): δ 180.8, 173.1, 159.1, 158.7, 158.0, 157.1, 155.4, 155.2, 153.3, 148.6, 147.2, 140.7, 132.4, 131.3, 131.2, 130.2, 129.9, 129.6, 129.1, 117.4, 116.8, 114.2, 109.4, 107.9, 105.4, 102.1, 101.7, 99.8, 80.6, 77.2, 75.5, 75.0, 73.8, 70.9, 69.2, 68.2, 67.1, 66.5, 61.6, 56.4, 55.9, 55.8, 45.6, 44.1, 39.7, 34.7, 34.5, 22.1, 19.8, 0.0. ESI-HRMS (ESI +) m/z calcd. for $[M+Na]^+$, 1078.36389; found, 1078.36515.

### References


1. Jurrus, E. *et al.* Improvements to the APBS biomolecular solvation software suite. *Protein Science* **27**, 112-128 (2018). https://doi.org/https://doi.org/10.1002/pro.3280
2. BEKKER, H. *et al.* in *4th International Conference on Computational Physics (PC 92).* (ed J Nadrchal RA DeGroot) 252-256 (World Scientific Publishing).
3. Lemkul, J. A. & Bevan, D. R. Assessing the Stability of Alzheimer's Amyloid Protofibrils Using Molecular Dynamics. *The Journal of Physical Chemistry B* **114**, 1652-1660 (2010). https://doi.org/10.1021/jp9110794
4. Gaussian 16 Rev. C.01 (Wallingford, CT, 2016).
5. Kanda, Y. Investigation of the freely available easy-to-use software 'EZR' for medical statistics. *Bone Marrow Transplantation* **48**, 452-458 (2013). https://doi.org/10.1038/bmt.2012.244
6. Kuriki, Y. *et al.* Development of a fluorescent probe library enabling efficient screening of tumour-imaging probes based on discovery of biomarker enzymatic activities. *Chemical Science* **13**, 4474-4481 (2022). https://doi.org/10.1039/D1SC06889J
7. Kawashima, S. *et al.* Rapid imaging of lung cancer using a red fluorescent probe to detect dipeptidyl peptidase 4 and puromycin-sensitive aminopeptidase activities. *Scientific Reports* **12**, 9100 (2022). https://doi.org/10.1038/s41598-022-12665-9
8. Hino, H. *et al.* Rapid Cancer Fluorescence Imaging Using A γ-Glutamyltranspeptidase-Specific Probe For Primary Lung Cancer. *Transl Oncol* **9**, 203-210 (2016). https://doi.org/10.1016/j.tranon.2016.03.007
9. Onoyama, H. *et al.* Rapid and sensitive detection of early esophageal squamous cell carcinoma with fluorescence probe targeting dipeptidylpeptidase IV. *Scientific Reports* **6**, 26399 (2016). https://doi.org/10.1038/srep26399
10. Kitagawa, Y. *et al.* A Novel Topical Fluorescent Probe for Detection of Glioblastoma. *Clinical cancer research : an official journal of the American Association for Cancer Research* **27**, 3936-3947 (2021). https://doi.org/10.1158/1078-0432.ccr-20-4518
11. Takahashi, R. *et al.* Real-Time Fluorescence Imaging to Identify Cholangiocarcinoma in the Extrahepatic Biliary Tree Using an Enzyme-Activatable Probe. *Liver Cancer*, 1-13 (2023). https://doi.org/10.1159/000530645
12. Kobayashi, K. *et al.* Rapid imaging of pulmonary metastasis from colorectal cancer with a red fluorescence probe targeting puromycin-sensitive aminopeptidase and dipeptidyl peptidase IV. *Scientific Reports* **15**, 43930 (2025). https://doi.org/10.1038/s41598-025-27717-z
13. Fujii, T., Kamiya, M. & Urano, Y. In Vivo Imaging of Intraperitoneally Disseminated Tumors in Model Mice by Using Activatable Fluorescent Small-Molecular Probes for Activity of Cathepsins. *Bioconjugate Chemistry* **25**, 1838-1846 (2014). https://doi.org/10.1021/bc5003289
14. Fujita, K. *et al.* Rapid and Accurate Visualization of Breast Tumors with a Fluorescent Probe Targeting α-Mannosidase 2C1. *ACS Central Science* **6**, 2217-2227 (2020). https://doi.org/10.1021/acscentsci.0c01189
15. Ito, H. *et al.* Red-Shifted Fluorogenic Substrate for Detection of lacZ-Positive Cells in Living Tissue with Single-Cell Resolution. *Angewandte Chemie International Edition* **57**, 15702-15706 (2018). https://doi.org/https://doi.org/10.1002/anie.201808670
16. Asanuma, D. *et al.* Sensitive β-galactosidase-targeting fluorescence probe for visualizing small peritoneal metastatic tumours in vivo. *Nature Communications* **6**, 6463 (2015). https://doi.org/10.1038/ncomms7463
17. Urano, Y. *et al.* Rapid Cancer Detection by Topically Spraying a γ-Glutamyltranspeptidase–

Activated Fluorescent Probe. *Science Translational Medicine* **3**, 110ra119-110ra119 (2011). https://doi.org/10.1126/scitranslmed.3002823
18 Sakabe, M. *et al.* Rational Design of Highly Sensitive Fluorescence Probes for Protease and Glycosidase Based on Precisely Controlled Spirocyclization. *Journal of the American Chemical Society* **135**, 409-414 (2013). https://doi.org/10.1021/ja309688m
